# Mapping ortholog-restricted ligandable cysteines in the wheat pathogen *Zymoseptoria tritici*

**DOI:** 10.64898/2026.02.16.706151

**Authors:** Minjin Yoo, Timothy Ware, Sebastián Moschen, Alex Reed, Christopher J. Reinhardt, Evert Njomen, Urvashi Thacker, Bruno Melillo, Gabriel Scalliet, Vlad Pascanu, Benjamin F. Cravatt

## Abstract

*Zymoseptoria (Z.) tritici* is the fungal phytopathogen responsible for Septoria tritici leaf blotch (STB), the main foliar disease of wheat. Fungicides mitigate crop loss caused by STB, but multidrug-resistant *Z. tritici* strains pose a major threat to the global food supply and underscore the need to identify new fungicidal targets. Here, we use activity-based protein profiling to generate a covalent ligandability map of *Z. tritici*, leading to the discovery of cysteines in a diverse set of essential proteins that are targeted by stereochemically defined, electrophilic small molecules (stereoprobes). We identify stereoprobes that display nucleotide-dependent reactivity with an ortholog-restricted cysteine in the GTPase SAR1, leading to accumulation of this protein and other COPII components at apparent ER exit sites and corresponding impairments in *Z. tritici* growth. Our findings thus expand the scope of ligandable, essential proteins in *Z. tritici* and illuminate opportunities to selectively target this destructive plant pathogen.

## Background

Fungal plant pathogens are responsible for widespread crop damage and severe reductions in annual food yield^1^. The fungal ascomycete *Zymoseptoria tritici* (formerly *Mycosphaerella graminicola*) is the causal agent of Septoria tritici blotch (STB), one of the most economically damaging diseases of wheat worldwide. STB is characterized by the appearance of necrotic lesions on wheat leaves that reduce photosynthetic capacity and accelerate premature senescence^2,3^. STB has become a critical concern in global agriculture, particularly in Europe, where it can reduce wheat yields by up to 40%, with average losses often reaching 10-20%^4–6^. Given the importance of wheat as a staple crop, the threat posed by *Z. tritici* directly impacts international food security^5,7^.

Fungicides play a critical role in mitigating global crop loss caused by *Z. tritici*^8^. However, in recent years, multidrug-resistant *Z. tritici* strains have emerged^9–11^, and efforts to counteract this challenge are hindered by the limited number of essential proteins targeted by current armamentarium of fungicides. All major classes of modern fungicides used to control STB are threatened by severe resistance pressure, including sterol biosynthesis inhibitors such as azoles (FRAC group G1)^12^, inhibitors of the respiratory chain Complex II (succinate dehydrogenase) such as pyrazole carboxamides (FRAC group C2)^13^, and strobilurins, which are Complex III (cytochrome b) Qo site inhibitors (FRAC group C3)^14^. The recent introduction of novel Complex III inhibitors targeting the Qi site, such as picoline amides (FRAC group C4), are likely to also experience resistance development like other Complex III agents^15,16^. The identification of additional antifungal targets representing essential, but as-of-yet undrugged *Z. tritici* proteins is thus a high priority. When considering new fungicidal targets, additional concerns arise about the potential for cross-reactivity with animal and plant orthologs. The next generation of “safe by design” fungicides would accordingly benefit from incorporating stringent considerations of species selectivity from the outset.

Covalent chemistry has historically served as a rich source of anti-infective agents to treat both human diseases (e.g., penicillin)^17^ and plant pests (e.g., carbamate and organophosphate pesticides)^18,19^. These compounds, however, react with conserved catalytic serine residues found in the active sites of a limited number of essential hydrolytic enzymes^20,21^. More recently, covalent ligand and drug discovery has been extended to target non-catalytic nucleophilic residues, in particular, cysteines, in a much broader diversity of proteins^22,23^. Covalent ligands offer several potential advantages over more conventional reversibly binding small molecules, including: 1) prolonged pharmacodynamic activity dependent on target turnover rather than compound half-life; 2) improved selectivity by engaging isoform-restricted nucleophilic residues found in one of several structurally related proteins; and 3) the potential to target proteins with shallow or dynamic pockets that were previously considered ‘undruggable’^24,25^. In the agrochemical context, these properties could translate into strong efficacy at lower field application rates and improved selectivity over non-target species.

Covalent ligand and drug discovery has benefited from its integration with the chemical proteomic method activity-based protein profiling (ABPP), which provides global maps of electrophilic small molecule-protein interactions directly in native biological systems^26–28^. The ligandability maps generated by ABPP have served as a rich source of first-in-class chemical tools for structurally and functionally diverse proteins^29–32^ and can also inform on proteome-wide selectivity of more advanced covalent drug candidates^33–35^. Ligand discovery by ABPP has been enhanced in recent years by the design and screening of stereochemically defined electrophilic small molecule (‘stereoprobe’) libraries, in which the stereoselectivity of covalent compound-protein interactions can be used as an identifier of druggable pockets across the proteome^32,36–39^. Covalent stereoprobes also provide a streamlined path for functional studies through the use of chemical (inactive enantiomer) and genetic (stereoprobe-resistant cysteine mutant protein) controls^32,36–39^. So far, however, the integration of stereoprobes with ABPP has mainly been applied to map the covalent ligandability of the human proteome.

Here, we use ABPP in combination with a set of structurally diverse acrylamide stereoprobes to generate a global covalent ligandability map of *Z. tritici*. Integration of these chemical proteomic results with the *Saccharomyces* (*S.*) *cerevisiae* gene deletion database identified structurally and functionally diverse proteins that are both ligandable and essential for fungal growth. Most of these *Z. tritici* proteins have human and wheat orthologs, but, notably, in some instances, the orthologs lack the ligandable cysteine. We found that azetidine acrylamides engage one such ortholog-restricted cysteine (C64) in the *Z. tritici* GTPase COPII component SAR1 in a nucleotide-dependent manner, leading to accumulation of this protein at intracellular membrane sites along the vesicular trafficking pathway and impairments in *Z. tritici* cell growth. These effects were stereoselective and not observed in cells expressing a C64A-SAR1 mutant. Our work thus provides a rich resource of ligandable, essential proteins in the fungal phytopathogen *Z. tritici* and describes an experimental roadmap for testing the functional impact of covalent ligands targeting these proteins as a source of future fungicides for counteracting STB.

## Results

### Initial assessment of stereoprobe reactivity in *Z. tritici*

Previous ABPP studies performed in human cells with diverse classes of stereoprobes have identified acrylamide compounds bearing azetidine and tryptoline cores that stereoselectively engage cysteines on a wide array of structurally and functionally distinct proteins^32,36–38^. We accordingly selected a combination of azetidine and tryptoline acrylamides for generating covalent ligandability maps of *Z. tritici*. We used one set of azetidine acrylamides and three sets of tryptoline acrylamides differing in their appendages (**Fig. 1a**), which allowed to assess the impact of both core (azetidine, tryptoline) and substituent diversification on stereoprobe-protein interactions in *Z. tritici*.

**Figure 1.**
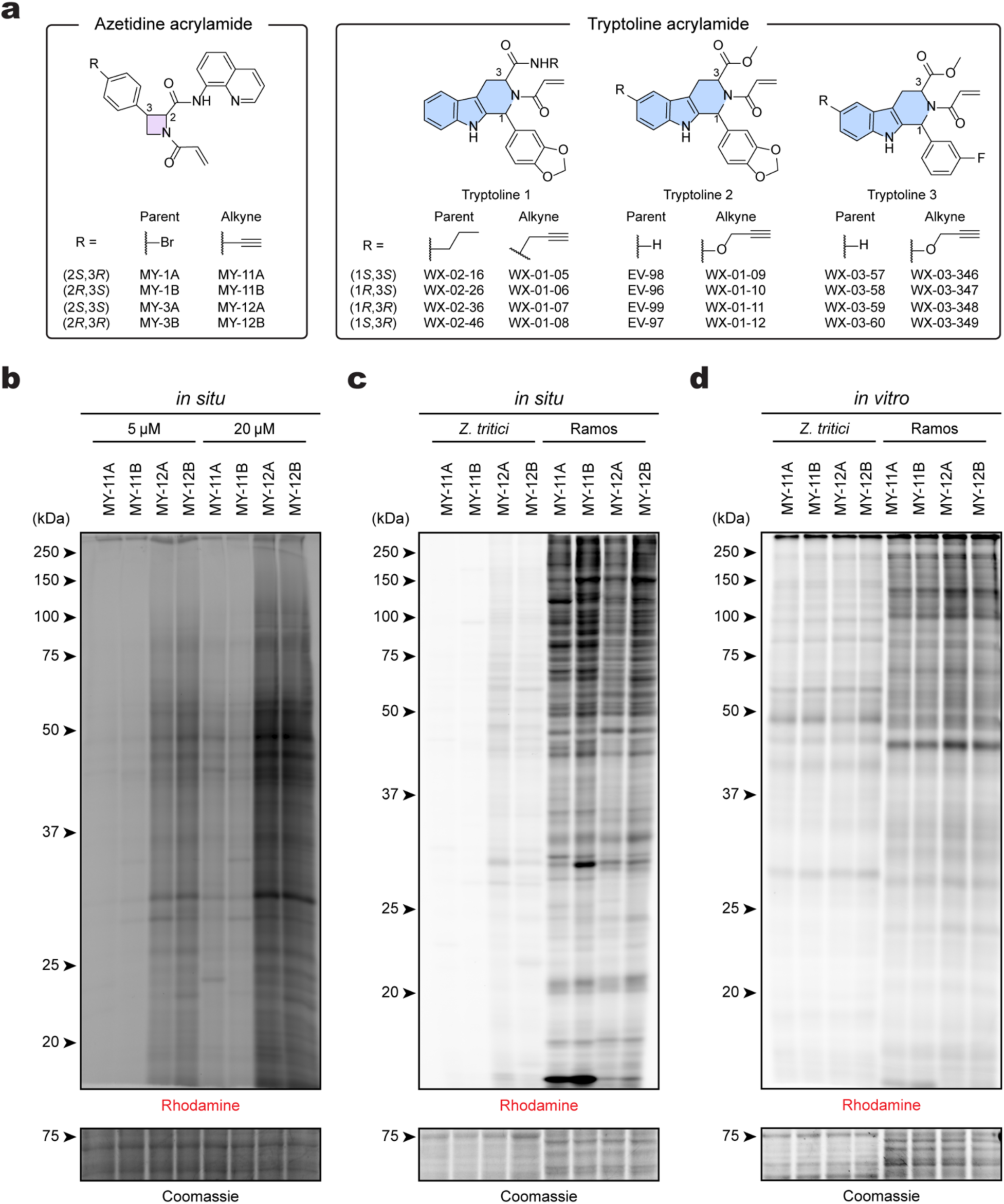
Gel-ABPP analysis of the proteome reactivity of acrylamide stereoprobes in *Z. tritici* cells. **a**, Structures of parent and alkynylated azetidine and tryptoline acrylamide stereoprobes used in this study. **b,** Gel-ABPP data showing the concentration-dependent *in situ* reactivity of alkynylated azetidine acrylamide stereoprobes (5 µM and 20 µM, 3 h) in *Z. tritici* as determined by gel-ABPP. Stereoprobe-reactive proteins are visualized by CuAAC conjugation to an azide-rhodamine reporter group, SDS-PAGE and in-gel fluorescence scanning. **c, d,** Gel-ABPP data showing (**c**) *in situ* or (**d**) *in vitro* reactivity of alkyne azetidine acrylamide stereoprobes (5 µM, 3 h) in *Z. tritici* and Ramos cells. For **b-d**, data are from a single experiment representative of two independent experiments.

We first investigated the general stereoprobe reactivity of liquid cultures of *Z. tritici* (IPO323 strain) by gel-ABPP. In these experiments, *Z. tritici* cells were treated with alkyne analogs of the stereoprobes (5 or 20 µM, 3 h) (**Fig. 1a**), lysed, and stereoprobe-modified proteins were conjugated to an azide-rhodamine tag by copper-catalyzed azide-alkyne cycloaddition (CuAAC, or click)^40,41^ chemistry and analyzed by SDS-PAGE and in-gel fluorescence scanning^42^. Most of the stereoprobes showed concentration-dependent increases in protein reactivity in *Z. tritici* cells, with the exception of one set of tryptoline acrylamides (WX-01-05 – WX-01-08), which showed negligible protein reactivity at either test concentration (**Extended Data Fig. 1a**). We also observed striking differences among the azetidine acrylamides, with the *trans* compounds ((2*S*, 3*S*); (2*R*, 3*R*)) displaying much greater protein reactivity than the *cis* compounds ((2*S*, 3*R*); (2*R*, 3*S*)) (**Fig. 1b**). Consistent with previous work^32,43^, these differences in proteomic reactivity were not observed for the azetidine and tryptoline acrylamides in human cells, which instead displayed more consistent reactivity profiles that matched the reported intrinsic electrophilicities of the stereoprobes as determined by reactivity with glutathione^32,43^ (**Fig. 1c** and **Extended Data Fig. 1b**). The azetidine and tryptoline stereoprobes also showed much greater overall protein reactivity in human cells compared to *Z. tritici* cells (**Fig. 1c** and **Extended Data Fig. 1b**). We interpreted these gel-ABPP data to indicate that stereoprobe uptake and/or metabolism differed between human cells and *Z. tritici* cells, leading to lower *in situ* reactivity in the latter cells. Also supportive of this hypothesis, more consistent and similar *in vitro* reactivity profiles were observed for stereoprobes in lysates of *Z. tritici* and human cells (**Fig. 1d** and **Extended Data Fig. 2a, b**).

Based on our initial gel-ABPP findings, we concluded that most of the azetidine and tryptoline acrylamide stereoprobes were capable of engaging proteins in *Z. tritici* cells, although with lower efficiency than in human cells. We therefore elected to generate both live-cell (*in situ*) and lysate (*in vitro*) covalent ligandability maps of *Z. tritici*, with the former providing a more physiologically relevant context for evaluating stereoprobe-protein interactions and the latter allowing for the testing of higher effective concentrations of the stereoprobes.

### Covalent stereoprobe ligandability maps of *Z. tritici*

We generated initial covalent stereoprobe ligandability maps of *Z. tritici* cells and cell lysates by protein-directed ABPP coupled with multiplexed quantitative tandem mass tagging (TMT^16plex^) mass spectrometry (MS)-based proteomics^32^. In brief, *Z. tritici* cells or cell lysates were treated with parent stereoprobes (40 µM) or DMSO for 2 h, followed by treatment with the corresponding alkyne stereoprobes (10 µM) for 1 h. Alkyne stereoprobe-modified proteins were then conjugated to an azide-biotin tag by CuAAC, processed and analyzed by MS-based proteomics (**Fig. 2a**). Quantified proteins were assigned as liganded if they fulfilled the following criteria: (1) greater than 3-fold enantioselective enrichment with at least one pair of alkyne stereoprobe enantiomers; and (2) >50% blockade of this enrichment by the corresponding parent stereoprobe. The stereoprobe ligandability maps were then displayed in quadrant plots (**Fig. 2b-d** and **Extended Data Fig. 3a-e**), where the positions of proteins on the *x*- and *y*-axes indicate enantioselective and diastereoselective enrichment, respectively, and the size of each dot represents the degree of blockade of this enrichment by the parent stereoprobe. The ligandability maps qualitatively matched the gel-ABPP data for each set of stereoprobes, as exemplified by comparison of the *in situ* versus *in vitro* maps for the azetidine acrylamides. The *trans* azetidine acrylamides (parent: MY-3A/3B; alkyne: MY-12A/12B) liganded many more proteins than their *cis* counterparts (parent: MY-1A/1B; alkyne: MY-11A/11B) *in situ* (**Fig. 2b**), while a more equivalent number of proteins was liganded by the enantiomeric pairs of azetidine acrylamides in *Z. tritici* lysates (**Fig. 2c**).

**Figure 2.**
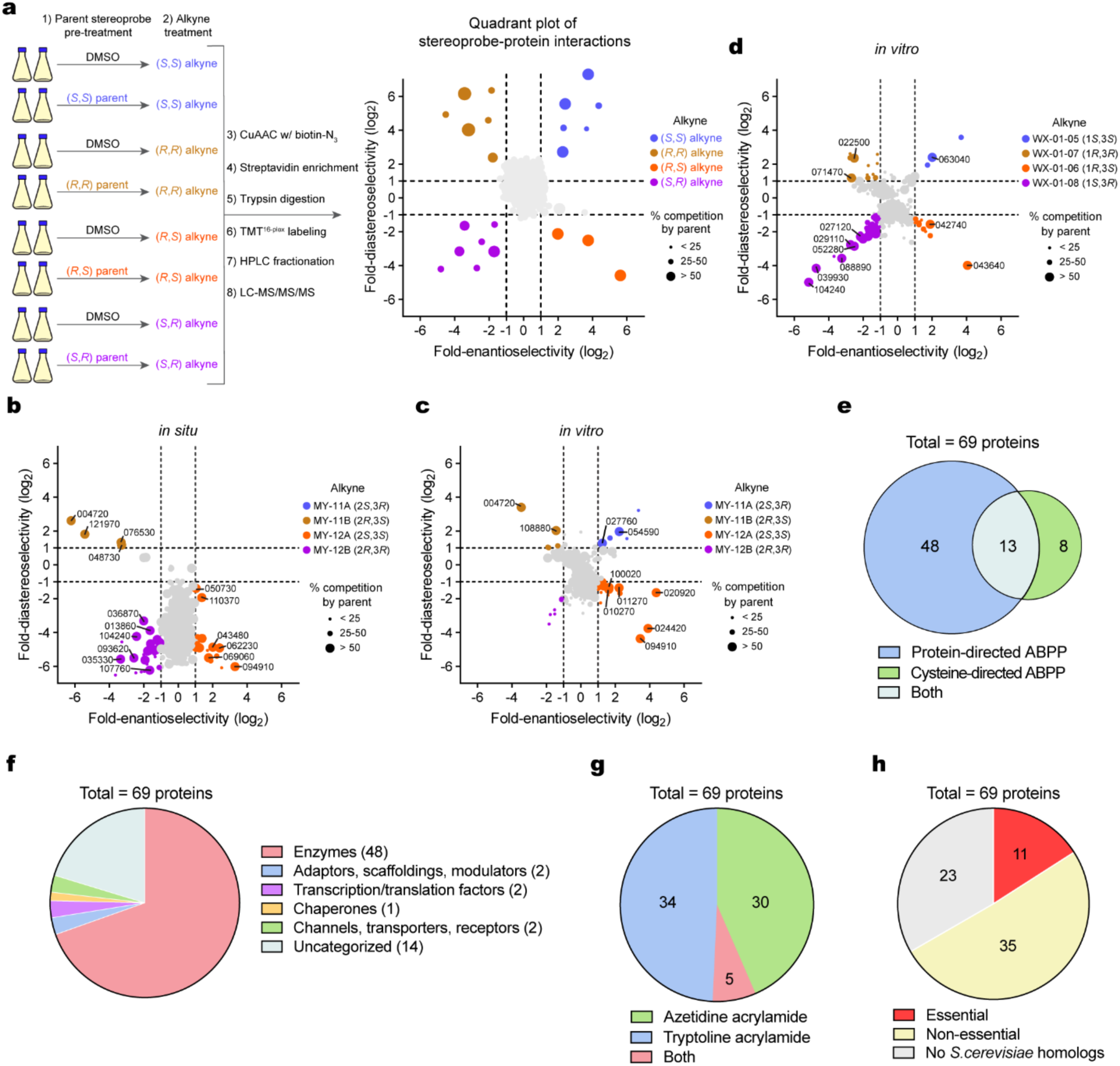
Mapping stereoprobe-liganded proteins and cysteines in *Z. tritici* by ABPP. **a**, Workflow of protein-directed ABPP and mock-up quadrant plot for data visualization. **b-d,** Protein-directed ABPP data shown as quadrant plots for azetidine (parent: MY-1A/1B/3A/3B (40 µM, 2 h); alkyne: MY-11A/11B/12A/12B (10 µM, 1 h)) (**b,** *in situ*; **c,** *in vitro*) and tryptoline (parent: WX-02-16/26/36/46 (40 µM, 2 h); alkyne: WX-01-05/06/07/08 (10 µM, 1 h)) (**d,** *in vitro*) acrylamide stereoprobes in *Z. tritici* cells (**b**) or lysates (**c, d**). Representative stereoprobe-liganded proteins are denoted in each quadrant plot by their gene IDs (i.e., ZtIPO323_XXXXXX). Data are average values from four independent experiments. **e,** Venn diagram showing fraction of stereoselectively liganded proteins mapped by cysteine- and/or protein-directed ABPP. **f,** Pie chart showing protein class distribution for stereoprobe-liganded proteins in *Z. tritici*. **g,** Pie chart showing fraction of proteins liganded by azetidine and/or tryptoline acrylamide stereoprobes. **h,** Pie chart showing fraction of stereoprobe-liganded proteins in *Z. tritici* that are essential in *S. cerevisiae*.

To supplement the protein-directed ABPP data and identify specific cysteines liganded by stereoprobes in *Z. tritici* proteins, we performed cysteine-directed ABPP^44^ following established protocols^45^. Each multiplexed (TMT^10plex^) experiment compared the four stereoisomers of a given stereoprobe set (40 µM each) alongside a DMSO control for blockade of cysteine reactivity with the broadly reactive agent iodoacetamide-desthiobiotin (IA-DTB). From an average of 6,910 quantified cysteines per stereoprobe set, we identified liganded cysteines in 13 of the 61 stereoprobe targets assigned by protein-directed ABPP, where liganding of a cysteine was defined as a >66.7% decrease in IA-DTB reactivity that was also at least 2.5-fold greater than that observed for the enantiomeric stereoprobe (**Fig. 2e** and **Supplementary Dataset 1**). The cysteine-directed ABPP experiments also identified an additional 8 stereoprobe-liganded cysteines in proteins that were not assigned as stereoprobe targets by protein-directed ABPP experiments (**Fig. 2e** and **Supplementary Dataset 1**). As described previously^32^, cysteine- and protein-directed ABPP have complementary attributes and limitations for assigning covalent small molecule-protein interactions, and a more complete covalent ligandability map can accordingly be generated by using both methods.

The 69 in total stereoselectively liganded *Z. tritici* proteins were enriched for enzymes but also included other protein classes that have historically proven more challenging to address with small molecules (e.g., adaptors, scaffolding proteins, transcription/translation factors) (**Fig. 2f**). As has been observed in human cells^32,37,38^, the azetidine and tryptoline acrylamides largely liganded distinct sets of proteins — 30 and 34 proteins, respectively (**Fig. 2g**) — emphasizing the importance of core diversity for increasing the breadth of stereoprobe interactions with the proteome. While the tryptoline stereoprobe ligandability map was mostly derived from *Z. tritici* lysates (*in vitro*), with only a handful of liganding events observed in cells (*in situ*), the azetidine acrylamides liganded a number of proteins *in situ* (**Fig. 2b** and **Supplementary Dataset 1**). Several of the azetidine acrylamide targets showed similar liganding profiles *in vitro* and *in situ*, but there were exceptional cases of proteins that were only liganded *in situ* (e.g., *Zt*NSDHL; ZtIPO323_018650; **Extended Data Fig. 3f**), pointing to the potential impact of the cellular environment of *Z. tritici* on modulating stereoprobe-protein interactions (see below). Finally, we cross-referenced the genes encoding stereoprobe-liganded *Z. tritici* proteins to the *Saccharomyces* Genome Database, which revealed that ∼16% of these genes have orthologs in *S. cerevisiae* that are essential (**Fig. 2h**).

### Characterization of stereoprobe-liganded *Z. tritici* proteins

A substantial fraction of the stereoprobe-liganded proteins in *Z. tritici* also have human orthologs or homologs based on NCBI BLAST searches (*E*-value cutoff <1e^-4^) (**Fig. 3a** and **Supplementary Dataset 1**). For these homologous protein pairs, we found only a few instances in which the stereoprobe liganding profiles of *Z. tritici* proteins were shared by their human counterparts (as determined by cross-referencing previously reported cysteine- and protein-directed ABPP data performed with azetidine/tryptoline acrylamides in human cells^32,37,38^) (**Fig. 3a**). This outcome was unchanged if we restricted the comparison to *in situ* liganding events from *Z. tritici* to better match previous human stereoprobe ligandability data, which were mainly generated in cells (**Extended Data Fig. 4a**). In rare cases where the stereoprobes liganded both *Z. tritici* and human orthologs of a protein, such as the methyltransferase NSUN2 (ZtIPO323_004720), these liganding events occurred at the same cysteine and exhibited the same core (azetidine acrylamide) and stereo-configuration (MY-1B) preferences (**Fig. 3b-d** and **Extended Data Fig. 4b, c**).

**Figure 3.**
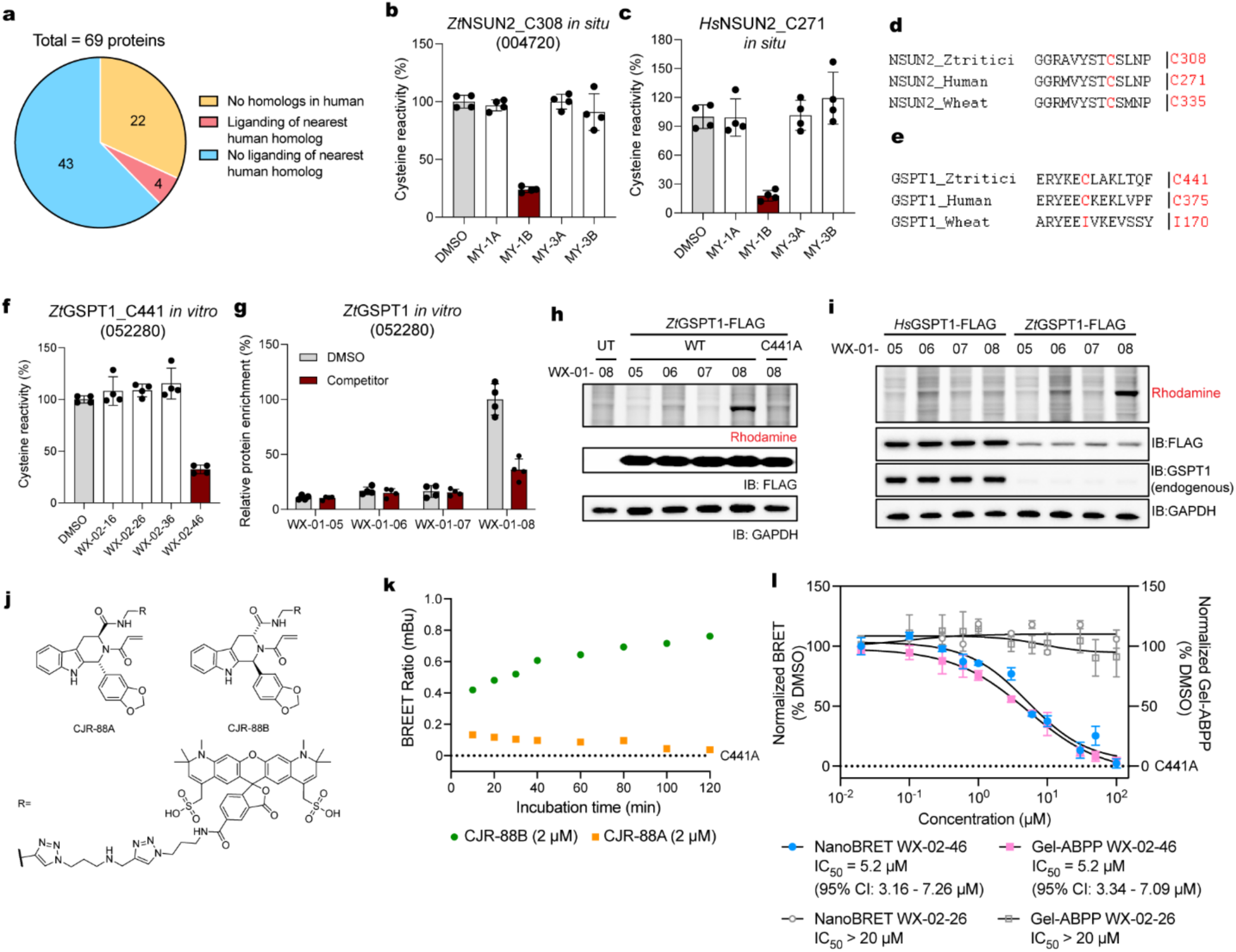
Characterization of stereoprobe-liganded essential proteins from *Z. tritici*. **a**, Pie chart showing fraction of stereoprobe-liganded cysteines/proteins from *Z. tritici* for which the nearest human homolog is also liganded. **b,** Cysteine-directed ABPP data showing stereoselective liganding of *Zt*NSUN2_C308 (ZtIPO323_004720) by MY-1B (40 µM, 2 h). Data are from *in situ* treatments of *Z. tritici*. **c,** Cysteine-directed ABPP data showing stereoselective liganding of *Hs*NSUN2_C271 by MY-1B (20 µM, 3 h). Data are from *in situ* treatments of Ramos cells. For **b**, **c**, data represent mean values ± s.d. of four independent experiments. **d,** Sequence alignment of NSUN2 orthologs showing conservation of the liganded cysteine (red) in *Z. tritici*, human, and wheat. **e,** Sequence alignment of GSPT1 orthologs showing conservation of the liganded cysteine (red) in *Z. tritici* and human, but not wheat. **f,** Cysteine-directed ABPP data showing stereoselective liganding of *Zt*GSPT1_C441 (ZtIPO323_052280) by WX-02-46 (40 µM, 2 h). Data are from *in vitro* treatments of *Z. tritici* lysates. **g,** Protein-directed ABPP data showing stereoselective liganding of *Zt*GSPT1 by the alkyne/parent stereoprobe pair WX-01-08/WX-02-46 (10 µM, 1 h; 40 µM, 2 h pre-treatment). Data are from *in vitro* treatments of *Z. tritici* lysates. For **f**, **g**, data represent mean values ± s.d. of four independent experiments. **h,** Gel-ABPP data showing stereoselective engagement of recombinant WT-*Zt*GSPT1, but not C441A-*Zt*GSPT1 by WX-01-08 (*in situ,* 2 µM, 1 h) in HEK293T cells. **i,** Gel-ABPP data showing stereoselective engagement of recombinant *Zt*GSPT1, but not *Hs*GSPT1 by WX-01-08 (5 µM, 1 h) in HEK293T cells. For **h**, **i**, data are from a single experiment representative of two independent experiments. **j,** Structures of NanoBRET-ABPP stereoprobe tracers CJR-88A and CJR-88B. **k,** NanoBRET-ABPP data showing time-dependent stereoselective reactivity of WT-*Zt*GSPT1-NLuc with CJR-88B (2 µM). **l,** Comparison of gel- and NanoBRET-ABPP data showing concentration-dependent, stereoselective blockade of CJR-88B (2 µM, 2 h) reactivity with WT-*Zt*GSPT1-NLuc by WX-02-46 (0.02-50 µM, 2 h). **k**, **l**, Experiments were performed in lysates from HEK293T cells expressing WT-*Zt*GSPT1-NLuc (NanoBRET-ABPP) or ZtGSPT1-FLAG proteins (gel-ABPP). Dashed horizontal lines mark the background signals for NanoBRET-ABPP assays performed with C441A-*Zt*GSPT1-NLuc. Data are the average values ± s.e.m. from three biological replicates.

While we anticipated that divergent stereoprobe liganding profiles for *Z. tritici* versus human orthologs may often occur at cysteines that are unique to the *Z. tritici* proteins (see below), we also observed instances of exclusive liganding of *Z. tritici* proteins at conserved cysteines (**Supplementary Dataset 1**). One example was the essential translation termination factor GSPT1, for which the conserved cysteine (C441 in *Z. tritici* GSPT1 (*Zt*GSPT1; ZtIPO323_052280); C375 in human GSPT1 (*Hs*GSPT1)) was stereoselectively liganded in *Zt*GSPT1 by tryptoline acrylamide WX-02-46 (**Fig. 3e-g**). *Zt*GSPT1 and *Hs*GSPT1 share ∼55% sequence identity, but our previous ABPP studies^32^ did not show evidence of liganding of *Hs*GPST1 by WX-02-46 (**Extended Data Fig. 4d**). To understand if this apparent difference in stereoprobe interactions was an intrinsic feature of the *Zt*GSPT1 versus *Hs*GSPT1 proteins, we stably expressed each protein in HEK293T cells. Exposure of these cells to alkyne tryptoline acrylamides WX-01-05/06/07/08 followed by gel-ABPP revealed stereoselective engagement of *Zt*GSPT1 by WX-01-08 (**Fig. 3h**), matching the stereoselective enrichment of endogenous *Zt*GSPT1 determined by protein-directed ABPP (**Fig. 3g**). We also found that pre-treatment with WX-02-46, but not WX-02-26, blocked WX-01-08-*Zt*GSPT1 interactions (**Extended Data Fig. 4e**). In contrast, neither the *Hs*GSPT1 protein (**Fig. 3i**) nor a C441A mutant of *Zt*GSPT1 (**Fig. 3h**) reacted with WX-01-08. We interpret these data to indicate that the preferential stereoprobe liganding of *Zt*GSPT1 is an intrinsic feature of this protein that is not shared by *Hs*GSPT1. While we do not yet understand the molecular basis for the differential ligandability of *Hs*GSPT1 and *Zt*GSPT1 proteins, structural models indicate at least five non-conserved amino acids are within 5 Å of the liganded cysteine – Y394, V423, K439, L442, and A443 in *Zt*GSPT1 (**Extended Data Fig. 4f, g**).

The liganding of *Zt*GSPT1 by WX-02-46 was observed in both *Z. tritici* lysates (**Fig. 3g**) and HEK293T cells (**Extended Data Fig. 4e**), but not in *Z. tritici* cells (**Extended Data Fig. 4h**). These data are consistent with the lack of engagement of *Zt*GSPT1 in *Z. tritici* cells being due to poor cellular uptake and/or rapid metabolism of tryptoline acrylamides. Toward the longer term goal of identifying alternative chemotypes for targeting *Zt*GSPT1, we leveraged the tryptoline acrylamide ligands to establish a high-throughput screening-compatible NanoBRET-ABPP assay^46–48^. In this assay, lysates of HEK293T cells expressing a nanoluciferase GSPT1 fusion protein (*Zt*GSPT1-NLuc) in HEK293T cells were exposed to a fluorescently tagged analog of WX-02-46 (CJR-88B) or an enantiomeric control compound (CJR-88A) (**Fig. 3j**), and bioluminescence energy transfer was then measured from the NLuc fusion protein donor to the fluorescent tracer acceptor. We first confirmed by gel-ABPP that CJR-88B stereoselectively engaged WT-*Zt*GSPT1-NLuc, but not C441A-*Zt*GSPT1-NLuc mutant (**Extended Data Fig. 4i**) and this interaction was stereoselectively blocked in a concentration-dependent manner by pretreatment with WX-02-46 (**Extended Data Fig. 4j, k**). Similar results were obtained by NanoBRET-ABPP, where we observed time-dependent increases in BRET signals for CJR-88B-treated, but not CJR-88A-treated WT-*Zt*GSPT1-NLuc (**Fig. 3k**), and the CJR-88B-WT-ZtGSPT1-NLuc signals were stereoselectively blocked by pre-treatment with WX-02-46 (**Fig. 3l**). A similar IC_50_ value of ∼5 µM was measured for WX-02-46 by both gel- and NanoBRET-ABPP (**Fig. 3l**). We anticipate that transitioning from lower throughput MS- and gel-ABPP experiments to NanoBRET-ABPP assays should facilitate future efforts to screen larger compound libraries against prioritized protein targets from the stereoprobe ligandability map of *Z. tritici*.

### Essential *Z. tritici* proteins with ortholog-restricted ligandable cysteines

From our *Z. tritici* ligandability map, we were particularly interested in essential proteins possessing liganded cysteines that were not conserved in orthologous human and wheat (*Triticum aestivum*) proteins (**Supplementary Dataset 1**), as such sites could provide a way to selectively perturb the functions of *Z. tritici* proteins. One example was the *N*-myristoyltransferase enzyme *Zt*NMT1 (ZtIPO323_063250), which exhibited stereoselective liganding by the alkyne/parent tryptoline acrylamide pair WX-03-349/WX-03-60 (**Fig. 4a, b**). The liganded cysteine residue in *Zt*NMT1 — C141 — is not found in either of the two human orthologs NMT1 and NMT2 nor the wheat NMT ortholog (**Fig. 4c**) and is predicted by AlphaFold structural modeling to be located proximal to the myristoyl-CoA cofactor binding pocket (**Fig. 4d**). NMT1 is an essential enzyme responsible for incorporating N-terminal myristic acid modifications into proteins and has been studied as an agrochemical target in *Z. tritici*^49^ as well as a drug target for several human pathogens, including *Plasmodium falciparum*^50^*, Leishmania donovani*^51^, and *Trypanosoma brucei*^52^. Previous inhibitors of *Z. tritici* NMT1 reversibly bind to the peptide substrate pocket (**Fig. 4d**) and generally show strong cross-reactivity with human NMT1^49^. Our findings point to a potential alternative way to generate ortholog-restricted ligands for *Z. tritici* NMT1 by covalent chemistry.

**Figure 4.**
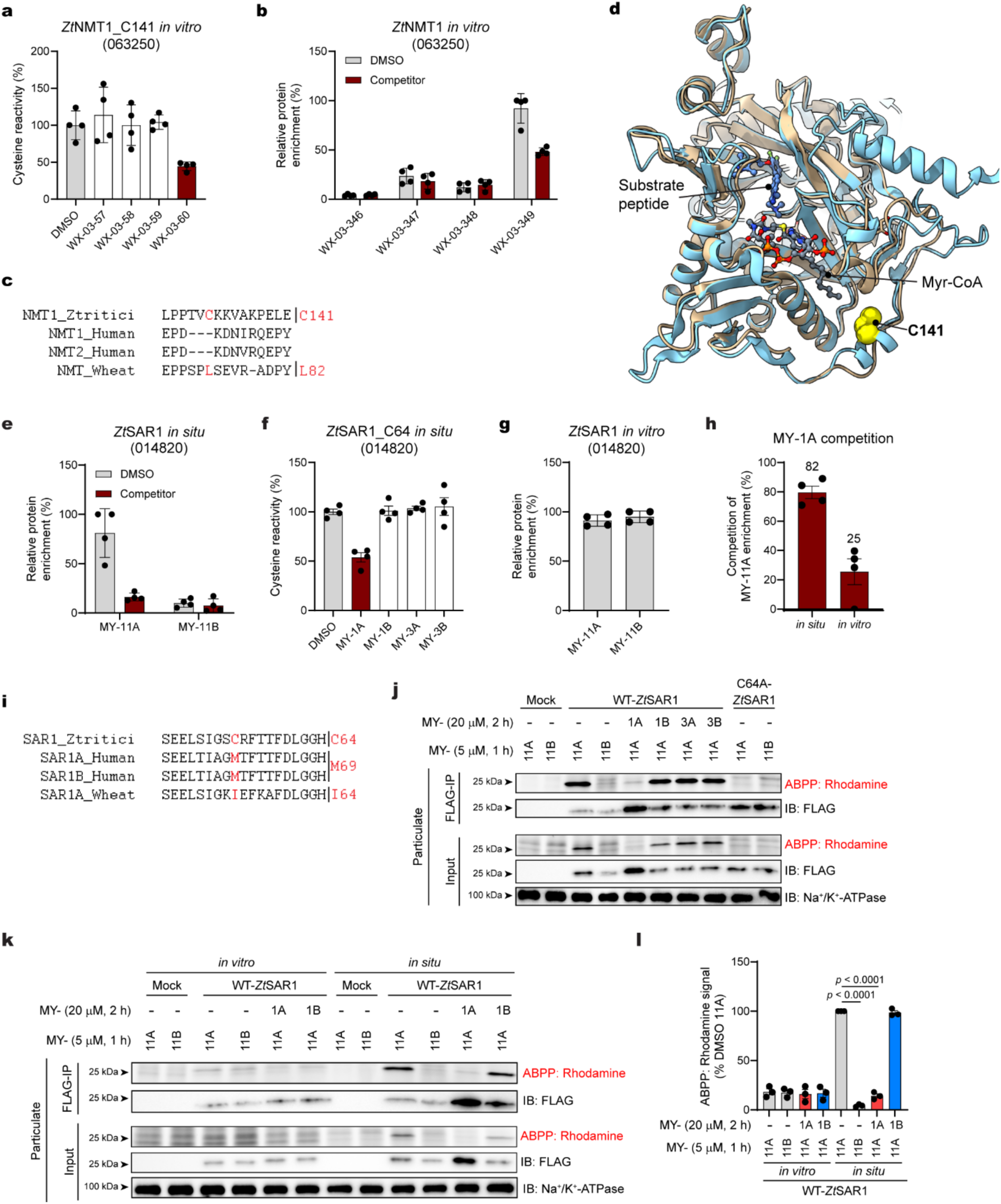
Characterization of ortholog-restricted, stereoprobe-liganded cysteines in *Z. tritici.* **a**, Cysteine-directed ABPP data showing stereoselective liganding of C141 of *Zt*NMT1 (ZtIPO323_063250) by WX-03-60 (40 µM, 2 h) in *Z. tritici* cell lysates. **b,** Protein-directed ABPP data showing stereoselective liganding of *Zt*NMT1 by the alkyne/parent stereoprobe pair WX-03-349/WX-03-60 (10 µM, 1 h; 40 µM, 2 h pre-treatment) in *Z. tritici* cell lysates. For **a**, **b**, data represent mean values ± s.d. of four independent experiments. **c,** Sequence alignment of NMT1 orthologs showing lack of conservation of stereoprobe-liganded C141 of *Zt*NMT1 in human and wheat orthologs. **d,** Overlay of a predicted AlphaFold structure of *Zt*NMT1 (AF-F9XBB7-F1) and the crystal structure of *Hs*NMT1 in complex with a substrate peptide and myristoyl-CoA (Myr-CoA) cofactor (PDB: 5MU6). The location of C141 in *Zt*NMT1 is shown yellow. **e, f,** *In situ* protein-directed (**e**) and cysteine-directed (**f**) ABPP data showing stereoselective liganding of SAR1 (ZtIPO323_014820) by the alkyne/parent stereoprobe pair MY-11A/MY-1A (10 µM, 1 h; 40 µM, 2 h pre-treatment). Data represent mean values ± s.d. of four independent experiments. **g,** *In vitro* protein-directed ABPP data showing a lack of stereoselective enrichment of SAR1 in *Z. tritici* cell lysates by MY-11A (10 µM, 1 h). Data represent mean values ± s.d. of four independent experiments. **h,** Bar graph showing the extent of competitive blockade of MY-11A (10 µM, 1 h) enrichment of *Zt*SAR1 by pre-treatment with MY-1A (40 µM, 2 h) from protein-directed ABPP experiments performed *in vitro* or *in situ.* Data represent mean values ± s.d. of four independent experiments. **i,** Sequence alignment of SAR1 orthologs showing lack of conservation of stereoprobe-liganded C64 of *Zt*SAR1 in human and wheat orthologs. **j,** Gel-ABPP data showing engagement of recombinant WT-*Zt*SAR1, but a C64A-*Zt*SAR1 mutant by MY-11A (5 µM, 1 h) in HEK293T cells, as well as stereoselective blockade of MY-11A engagement of WT-*Zt*SAR1 by pre-treatment with MY-1A (20 µM, 2 h). Data are from a single experiment representative of three independent experiments. **k,** Gel-ABPP data showing engagement of recombinant WT-*Zt*SAR1 by MY-11A (5 µM, 1 h) in HEK293T cells, as well as stereoselective blockade of this engagement by pre-treatment with MY-1A (20 µM, 2 h) in HEK293T cells (*in situ*), but not cell lysates (*in vitro*). **l,** Quantification of gel-ABPP data shown in (**k**). Data represent mean values ± s.e.m. for three independent experiments; *p* values from unpaired two-tailed t tests.

Our protein-directed ABPP data identified another essential protein – the small COPII coat GTPase SAR1 (*Zt*SAR1; ZtIPO323_014820) — that was stereoselectively liganded by the alkyne/parent azetidine acrylamide pair MY-11A/MY-1A (**Fig. 4e**). We also found evidence of stereoselective liganding of C64 in *Zt*SAR1 by MY-1A that fell just below our threshold of 50% blockade of IA-DTB reactivity in cysteine-directed ABPP experiments (47% blockade; **Fig. 4f**). Of note, the liganding of *Zt*SAR1 by azetidine acrylamides was only observed in *Z. tritici* cells and not in *Z. tritici* lysates, as assessed by either stereoselective enrichment by alkyne MY-11A (compare Fig. 4e and **g**) or blockade of this enrichment by MY-1A (**Fig. 4h**).

C64 is unique to *Zt*SAR1 in comparison to the two human orthologs SAR1A and SAR1B and the wheat ortholog SAR1A (**Fig. 4i**), with the proteins otherwise sharing high sequence identity (**Extended Data Fig. 5a**). A review of our protein-directed ABPP data of human cells also found no evidence for stereoselective engagement of *Hs*SAR1A by MY-11A/MY-1A (**Extended Data Fig. 5b**), supporting that azetidine acrylamides act as ortholog-restricted ligands for *Zt*SAR1. We found by gel-ABPP that recombinant WT-*Zt*SAR1, but not a C64A-*Zt*SAR1 mutant, was stereoselectively engaged by MY-11A in HEK293T cells, and this interaction was stereoselectively blocked by pre-treatment with MY-1A (**Fig. 4j** and **Extended Data Fig. 5c, d**). Consistent with the behavior of endogenous *Zt*SAR1, the recombinant protein only reacted with MY-11A in HEK293T cells and not in cell lysates (**Fig. 4k, l**).

SAR1 is responsible for initiating anterograde vesicular trafficking through the recruitment of coat protein complex II (COPII) proteins to endoplasmic reticulum (ER) exit sites^53,54^. In this role, SAR1, upon binding GTP, undergoes a conformational change exposing an amphipathic helix that binds to the ER membrane^54–57^. Interestingly, we consistently observed that recombinant WT-*Zt*SAR1 protein accumulated in the particulate fraction of HEK293T cells treated with MY-1A (**Fig. 4j, k**, **Fig. 5a-c** and **Extended Data Fig. 5c, e**). This effect was both stereoselective (not observed with MY-1B; **Fig. 4j, k**, **Fig. 5a-c** and **Extended Data Fig. 5c, e**) and concentration-dependent (**Extended Data Fig. 6a, b**), displaying an EC_50_ value (11.1 µM) that matched the IC_50_ value measured for engagement of WT-*Zt*SAR1 as determined by gel-ABPP (11.7 µM; **Extended Data Fig. 6a, b**). The MY-1A-induced increase of WT-*Zt*SAR1 in the particulate fraction of cells occurred within 30 min of treatment and without substantial alterations in the expression level of the protein (**Extended Data Fig. 6c-e**).

**Figure 5.**
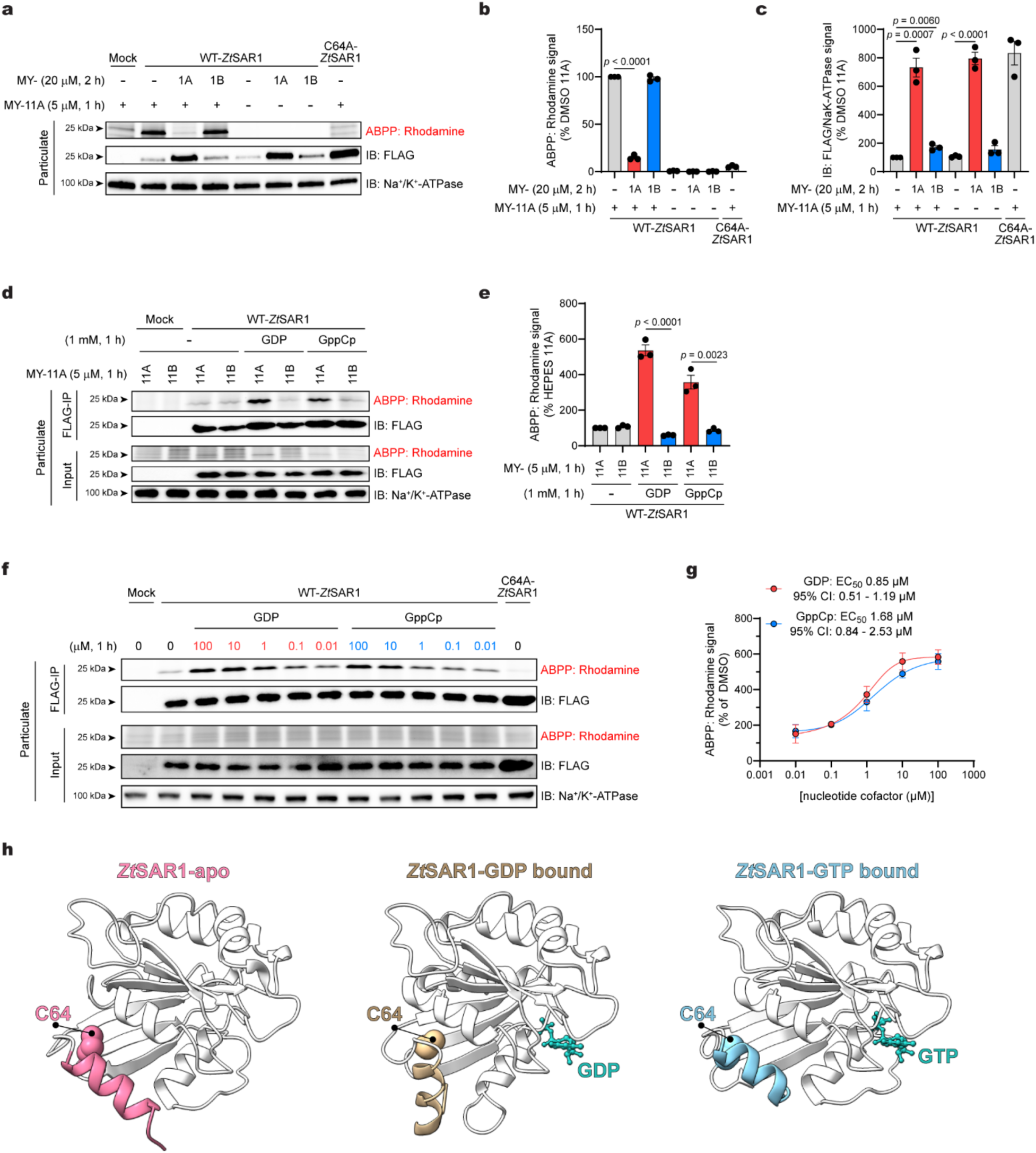
Azetidine acrylamide stereoprobes promote *Zt*SAR1 accumulation in the particulate fraction of cells. **a**, Gel-ABPP data showing stereoselective blockade of MY-11A (5 µM, 1 h) engagement of WT-*Zt*SAR1 by pre-treatment with MY-1A (20 µM, 2 h) in HEK293T cells and concomitant accumulation of WT-*Zt*SAR1 in the particulate fraction. **b,** Quantification of gel-ABPP data shown in (**a**). **c,** Quantification of ZtSAR1 immunoblotting signals shown in (**a**). **d**, Gel-ABPP data showing the effect of GDP/GppCp (1 mM, 1 h) on the stereoselective engagement of WT-*Zt*SAR1 by MY-11A (5 µM, 1 h) in HEK293T cell lysates. **e,** Quantification of the gel-ABPP data shown in (**d**). For **b**, **c**, and **e**, data represent mean values ± s.e.m. for three independent experiments; *p* values from unpaired two-tailed t tests. **f,** Gel-ABPP data showing the concentration-dependent effects of GDP/GppCp on MY-11A (5 µM, 1 h) engagement of WT-*Zt*SAR1. **g,** Quantification of gel-ABPP data shown in (**f**). For **g**, data represent mean values ± s.d. of three independent experiments. **h,** Boltz-2-predicted conformational changes in predicted structures of *Zt*SAR1 in the apo form (pink, AF-F9X159-F1) or bound to GDP or GTP (brown and teal respectively, Boltz-2 predictions). Structures highlight that the positioning of C64 of *Zt*SAR1 in relation to the N-terminal helix (residues 1-23) is predicted to be affected by binding of GNP cofactor.

The aforementioned results suggested that MY-1A binding to WT-*Zt*SAR1 leads to increased association of this protein with the membrane compartment of cells. Considering further that GTP binding also promotes WT-*Zt*SAR1 translocation to cell membranes^54–57^, and we did not observe the binding of MY-1A to WT-*Zt*SAR1 in HEK293T cell lysates, we wondered whether azetidine acrylamide stereoprobes might preferentially engage the guanine nucleotide-bound state of WT-*Zt*SAR1. In support of this hypothesis, we found that incubating lysates from HEK293T cells expressing WT-ZtSAR1 with GDP or the non-hydrolysable GTP analog GppCp promoted stereoselective engagement of this protein by MY-11A (**Fig. 5d, e**). This effect was stereoselective (**Fig. 5d, e**), site-specific (**Extended Data Fig. 6f**), and concentration-dependent (GDP EC_50_ = 0.85 µM, GppCp EC_50_ = 1.68 µM, **Fig. 5f, g**). An AlphaFold model places *Zt*SAR1_C64 at a location distal to the GTPase active site of *Zt*SAR1, but proximal to the N-terminal amphipathic helix that is necessary for ER membrane association (**Fig. 5h**). Analysis of *Zt*SAR1 structures modeled using Boltz-2^58^ further suggested that binding of GDP/GTP influences the conformation of the N-terminal helix (**Fig. 5h**), which could in turn explain the nucleotide-dependent ligandability of *Zt*SAR1_C64.

### Stereoprobe reactivity perturbs *Zt*SAR1 trafficking and *Z. tritici* cell growth

We next generated HEK293T cell models stably expressing eGFP fusions of WT- and C64A-*Zt*SAR1 proteins to evaluate the impact of MY-1A on the subcellular localization of *Zt*SAR1. These studies revealed that MY-1A promoted the rapid (within 2 h post-treatment) accumulation of WT-*Zt*SAR1-eGFP in ER-localized puncta (**Fig. 6a, b**). This effect was stereoselective (not observed in MY-1B-treated WT-*Zt*SAR1-eGFP cells), site-specific (not observed in MY-1A-treated C64A-*Zt*SAR1-eGFP cells) (**Fig. 6a, b**), and both concentration- (**Extended Data Fig. 7a, b**) and time-(**Extended Data Fig. 7c**) dependent. As SAR1 is vital for COPII vesicle formation and trafficking in both mammalian and fungal systems^59^, we also evaluated the effects of MY-1A on other members of the COPII complex. To accomplish this, we transiently expressed Halo tag-conjugated mammalian COPII complex proteins in WT-*Zt*SAR1-eGFP HEK293T cells and tracked their localization using a fluorescent Halo-tag ligand (JFX650)^60,61^. Using these cell models, we found that MY-1A promoted the colocalization of several COPII coat complex components^62–64^ (e.g., SEC23B, SEC31A; **Fig. 6c, d**) and COPII receptor/cargo proteins^65–68^ (e.g., TMED2, TRAPPC1, STX5, TFG; **Fig. 6c, d**) with WT-*Zt*SAR1-eGFP. Taken together, these results indicate that MY-1A reactivity with *Zt*SAR1 leads to its accretion alongside other COPII complex members and cargo proteins at specific locations in ER membranes that may represent ER exit sites^69–73^.

**Figure 6.**
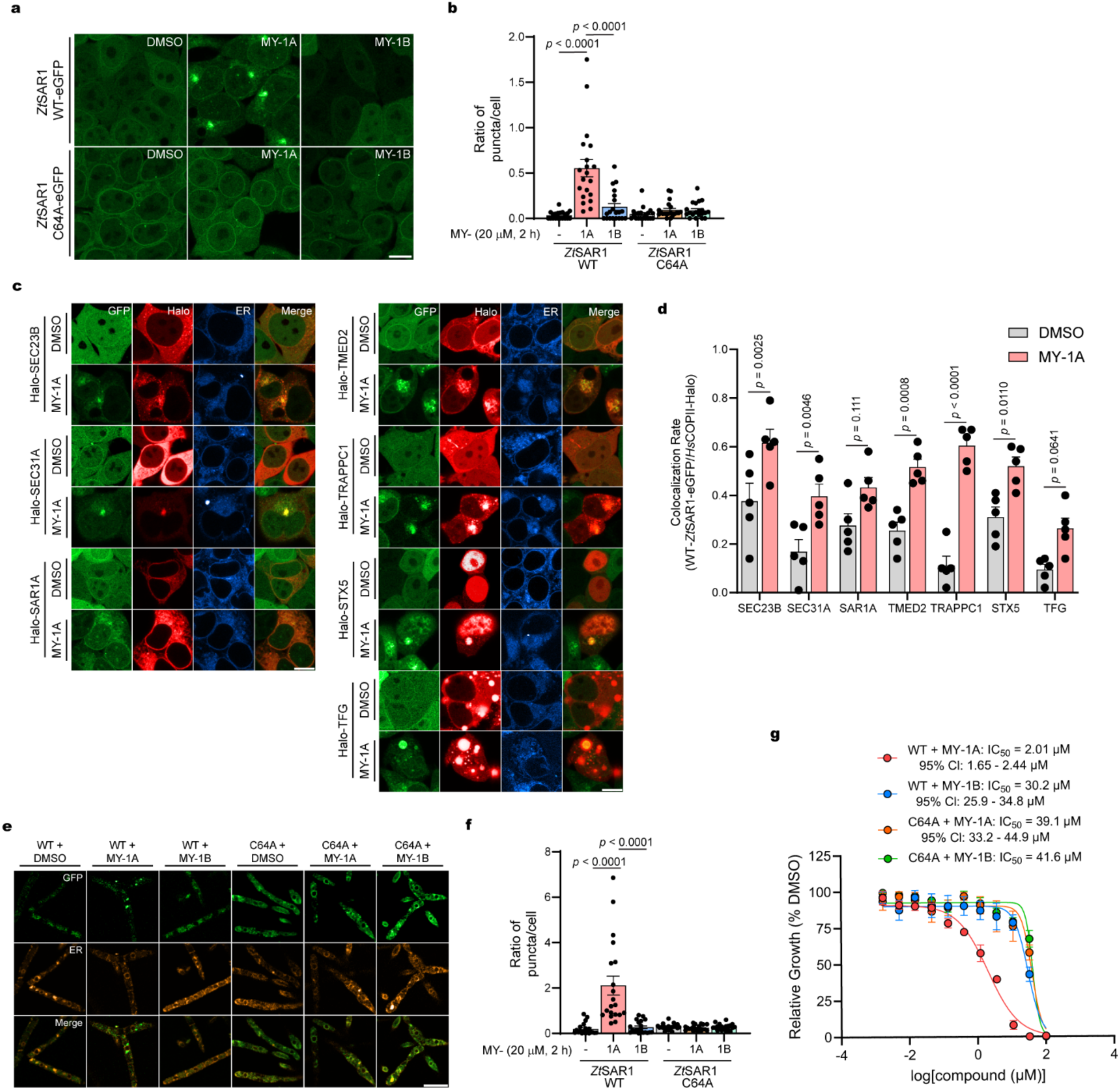
Stereoprobe MY-1A promotes accumulation of *Zt*SAR1 in ER puncta and impairs *Z. tritici* cell growth. **a**, Confocal microscopy images of recombinant *Zt*SAR1-eGFP in HEK293T cells showing aggregation of WT-, but not C64A-*Zt*SAR1-eGFP, in ER-associated puncta following treatment with MY-1A, but not MY-1B (20 µM, 2 h); scale bar = 10 µm. **b,** Quantification of GFP puncta per cell from microscopy images represented by (**a**). Data represent mean values ± s.e.m. for *n* = 20 focal planes (125 µm x 125 µm area) and *n* = 18 cells on average per focal plane; one-way ANOVA. **c,** Confocal microscopy images of co-expressed recombinant human Halo-COPII vesicle proteins with WT-*Zt*SAR1-eGFP in HEK293T cells showing increased colocalization of Halo and eGFP signal in the presence of MY-1A, but not MY-1B (20 µM, 2 h). Merge channel is the overlay of GFP and Halo channels; scale bar = 10 µm. **d,** Quantification of Pearson correlation rates from microscopy images represented by (**c**). Data represent mean values ± s.e.m. for *n* = 5 focal planes (125 µm x 125 µm area) and *n* = 21 cells on average per focal plane; two-way ANOVA. **e,** Confocal microscopy images of *Zt*SAR1 WT- and C64A-eGFP in *Z. tritici* cells showing aggregation of WT-, but not C64A-*Zt*SAR1-eGFP, in ER-associated puncta following treatment with MY-1A, but not MY-1B (40 µM, 2 h). Merge channel is the overlay of GFP and ER channels; scale bar = 10 µm. **f,** Quantification of GFP puncta per cell from microscopy images represented in (**e**). Data represent mean values ± s.e.m. for *n* = 20 focal planes (125 µm x 125 µm area) and *n* = 24 cells on average per focal plane; two-way ANOVA. **g,** Growth assay for *Z. tritici* cells showing stereoselective effects of MY-1A (0–100 µM, 7 days) in WT-, but not C64A-*Zt*SAR1-expressing cells. Data represent mean values ± s.e.m. for four independent experiments.

We extended our investigation of MY-1A effects on *Zt*SAR1 localization to *Z. tritici* cells by establishing transgenic strains that express WT- or C64A-*Zt*SAR1-eGFP under the control of a Tet-off promoter^74,75^. As we observed in HEK293T cell models, MY-1A promoted the stereoselective and site-specific accumulation of WT-*Zt*SAR1 in ER-localized puncta in *Z. tritici* cells (**Fig. 6e, f**). Considering that SAR1 depends on dynamic interactions between its N-terminal helix and ER membranes to facilitate COPII vesicle fission and uncoating for ER-to-Golgi transport^56^, and that the *S. cerevisiae* ortholog of *Zt*SAR1 is an essential protein, we next investigated how the apparent trapping of ZtSAR1 at specific sites in the ER affected *Z. tritici* cell growth. These experiments revealed that MY-1A impaired the growth of WT-*Zt*SAR1 to a much greater extent than MY-1B and this differential activity was ameliorated in C64A-*Zt*SAR1 cells (**Fig. 6g**).

We finally investigated whether the allosteric ligandable pocket in *Zt*SAR1 might exist in orthologous proteins that lack C64. Leveraging a strategy that assesses the preservation of small molecule-binding pockets in sequence-related proteins displaying non-conserved ligandable cysteines^48^, we introduced neo-cysteine mutations into the phytopathogenic ascomycete *B. cinerea* and the basidiomycete *P. graminis* SAR1 proteins at positions corresponding to *Zt*SAR1_C64 (V64C-*Bc*SAR1 and V64C-*Pg*SAR1, respectively; **Extended Data Fig. 8a**). We similarly introduced a cysteine into human SAR1A (M69C) at a position aligned with *Zt*SAR1_C64. Gel-ABPP of the parental (WT) and neo-cysteine mutant proteins expressed in HEK293T cells revealed that the V64C-*Bc*SAR1 mutant, but not WT-*Bc*SAR1 stereoselectively reacted with MY-11A, and this reaction was stereoselectively blocked by pre-treatment with MY-1A (**Extended Data Fig. 8b, c**). In contrast, the neo-cysteine mutants of *Pg*SAR1 and *Hs*SAR1A did not react with MY-11A or MY-1A stereoprobes (**Extended Data Fig. 9a-f**). Interestingly, MY-1A liganding of V64C-*Bc*SAR1 also increased the amount of this protein in the particulate fraction (**Extended Data Fig. 8b, d**) and promoted accumulation of a GFP-V64C-*Bc*SAR1 fusion in ER-localized puncta of HEK239T cells (**Extended Data Fig. 8e, f**), mirroring the effects of this compound on WT-*Zt*SAR1 (**Figs. 4j, 5a-c** and **6a**). These results thus indicate that the ligandable allosteric pocket in *Zt*SAR1 is preserved in at least a subset of additional phytopathogenic fungi, suggesting that broader spectrum pan-binding reversible ligands could also be pursued in the future, by utilizing, for instance, the nanoBRET-ABPP assay described above.

## Discussion

The emergence of multidrug-resistant field strains of *Z. tritici* combined with a limited number of fungicidal modes of action has highlighted the need to identify additional essential proteins in this phytopathogen that can be targeted by small molecules^76–79^. Here, we have shown that covalent stereoprobe ligandability maps of *Z. tritici* generated by ABPP include several proteins that are essential for fungal growth as determined by cross-referencing to the Saccharomyces Genome Database. While some of these proteins were only liganded in *Z. tritici* lysates (see below for more discussion), others like *Zt*SAR1 showed the opposite profile of exclusive engagement by stereoprobes in live *Z. tritici* cells. The restoration of stereoprobe engagement of *Zt*SAR1 in lysates in the presence of guanine nucleotides underscores the potential for small molecules to bind allosteric sites on proteins in a state-dependent manner in cells. Our findings with *Zt*SAR1 are, in this regard, reminiscent of the guanine nucleotide-dependent binding of inhibitors of human KRAS^80^ and suggest that GTPases, due to the substantial conformational changes they undergo upon GDP/GTP binding, may represent a protein class enriched in dynamic, ligandable allosteric pockets.

SAR1 is one of five core components of the COPII complex that mediates vesicular trafficking from the ER to Golgi^81,82^. SAR1 plays the specific role of acting as a membrane anchor to facilitate COPII coat assembly, and this process is initiated by GTP-dependent exposure of an *N*-terminal amphipathic helix which induces SAR1 translocation from the cytosol to the membrane^56,83^. While we do not yet understand what guanine nucleotide state of *Zt*SAR1 reacts with stereoprobes *in situ*, the increased quantity of this protein in the particulate fraction of cells treated with MY-1A may indicate that stereoprobes bind and stabilize the GTP-associated state of *Zt*SAR1. Also consistent with this conclusion is the accumulation of *Zt*SAR1 in ER-proximal puncta in MY-1A-treated cells that colocalize with other COPII proteins and cargo. This profile is reminiscent of GTP-locked H79A/G mutants of mammalian SAR1, which have also been found to co-localize with COPII complex proteins and cargo at membrane-associated puncta that have been interpreted to represent ER exit sites^60,84,85^. Our data thus, taken together, suggest that stereoprobes reacting with C64 of *Zt*SAR1 may act in a dominant-negative manner through stabilizing or trapping this protein and its COPII complex members at ER exit sites, resulting in impaired vesicular transport and corresponding blockade of *Z. tritici* growth. The lack of conservation of C64 in wheat and human SAR1 proteins points to the potential for more advanced covalent ligands targeting this site in *Zt*SAR1 to act as selective fungicides. This fungicidal effect would, however, be limited to *Z. tritici*, as other major fungal phytopathogens, such as *B. cinerea* and *P. graminis*, also lack C64. We found that a neo-cysteine-engineered *Bc*SAR1 protein did, however, react with azetidine acrylamide stereoprobes, pointing to the conservation of an allosteric ligandable pocket in this protein that might be addressable, in the future, by reversibly binding small molecules.

A broader assessment of our covalent ligandability maps of *Z. tritici* showcases both the potential opportunities and challenges facing chemical proteomic efforts to expand the landscape of candidate fungicidal targets. On the one hand, our ABPP experiments identified several essential *Z. tritici* proteins liganded by stereoprobes at cysteines that were either not conserved in the orthologous wheat and human proteins (e.g., *Zt*NMT1) or conserved, but not liganded in the human ortholog (e.g., *Zt*GSPT1). These data support that the evolutionary distance between a fungal pathogen like *Z. tritici* and plants and humans is sufficiently large to create many instances of species-restricted covalent small-molecule interactions, even on proteins that are conserved, essential regulators of cell growth. A similar finding was recently made in the pursuit of trypanocidal compounds to treat Chagas disease, where phenotypic screens identified compounds that inhibit *T. brucei* growth by covalently targeting a cysteine found in the trypanosomatid, but not human topoisomerase II enzyme^86^.

Our covalent ligandability maps further point to the potential for electrophilic small molecules to engage structurally and functionally diverse protein classes in *Z. tritici* (**Fig. 2f**) including proteins that lack discernable homologs in plants, humans, and even other yeast species (**Fig. 2h** and **3a**). Nonetheless, many of these stereoprobe-protein interactions were only observed *in vitro* due to a striking reduction in stereoprobe reactivity in *Z. tritici* cells. Although we cannot yet definitively state why the electrophilic stereoprobes showed such a dramatic loss in reactivity in *Z. tritici* cells, other studies have also reported substantial reductions in compound potency in *Z. tritici* cells that could be attributed to a rapid adaptive response to exposure to small molecules involving upregulation of cytochrome P450-related proteins^49,87^. Additionally, *Z. tritici*, as well as other phytopathogenic fungi, express multidrug efflux pumps that facilitate the cellular export of small molecules^88–90^. Future studies where adaptive response components such as transcription factors or drug-metabolizing enzymes and efflux pumps are deleted from *Z. tritici* may enhance the cellular ligandability maps generated by ABPP. Even for liganding events exclusively mapped *in vitro*, however, we believe there is a path for advancement that may involve, for instance, screening newly identified pockets for higher potency ligands with better drug-like properties. Such screens can be constructed from the stereoprobe hits themselves, as we showed for *Zt*GSPT1.

Projecting forward, it is important to emphasize that our ligandability maps of *Z. tritici* were generated with a limited number of electrophilic stereoprobes, and we expect that these maps will be further enriched by evaluating additional types of chemical proteomic probes including, for instance, compounds that assess covalent binding events at residues beyond cysteine^91–99^, as well as reversible small molecule binding events (through photoreactivity^100–103^). We also evaluated a single growth condition supporting yeast-like asexual blastospores and representing only one of several developmental stages of the complex lifecycle of *Z. tritici*^104^. It is likely that additional ligandability events will be identified in chemical proteomic investigations of other lifecycle stages of *Z. tritici*^105,106^. A more complete understanding of essential genes in *Z. tritici* would also be helpful for prioritizing ligandable proteins for functional studies. In our current investigation, we relied on the gene essentiality map from *S. cerevisiae* as a reference, which does not account for *Z. tritici* genes lacking orthologs or genes that may be essential for virulence and successful colonization of wheat plants during additional stages of the *Z. tritici* lifecycle.

In summary, our findings provide a rich resource of covalent small molecule-protein interactions in *Z. tritici, a* phytopathogen that constitutes a critical risk to global food supply. This covalent ligandability map includes an ortholog-restricted cysteine in an essential vesicular transport protein SAR1 that, upon engagement by azetidine acrylamides, sequesters the COPII signaling complex in ER-localized foci and impairs *Z. tritici* growth. We believe that our work more generally demonstrates the utility of chemical proteomics and covalent chemistry for expanding the scope of ligandable proteins in pathogenic fungi, which holds promise for the future development of new antifungal agents for both crop management and human health.

## Acknowledgements

This work was supported by Syngenta. We are grateful to Quynh Nguyen Wong and Jason Lee (Scripps Research; Automated Synthesis Facility) for HRMS measurements. We are grateful to Kathryn Spencer and Scott Henderson (Scripps Research; Core Microscopy Facility) for microscopy analysis. Molecular graphics and analyses were performed with UCSF ChimeraX, developed by the Resource for Biocomputing, Visualization, and Informatics at the University of California, San Francisco (UCSF), with support from National Institutes of Health (NIH) grant R01-GM129325 and the Office of Cyber Infrastructure and Computational Biology, National Institute of Allergy and Infectious Diseases.

## Author Contributions Statement

V.P. and B.F.C. conceptualized and supervised the study. M.Y., T.W., and A.R. generated proteomic data in *Z. tritici*. M.Y., T.W., B.M., and B.F.C. performed analysis of proteomic data.

M.Y. and T.W. performed the gel-ABPP, NanoBRET, live-cell imaging and all other experiments.

B.M., M.Y., and T.W. performed structural prediction and molecular docking experiments. C.J.R. synthesized CJR-88A//B compounds. M.Y. and E.N. generated proteomic data in human cells.

S.M. and M.Y. performed sensitivity assay of *Z. tritici* SAR1 cell lines. G.S. and S.M. designed molecular strategies and generated genetically engineered *Z. tritici* cell lines. U.T. and G.S. provided support to *Z. tritici* handling and processing. All authors contributed to writing and reviewing the manuscript.

---

**Extended Data Figure 1.**
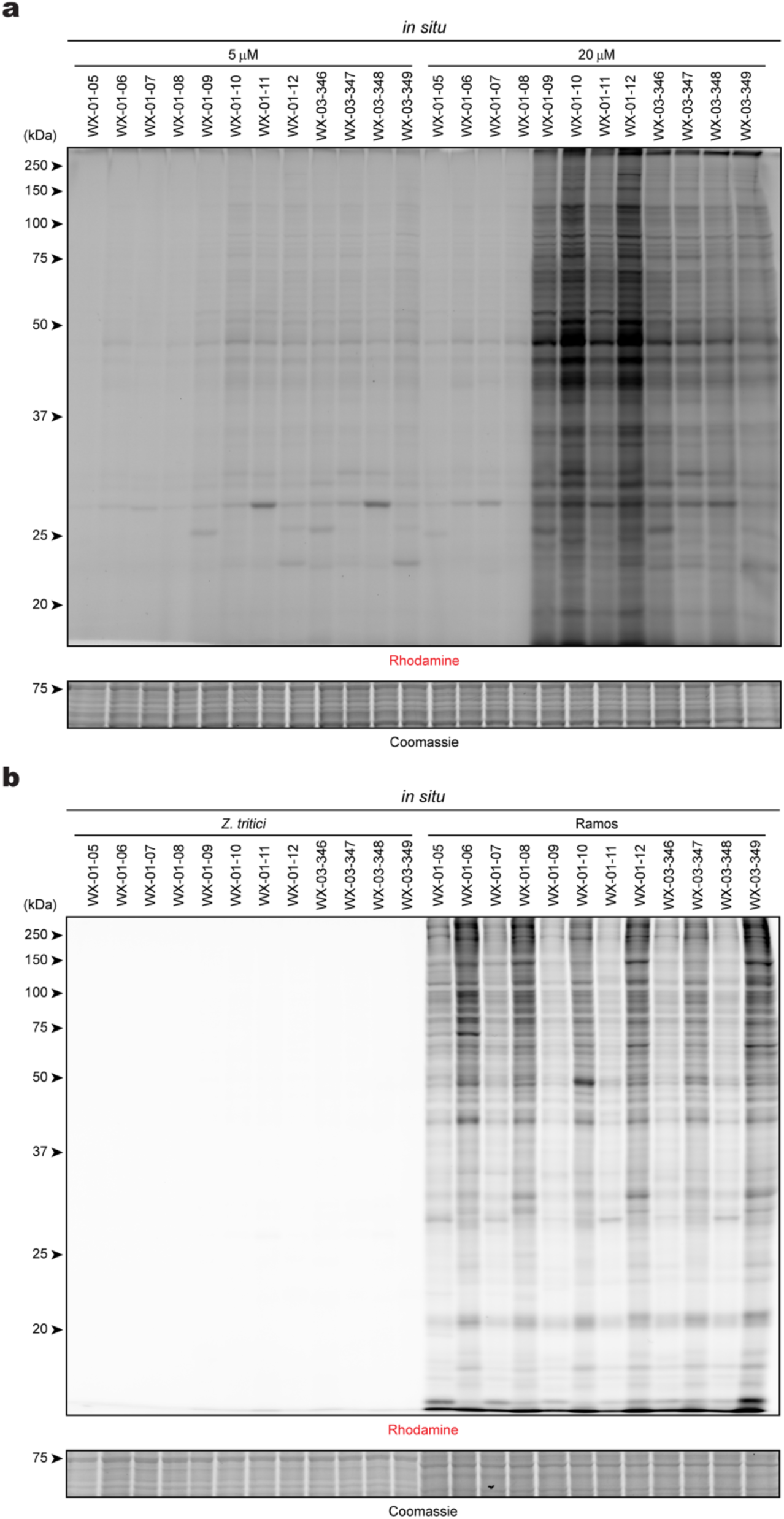
Gel-ABPP analysis of the proteome reactivity of tryptoline acrylamide stereoprobes in *Z. tritici* and Ramos cells. a, Gel-ABPP data showing the concentration-dependent *in situ* reactivity of alkynylated tryptoline acrylamide stereoprobes in *Z. tritici* (5 µM and 20 µM, 3 h). b, Gel-ABPP data showing *in situ* reactivity of alkynylated tryptoline acrylamide stereoprobes (5 µM, 3 h) in *Z. tritici* and Ramos cells. For a, b, data are from a single experiment representative of two independent experiments.

**Extended Data Figure 2.**
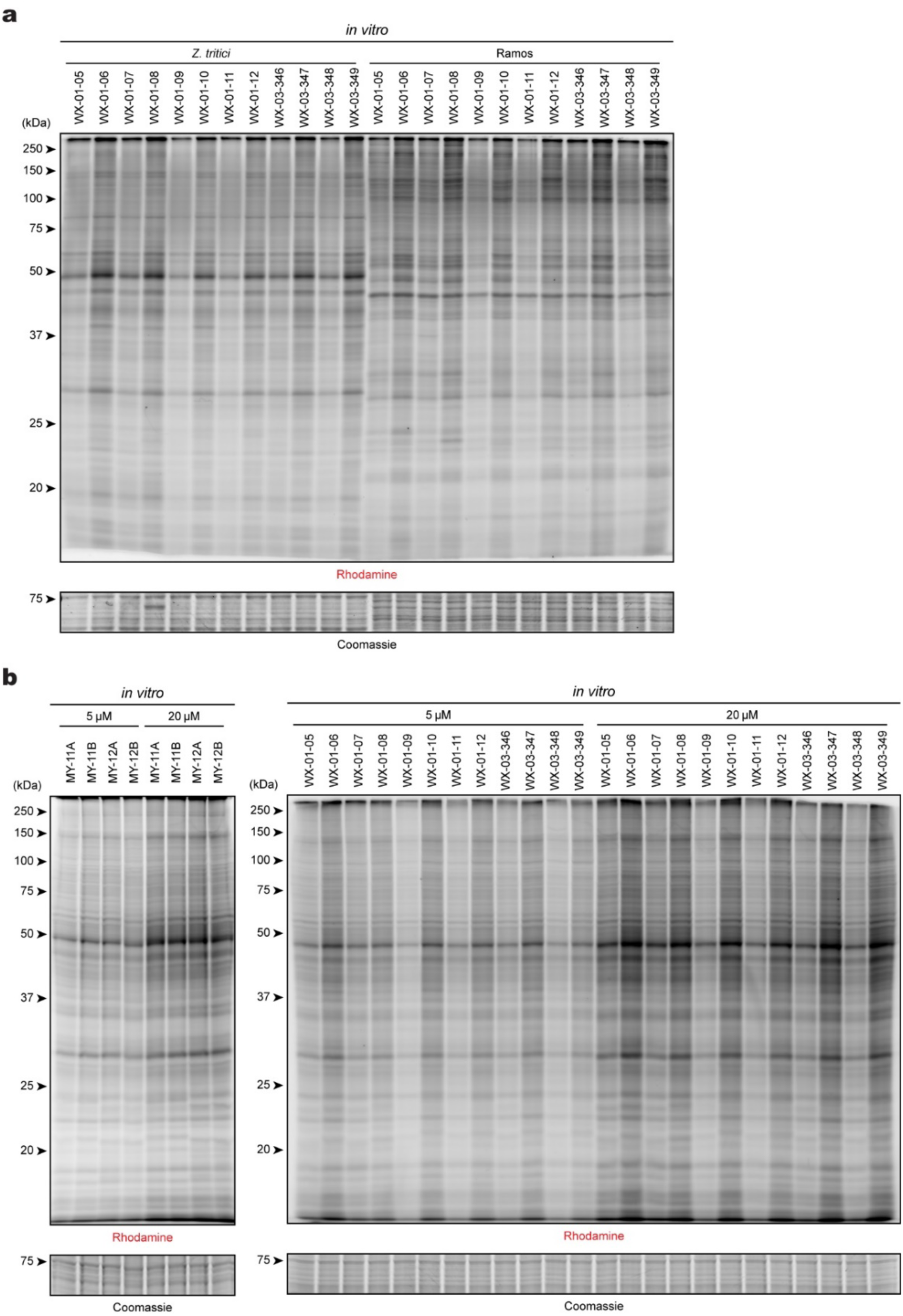
Gel-ABPP analysis of the proteome reactivity of acrylamide stereoprobes in *Z. tritici* and Ramos cell lysates. a, Gel-ABPP data showing the *in vitro* reactivity of alkynylated tryptoline acrylamide stereoprobes (5 µM, 3 h) in *Z. tritici* and Ramos cell lysates. b, Gel-ABPP data showing the concentration-dependent *in vitro* reactivity of alkynylated azetidine (left) or tryptoline (right) acrylamide stereoprobes (5 µM and 20 µM, 3 h) in *Z. tritici* lysates. For a, b, data are from a single experiment representative of two independent experiments.

**Extended Data Figure 3.**
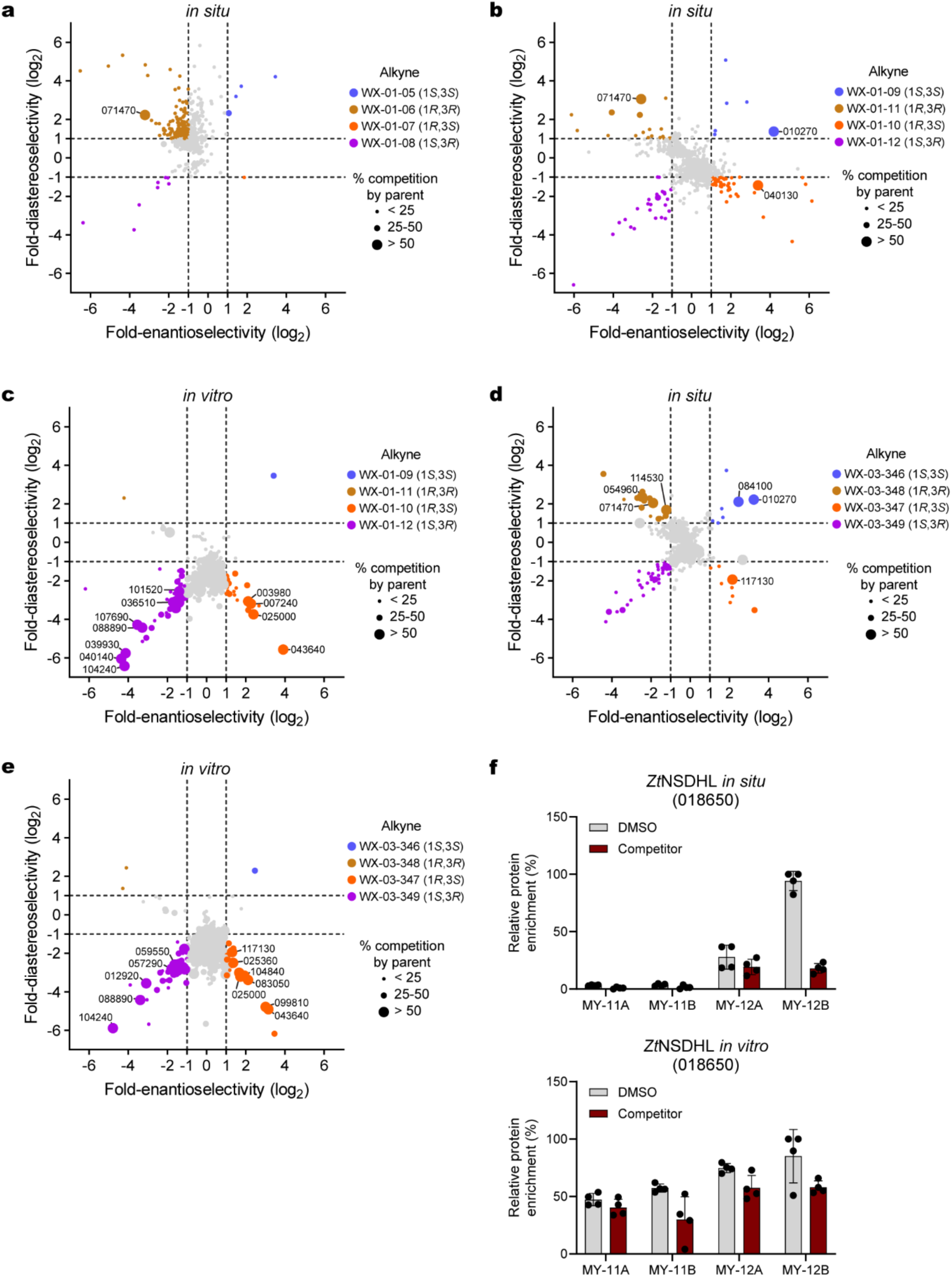
Mapping stereoprobe-liganded proteins and cysteines in *Z. tritici* by ABPP. a-e, Protein-directed ABPP data shown as quadrant plots for tryptoline acrylamide stereoprobes: a, parent: WX-02-16/26/36/46; alkyne: WX-01-05/06/07/08; *in situ*; b, parent: EV-98/96/99/97; alkyne: WX-01-09/10/11/12, *in situ*, c, parent: EV-98/96/99/97; alkyne: WX-01-09/10/11/12; *in vitro*; d, parent: WX-03-57/58/59/60; alkyne: WX-03-346/347/348/349; *in situ*; and e, parent: WX-03-57/58/59/60; alkyne: WX-03-346/347/348/349; *in vitro*. In all cases, *Z. tritici* cells were treated with parent stereoprobes (40 µM, 2 h) followed by their corresponding alkynylated stereoprobes (10 µM, 1 h). Representative stereoprobe-liganded proteins are denoted in each quadrant plot by their gene IDs (i.e., ZtIPO323_XXXXXX). Data are average values from four independent experiments. f, Protein-directed ABPP data showing stereoselective liganding of *Zt*NSDHL (ZtIPO323_018650) by azetidine acrylamides *in situ* (top) but not *in vitro* (bottom). Data represent mean values ± s.d. of four independent experiments.

**Extended Data Figure 4.**
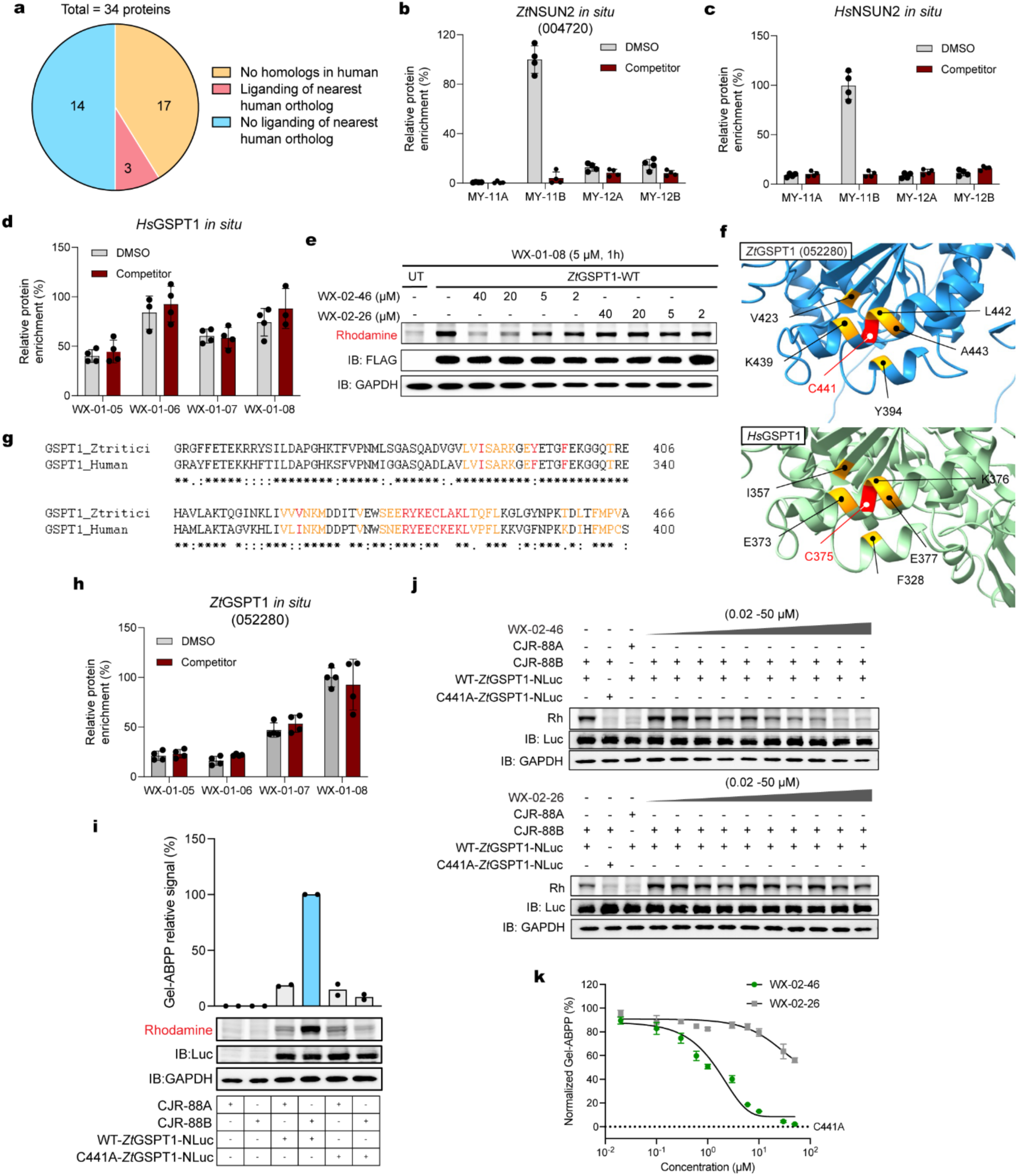
Additional characterization of stereoprobe-liganded essential proteins from *Z. tritici*. a, Pie chart showing fraction of *in situ* stereoprobe-liganded cysteines/proteins from *Z. tritici* for which the nearest human homolog is also liganded. b, Protein-directed ABPP data showing stereoselective liganding of *Zt*NSUN2 by the parent/alkyne stereoprobe pair MY-1B/MY-11B (40 µM, 2 h; 10 µM, 1 h). Data are from *in situ* treatment of *Z. tritici* cells. c, Protein-directed ABPP data showing stereoselective liganding of *Hs*NSUN2 by the alkyne/parent stereoprobe pair MY-11B/MY-1B (5 µM, 1 h; 20 µM, 2 h pre-treatment). Data are from *in situ* treatments of Ramos cells. d, Protein-directed ABPP data showing lack of stereoselective liganding of *Hs*GSPT1 by the alkyne/parent stereoprobe pair WX-01-08/ WX-02-46 (10 µM, 1 h; 40 µM, 2 h pre-treatment). Data are from *in situ* treatments of Ramos cells. e, Gel-ABPP data showing concentration-dependent, stereoselective blockade of WX-01-08 (5 µM, 1 h) engagement of WT-*Zt*GSPT1 by pre-treatment with WX-02-46 (2-40 µM, 2 h) in HEK293T cells. Data are from a single experiment representative of two independent experiments. f, Structural comparison of the predicted AlphaFold structure of *Zt*GSPT1 (AF-F9X8V1-F1) and crystal structure of *Hs*GSPT1 (PDB ID: 3J5Y) highlighting residues around C441 of *Zt*GSPT1. g, Sequence alignment of *Zt*GSPT1 and *Hs*GSPT1, highlighting proximal residues around the stereoprobe-liganded cysteine C441 in *Zt*GSPT1. The residues are colored in red if within 5 Å of C441 or orange if within 10 Å. h, Protein-directed ABPP data showing stereoselective enrichment of *Zt*GSPT1 by WX-01-08 (10 µM, 1 h) and lack of blockade of this enrichment by pre-treatment with WX-02-46 (40 µM, 2 h). Data are from *in situ* treatment of *Z. tritici* cells. For d, h, data represent mean values ± s.d. of four independent experiments. i, Gel-ABPP data showing stereoselective engagement of recombinant WT-*Zt*GSPT1-NLuc but not C441A-*Zt*GSPT1-NLuc by CJR-88B (2 µM, 1 h; bottom, representative gel-ABPP data; top, quantification of data from two independent experiments). j, Gel-ABPP data showing concentration-dependent, stereoselective blockade of CJR-88B engagement of WT-*Zt*GSPT1-NLuc by WX-02-46. Lysates from HEK293T cells transfected with WT-*Zt*GSPT1-NLuc or C441A-*Zt*GSPT1-NLuc were treated with WX-02-26 or WX-02-46 (0.02–50 µM, 2 h) followed by CJR-88B (2 µM, 1 h) and gel-ABPP analysis. Data are from a single experiment representative of three independent experiments. k, Quantification of gel-ABPP data shown in (j) wherein data are the average values ± s.e.m. from three independent experiments.

**Extended Data Figure 5.**
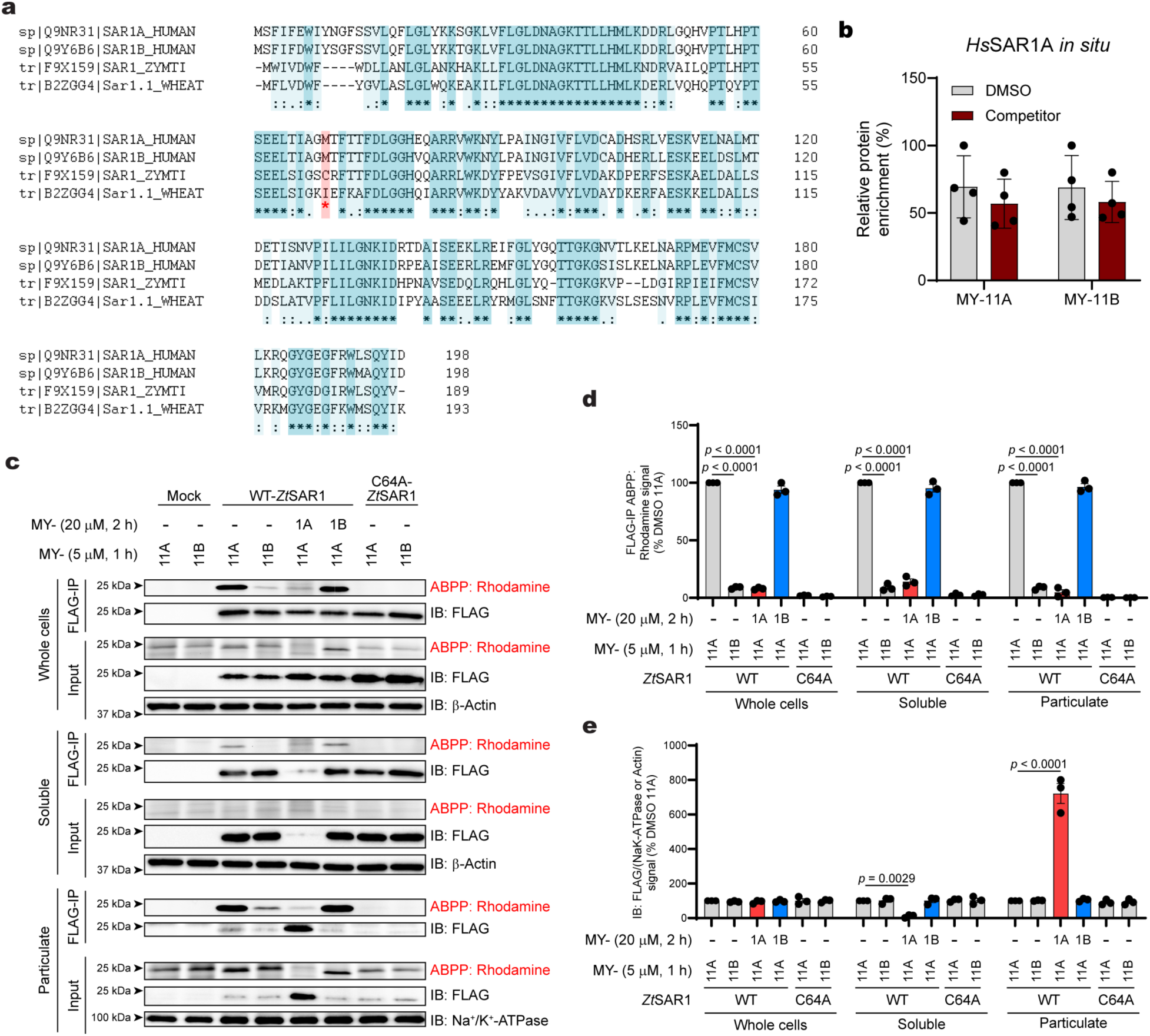
Additional characterization of an ortholog-restricted, stereoprobe-liganded cysteine in *Zt*SAR1. a, Sequence alignment of *Z. tritici* SAR1 with other SAR1 orthologs from human and wheat (*T. aestivum*). Aligned residues showing identical (opaque) and similar (transparent) sequences are highlighted. b, Protein-directed ABPP data showing a lack of stereoselective enrichment of human SAR1A from Ramos cells treated with MY-11A or MY-11B (5 µM, 3 h). Data represent mean values ± s.d. for four independent experiments. c, Gel-ABPP data showing stereoselective engagement of recombinant WT-*Zt*SAR1 by MY-11A (5 µM, 1 h) and stereoselective blockade of this engagement by pre-treatment with MY-1A (20 µM, 2 h) in HEK293T cells. After stereoprobe treatments, cells were lysed and analyzed by gel-ABPP as either whole cell lysates or as soluble and particulate fractions. d, Quantification of gel-ABPP data shown in (c). e, Quantification of *Zt*SAR1 immunoblotting signals shown in (c). For d, e, data represent mean values ± s.e.m. for three independent experiments; two-way ANOVA.

**Extended Data Figure 6.**
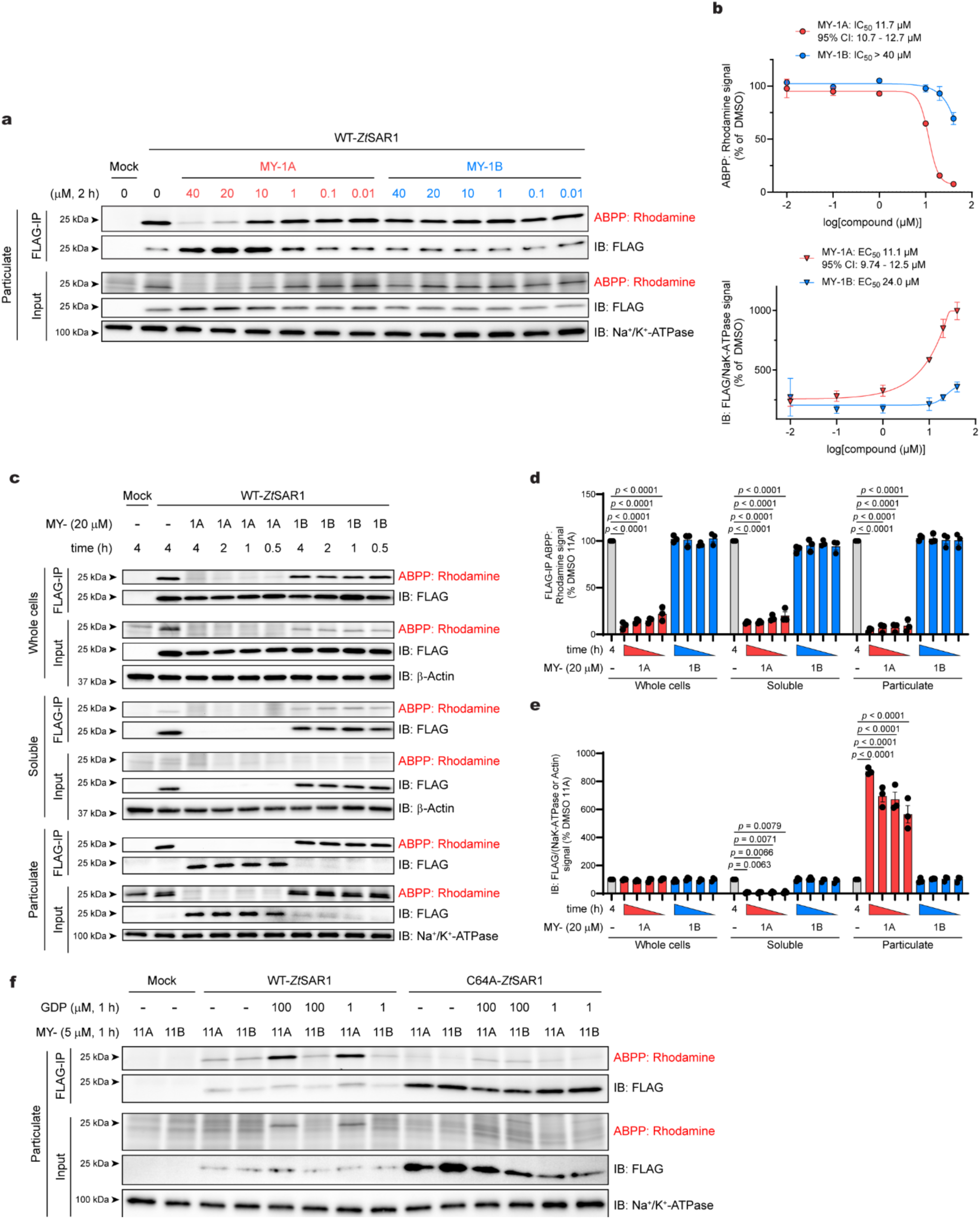
Additional characterization of an ortholog-restricted, stereoprobe-liganded cysteine in *Zt*SAR1. a, Gel-ABPP data showing concentration-dependent, stereoselective blockade of MY-11A (5 µM, 1 h) engagement of recombinant WT-*Zt*SAR1 by MY-1A (0.01-40 µM, 2 h), along with concomitant accumulation of WT-*Zt*SAR1 in the particulate fraction of HEK293T cells. b, Quantification of gel-ABPP data (top graph) and *Zt*SAR1 immunoblotting signals (bottom graph) shown in (a). Data represent mean values ± s.d. for three independent experiments. c, Gel-ABPP data showing engagement of recombinant WT-*Zt*SAR1 by MY-11A (5 µM, 1 h) and time-dependent stereoselective blockade of this engagement by MY-1A (20 µM) in HEK293T cells. After stereoprobe treatments, cells were lysed and analyzed by gel-ABPP as either whole cell lysates or as soluble and particulate fractions of these lysates. d, Quantification of gel-ABPP data shown in (c). e, Quantification of *Zt*SAR1 immunoblotting signals shown in (c). For d, e, data represent mean values ± s.e.m. for three independent experiments; two-way ANOVA. f, Gel-ABPP data showing increases in MY-11A (5 µM, 1 h) engagement of WT-*Zt*SAR1, but not C64A-*Zt*SAR1 in the presence of GDP cofactor (1 and 100 µM, 1 h). Data are from a single experiment representative of three independent experiments.

**Extended Data Figure 7.**
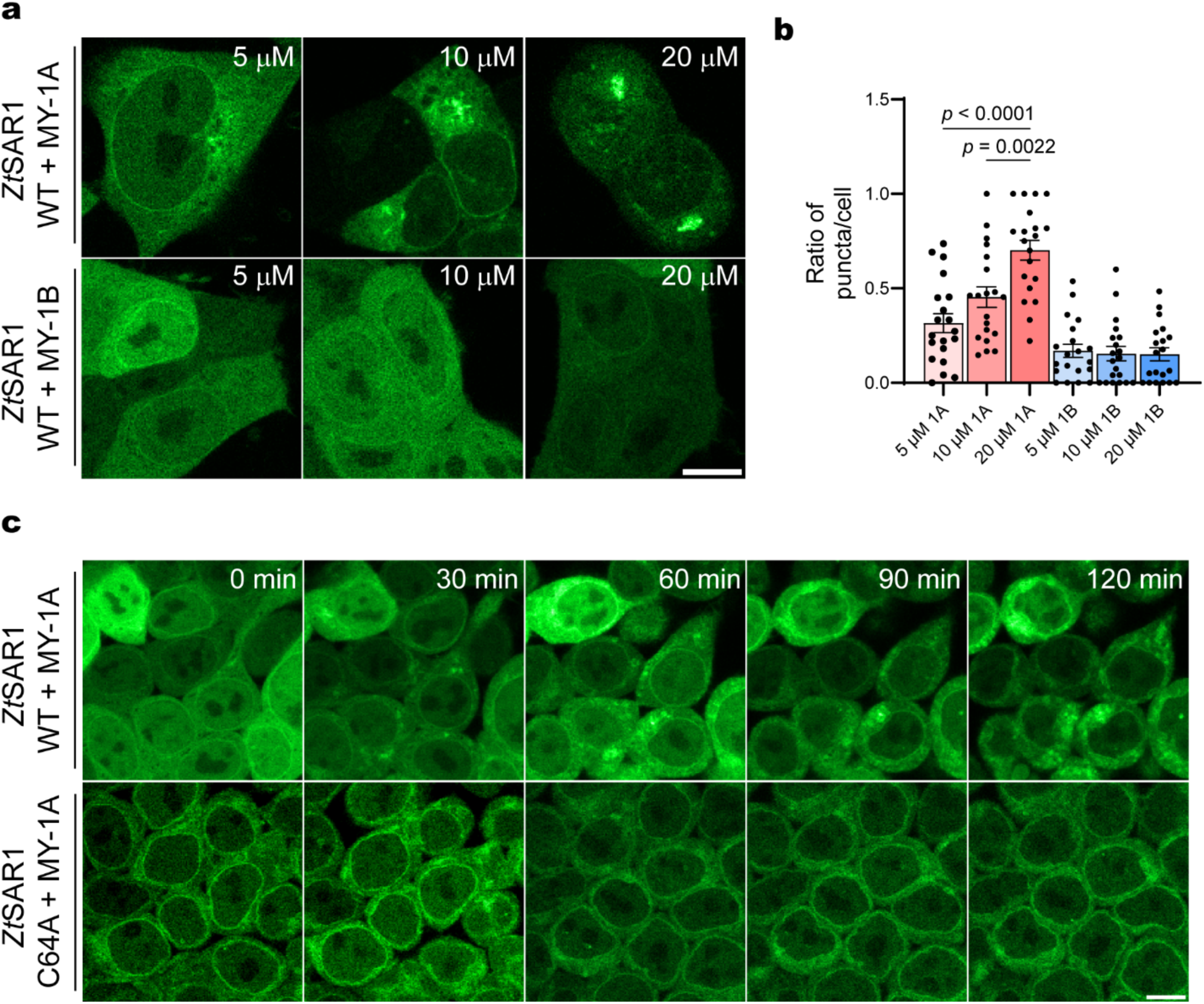
Stereoprobe MY-1A promotes accumulation *Zt*SAR1 in ER puncta. a, Confocal microscopy images demonstrating concentration-dependent aggregation of WT-*Zt*SAR1-eGFP stably expressed in HEK293T cells treated with MY-1A, but not MY-1B (2 h treatments); scale bar = 10 µm. b, Quantification of GFP puncta per cell from microscopy images represented in (a). Data represent mean values ± s.e.m. *n* = 20 focal planes (125 µm x 125 µm area); *n* = 17 cells on average per focal plane; one-way ANOVA. c, Timelapse of confocal microscopy images of WT-*Zt*SAR1-eGFP and C64A-*Zt*SAR1-eGFP stably expressed in HEK293T cells following treatment with MY-1A (20 µM); scale bar = 10 µm.

**Extended Data Figure 8.**
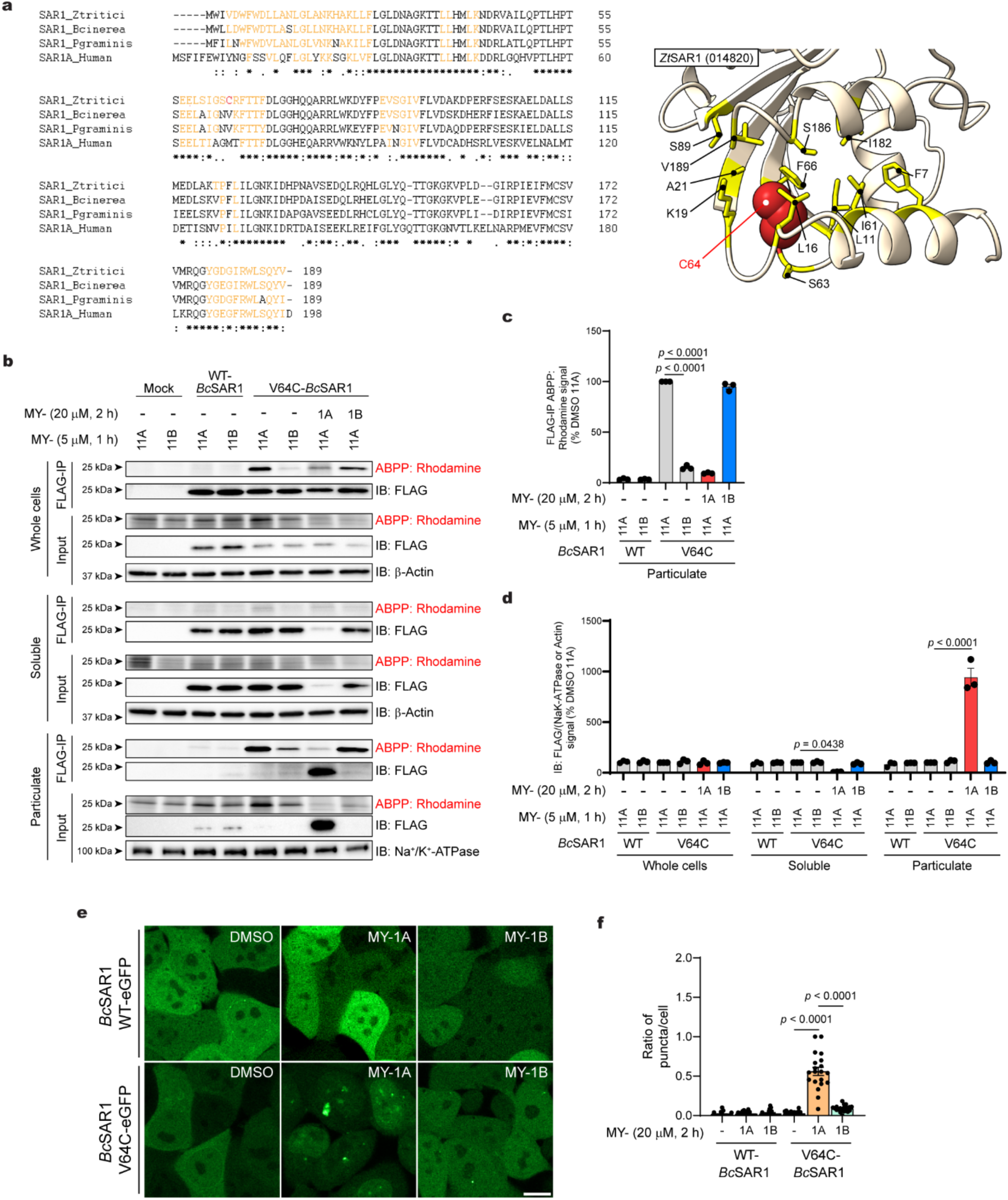
Assessing stereoprobe ligandability of a neo-cysteine *B. cinerea* SAR1 variant. a, Left, Sequence alignment of *Z. tritici* SAR1 with other SAR1 orthologs (*B. cinerea*, *P. graminis*, and *H. sapiens*) showing conserved residues (yellow) that are proximal (<15 Å) to C64 of *Zt*SAR1. Right, AlphaFold-predicted structure of *Zt*SAR1 (AF-F9X159-F1) highlighting locations of conserved residues proximal (< 15 Å) to C64. b, Gel-ABPP data showing stereoselective engagement of V64C-, but not WT-BcSAR1 by MY-11A (5 µM, 1 h) and stereoselective blockade of this engagement by pre-treatment with MY-1A (20 µM, 2 h) in HEK293T cells. After stereoprobe treatments, cells were lysed and analyzed by gel-ABPP as either whole cell lysates or as soluble and particulate fractions of these lysates. c, Quantification of gel-ABPP data for the particulate fraction shown in (b). d, Quantification of *Bc*SAR1 immunoblotting signals (b). For c, data represent mean values ± s.e.m. n = 3; unpaired two-tailed *t* test. For d, data represent mean ± s.e.m. *n* = 3; two-way ANOVA. e, Confocal microscopy images of recombinant *Bc*SAR1-eGFP demonstrating stereoselective aggregation of V64C-, but not WT-*Bc*SAR1-eGFP, by MY-1A (20 µM, 2 h) in HEK293T cells; scale bar = 10 µm. f, Quantification of GFP puncta per cell from microscopy images represented in (e). Data represent mean values ± s.e.m. *n* = 20 focal planes (125 µm x125 µm area) and *n* = 15 cells on average per focal plane; one-way ANOVA.

**Extended Data Figure 9.**
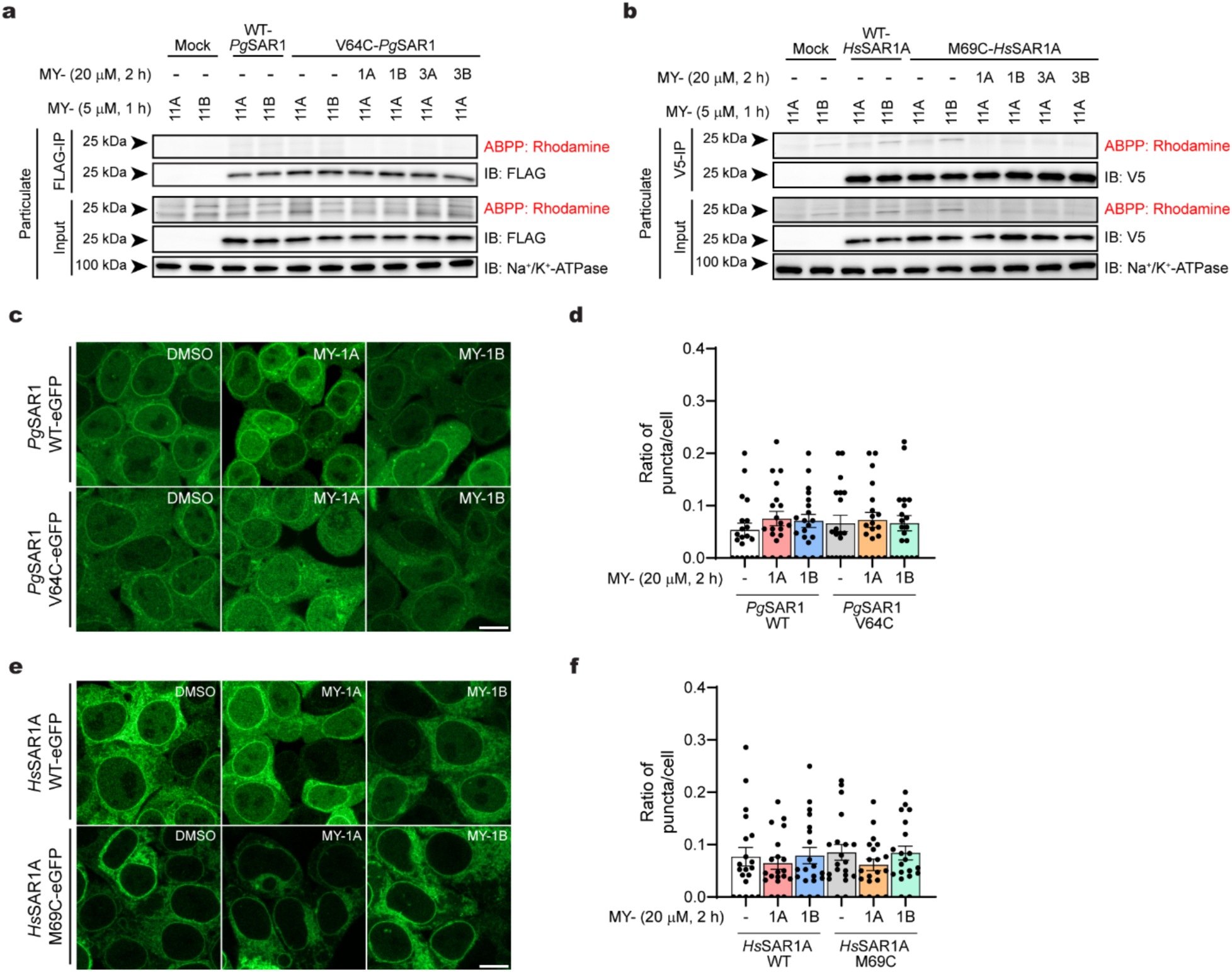
Assessing stereoprobe ligandability of neo-cysteine *P. graminis and H. sapiens* SAR1 variants. a, b, Gel-ABPP data showing lack of MY-11A (5 µM, 1 h) engagement of WT or neo-cysteine variants of *Pg*SAR1 (a) and *Hs*SAR1A (b) proteins. Neo-cysteine variants; V64C-*Pg*SAR1, M69C-*Hs*SAR1A. Data are from a single experiment representative of three independent experiments. c-f, Confocal microscopy images demonstrating lack of stereoselective aggregation of WT- or neo-cysteine variants of *Pg*SAR1-eGFP and *Hs*SAR1A-eGFP by MY-1A (20 µM, 2 h) in HEK293T cells. c, e, representative confocal images; scale bar = 10 µm; d and f, quantification of GFP puncta per cell from microscopy images represented in (c) and (e), respectively. Data represent mean values ± s.e.m. *n* = 20 focal planes (125 µm x 125 µm area) and *n* = 16 (*Pg*SAR1-eGFP) or 19 (*Hs*SAR1A-eGFP) cells on average per focal plane; one-way ANOVA.

## METHODS

### Chemical synthesis

Details on chemical synthesis can be found in the **Supplementary Information.**

### Cell lines and cell culture

HEK293T (ATCC, CRL-3216, human, female) were grown and maintained in high-glucose DMEM (Corning) supplemented with 10% fetal bovine serum (Omega Scientific), 2 mM GlutaMAX (Gibco), penicillin (100 U mL^-1^), and streptomycin (100 µg mL^-1^) at 37 °C in a humidified 5% CO_2_ atmosphere. Ramos (ATCC, CRL-1596, human, male) were grown in RPMI (Gibco) supplemented with 10% fetal bovine serum, 2 mM GlutaMAX, penicillin (100 U mL^-1^), and streptomycin (100 µg mL^-1^) at 37 °C in a humidified 5% CO_2_ atmosphere. *Zymoseptoria tritici* spores (IPO323 strain) were grown in potato dextrose broth (PDB) media (Difco^TM^) and propagated in 150 mL shaking flasks at 200 rpm and 25 °C (r.t.) for 3 days.

### Gel-ABPP for proteome-wide reactivity of stereoprobes

#### *In situ* treatment and sample processing

*Z. tritici* cells (7 mL of 1 x 10^6^ cells mL^-1^) were treated in 6-well suspension dishes (r.t., shaking at 150 rpm) with DMSO (0.1% v/v) or alkyne stereoprobe (5–20 µM, 3 h). Cells were harvested via pipetting on ice, washed twice with cold DPBS and stored at −80 °C. The cell pellets were resuspended in 500 μL cold lysis buffer (50 mM HEPES pH 7.5, 150 mM NaCl, and 0.1% Triton X-100; Sigma-Aldrich, X100) with EDTA-free complete protease inhibitor cocktail (Roche, 04693159001). Cells were lysed by glass bead milling (0.5 mm diameter) using a Bullet Blender® homogenizer instrument (speed setting 7, 5 min, 4 °C), followed by probe pulse sonication using a Branson Ultrasonics Sonifier S-250A cell disruptor (15 pulses, 10% power output, 1s on/1s off) and centrifuged (1000 x *g*, 5 min, 4 °C). 50 µL of 1 mg mL^-1^ of normalized (BSA standard) whole cells lysates (DC protein assay, Bio-Rad) were treated with 5.5 µL of click master mix [1 µL of 50 mM CuSO_4_ in H_2_O, 3 µL of 1.7 mM TBTA in *t*-BuOH:DMSO (4:1, v/v), 1 µL of freshly prepared 50 mM Tris(2-carboxyethyl)phosphine (TCEP) in H_2_O, 1 µL of 1.25 mM TAMRA-N_3_ in DMSO] and incubated at r.t. for 1 h. Afterwards, 17 µL of 4x Laemmli loading dye was added and samples were resolved by SDS-PAGE (in-house casted 10% acrylamide long gel, 275 V, 3 h) and visualized by in-gel fluorescence scanning on a ChemiDoc MP Imaging System (Bio-Rad). The images were processed using Image Lab software (v6.1).

#### Modifications for mammalian suspension cells

Ramos cells (10 mL of 3 x 10^6^ cells mL^-1^; seeded 3 h before treatment) were treated (37 °C, 5% CO_2_) with alkyne stereoprobe (5 - 20 µM, 3 h). Cells were harvested by centrifugation (2000 x *g*, 3 min), washed twice with cold DPBS and stored at −80 °C. The cell pellets were resuspended in 200 µL cold lysis buffer (DPBS) with EDTA-free complete protease inhibitor cocktail. Cells were lysed by probe pulse sonication (15 pulses, 10% power output, 1s on/1s off). 50 µL of 1 mg mL^-1^ of normalized whole cells lysates were further processed for click reaction and SDS-PAGE long gel analysis as described as above.

#### *In vitro* treatment and sample processing

Cell lysates (50 µL, 1 mg mL^-1^) were treated with 1 µL of 50x alkyne probe (5–20 µM final, 3 h) at r.t., followed by treatment with 5.5 µL of click master mix, and then resolved by SDS-PAGE long gel analysis as described above.

### Protein-directed MS-ABPP

#### *In situ* treatment and sample processing of *Z. tritici* proteome

*Z. tritici* cells (1 x 10^6^ cells mL^-1^) were seeded in 150 mL flasks (100 mL of PDB) and incubated for 3 days (r.t., shaking at 150 rpm). Afterwards, 7 mL of *Z. tritici* cells were treated with DMSO (0.1% v/v) or parent stereoprobe (40 µM, 2 h), followed by treatment with a stereochemically matched alkyne probe (10 µM, 1 h). Cells were harvested via pipetting on ice, washed twice with cold DPBS and stored at −80 °C. The cell pellets were resuspended in 500 μL cold lysis buffer (50 mM HEPES pH 7.5 and 150 mM NaCl) with EDTA-free complete protease inhibitor cocktail. Cells were lysed by glass bead milling (0.5 mm diameter) using a Bullet Blender® homogenizer instrument (speed setting 7, 5 min, 4 °C), followed by probe pulse sonication (15 pulses, 10% power output, 1s on/1s off) and centrifuged (1000 x *g*, 5 min, 4 °C). 500 µL of 2 mg mL^-1^ of normalized (BSA standard) whole cells lysates (Pierce BCA protein assay, Thermo) were treated with 55 μL of click MS-Master-mix [10 µL of 50 mM CuSO_4_ in H_2_O, 30 µL of 1.7 mM TBTA in 4:1 *t*-BuOH:DMSO, 10 µL of freshly prepared 50 mM TCEP in H_2_O, 10 µL of 10 mM Biotin-PEG4-azide]. Proteins were precipitated with cold methanol (600 μL), chloroform (200 μL) and water (100 μL), vortexed, and then centrifuged (16,000 x *g*, 10 min, 4 °C). The top and bottom layers were aspirated, and the protein disk was sonicated in 1 mL of methanol and pelleted (16,000 x *g*, 10 min, 4 °C). After the methanol was completely aspirated, protein pellets were immediately processed or stored at −80 °C. Pellets were resuspended in 500 μL freshly prepared 8 M urea in DPBS, followed by the addition of 10 μL of 10 wt% SDS. Samples were then probe pulse sonicated until clear (10% power output, 1s on/1s off). The samples were reduced with 25 μL of 200 mM dithiothreitol (DTT) at 65 °C for 15 min, followed by alkylation with 25 μL of 400 mM iodoacetamide for 30 min at 37 °C. Then, 100 μL of 10 wt% SDS was added, and the samples were transferred to a 15 mL tube in a total volume of 5.5 mL with DPBS. Washed streptavidin beads (Thermo, #20353; 100 μL of 50% slurry/sample) were then added and proteins were enriched for 2 h at r.t. with rotation. After incubation, the beads were pelleted (2 min at 2,000 x *g*) and washed with 0.2% wt% SDS in DPBS (2 x 10 mL), DPBS (1 x 5 mL), HPLC-grade water (2 x 1 mL) and 200 mM 4-(2-hydroxyethyl)-1-piperazinepropanesulfonic acid (EPPS; 1 mL, pH 8.0). Enriched proteins were digested on-bead overnight with 200 μL of trypsin mix [2 M urea, 1 mM CaCl_2_, 10 μg mL^-1^ trypsin (Promega, #V5111) in 200 mM EPPS, pH 8.0]. The beads were pelleted at 2,000 x *g*, the supernatant was collected and then diluted with 100 μL acetonitrile (30% final). Samples were then labeled with 6 μL of 20 mg mL^-1^ (in dry acetonitrile) TMTpro^16^plex tag (Thermo, #A44520) for 1 h at r.t (vortexing every 15 min). TMT labelling was quenched by the addition of hydroxylamine (6 μL of a 5% solution in H_2_O) and incubated for 15 min at r.t.. Samples were then acidified with 20 μL formic acid, combined and dried using a SpeedVac at 46 °C. Samples were desalted with a Sep-Pak column Vac 1 cc (50 mg) (Waters, #WAT054955) and then fractionated into 12 fractions using an acetonitrile/NH_4_HCO_3_ (10 mM) HPLC gradient (see HPLC fractionation) and analyzed by mass spectrometry (see TMT LC-MS/MS Analysis).

#### Modifications for mammalian suspension cells

Ramos cells (12.5 mL of 3 million cells mL^-1^; seeded 3 h before treatment) were treated (37 °C, 5% CO_2_) with DMSO (0.01% v/v) or parent stereoprobe for (20 µM, 2 h), followed by treatment with stereochemically matched alkyne probe (5 µM, 1 h). Cells were harvested by centrifugation (2000 x *g*, 3 min, r.t.), washed twice with cold DPBS and stored at −80 °C. The cell pellets were resuspended in 500 µL cold lysis buffer (DPBS) with EDTA-free complete protease inhibitor cocktail. Cells were lysed by probe pulse sonication (15 pulses, 10% power output, 1s on/1s off). 500 µL of 2 mg mL^-1^ normalized whole cells lysates (Pierce BCA protein assay) were further processed for protein-directed MS ABPP analysis as described above.

#### *In vitro* treatment and sample processing

Cell lysates (500 µL, 2 mg mL^-1^) of proteome were treated with 5 µL of 100x parent stereoprobe (40 µM final, 2 h, r.t.), followed by 5 µL of 100x alkyne probe (10 µM final, 1 h, r.t., final DMSO 2% v/v) and processed for protein-directed MS ABPP analysis as described above.

#### Data processing

Raw files were uploaded to the Integrated Proteomics Pipeline (IP2, version 6.7.1) available at http://ip2.scripps.edu/ip2/mainMenu.html, and MS2 and MS3 files were extracted from the raw files using RAW Converter (version 1.1.0.22, available at http://fields.scripps.edu/rawconv/) and searched using the ProLuCID algorithm using a reverse concatenated, non-redundant variant of the *Z. tritici* proteome generated from the predicted protein database (available at https://mycocosm.jgi.doe.gov/Zymtr1/Zymtr1.home.html)^107^ and the ProLuCID algorithm using a reverse concatenated, non-redundant variant of the Human UniProt database (release 2016-07). Cysteine residues were searched with a static modification for carboxyamidomethylation (+57.02146 Da). N termini and lysine residues were also searched with a static modification corresponding to the TMT tag (+304.2071 Da for 16-plex). Peptides were required to be at least six amino acids long. ProLuCID data were filtered through DTASelect (version 2.0) to achieve a spectrum false-positive rate below 1%. We included a keratin filter, accepted non-unique peptides, required at least one tryptic cleavage site and two peptides per protein. The MS3-based peptide quantification was performed with reporter ion mass tolerance set to 20 ppm with the Integrated Proteomics Pipeline (IP2).

#### Data analysis

Enrichment ratios (probe versus probe) were calculated for each peptide-spectrum match (PSM) by dividing the TMT reporter ion intensity by the total intensity across all channels. PSMs were grouped by protein ID, excluding peptides with summed reporter ion intensities <10,000, signal-to-noise ratio (S/N) <1.0, or isolation purity <0.5. Proteins supported by fewer than two peptides were also excluded. Replicate channels were grouped across each experiment, and mean values were calculated for each protein. To assess measurement variability, the coefficient of variation (CV), defined as the ratio of the standard deviation to the mean, was calculated across replicate channels. Proteins with a CV ≥ 0.2 in the most enriched (DMSO + alkyne) channel were excluded from further analysis. A variability metric was calculated across replicate channels as the median absolute deviation divided by the mean and expressed as a percentage.

#### Criteria for liganding in protein-directed ABPP

Proteins were initially defined as stereoselectively liganded if at least one of the following criteria were met: (1) the average enrichment by the alkyne probe was greater than 3-fold that of its enantiomer and greater than 2-fold the enrichment observed following treatment with a stereochemically matched non-alkyne competitor; (2) the variability is less than 20% and the average enrichment by the alkyne probe was greater than 2-fold that of its enantiomer and greater than 2-fold the enrichment observed following treatment with a stereochemically matched non-alkyne competitor, and at least one site in the protein was deemed liganded in cysteine-directed ABPP experiments. All proteins passing the initial filters for liganding were manually reviewed to remove proteins showing additional evidence of high variability.

### Cysteine-directed ABPP

#### *In situ* treatment and sample processing

*Z. tritici* cells (1 x 10^6^ cells mL^-1^) were seeded in 150 mL flasks (100 mL of PDB) and incubated for 3 days (r.t., shaking at 150 rpm). Afterwards, 7 mL of *Z. tritici* cells were treated with DMSO (0.1% v/v) or parent stereoprobe for (40 µM, 2 h). Cells were harvested via pipetting on ice, washed twice with cold DPBS and stored at −80 °C. The cell pellets were resuspended in 500 μL cold lysis buffer (50 mM HEPES pH 7.5 and 150 mM NaCl) with EDTA-free complete protease inhibitor cocktail. Cells were lysed by glass bead milling (0.5 mm diameter) using a Bullet Blender® homogenizer instrument (speed setting 7, 5 min, 4 °C), followed by probe pulse sonication (15 pulses, 10% power output, 1s on/1s off) and centrifuged (1000 x *g*, 5 min, 4 °C). 500 µL of 2 mg mL^-1^ of normalized whole cells lysates (Pierce BCA protein assay) were treated with 5 μL of 50 mM IA-DTB (500 µM final, 1 h, r.t.) with vortexing every 20 min. Proteins were precipitated with cold methanol (600 μL), chloroform (200 μL) and HPLC-grade water (100 μL), followed by vortexing and centrifugation (16,000 x *g*, 10 min, 4 °C). Without disrupting the protein disk, both the top and bottom layers were aspirated, and the protein disk was sonicated again in 1 mL of methanol and pelleted (16,000 x *g*, 10 min, 4 °C). After the methanol was completely aspirated, protein pellets were immediately processed or frozen at −80 °C. Pellets were resuspended in 90 μL of denaturing/reducing buffer [9 M urea, 10 mM DTT, 50 mM triethylammonium bicarbonate (TEAB) pH 8.5]. The samples were reduced by heating at 65 °C for 20 min, followed by alkylation with 10 μL of 500 mM iodoacetamide at 37 °C for 30 min. The samples were then centrifuged (16,000 x *g*, 2 min) to pellet any insoluble precipitate and probe pulse sonicated (10% power output, 1s on/1s off) once more to ensure complete resuspension, and then diluted with 300 μL of 50 mM TEAB pH 8.5 to reach a final urea concentration of 2 M. Trypsin (4 μL of 0.25 μg μL^-1^ in trypsin resuspension buffer with 25 mM CaCl_2_) was added to each sample and digested at 37 °C overnight. Digested samples were then diluted with 300 μL of enrichment buffer [50 mM TEAB pH 8.5, 150 mM NaCl, 0.2% Nonidet P-40 (IGEPAL® CA-630, Sigma-Aldrich, #I8896)] containing streptavidin-agarose beads (50 μL of 50% slurry/sample) and were rotated at r.t. for 2 h. The samples were centrifuged (2,000 x *g*, 2 min) and the entire content transferred to BioSpin columns and washed (3 x 1 mL enrichment buffer, 3 x 1 mL DPBS, 3 x 1 mL water). Enriched peptides were eluted from the beads with 300 μL of 50% acetonitrile with 0.1% formic acid and dried using a SpeedVac at 46 °C. Enriched peptides were resuspended in 100 μL EPPS buffer (200 mM, pH 8.0) with 30% acetonitrile, vortexed and water bath-sonicated for 5 min. The samples were TMT-labelled by the addition of 3 μL of 20 mg mL^-1^ (in dry acetonitrile) of corresponding TMT^10^plex tag (Thermo, #90406) for 1 h at r.t with vortexing every 30 min. TMT labelling was quenched by the addition of hydroxylamine (3 μL of 5% solution in H_2_O) and incubated for 15 min at r.t. Samples were then acidified with 5 μL formic acid, combined and dried using a SpeedVac. Samples were desalted with a Sep-Pak column Vac 1 cc (50 mg) and then fractionated into 12 fractions using an acetonitrile/NH_4_HCO_3_ (10 mM) HPLC gradient (see HPLC fractionation) and analyzed by mass spectrometry (see TMT LC-MS/MS Analysis).

#### Modifications for mammalian suspension cells

Ramos cells (12.5 mL of 3 million cells mL^-1^; seeded 3 h before treatment) were treated (37 °C, 5% CO_2_) with DMSO (0.01% v/v) or parent stereoprobe for (20 µM, 3 h), Cells were harvested by centrifugation (2000 x *g*, 3 min), washed twice with cold DPBS and stored at −80 °C. The cell pellets were resuspended in 500 µL cold lysis buffer (DPBS) with EDTA-free complete protease inhibitor cocktail. Cells were lysed by probe pulse sonication (15 pulses, 10% power output, 1s on/1s off). 500 µL of 2 mg mL^-1^ of normalized whole cells lysates (Pierce BCA protein assay) were treated with 5 μL of 10 mM IA-DTB (10 µM final, 1 h, r.t.) with vortexing every 20 min. The samples were further processed for cysteine-directed MS ABPP analysis as described above.

#### *In vitro* treatment and sample processing

Cell lysates (500 µL, 2 mg mL^-1^) of proteome were treated with 5 µL of 100x parent probe (40 µM final) for 2 h, followed 5 μL of 50 mM IA-DTB (500 µM final, 1 h, r.t., final DMSO 2% v/v) and processed for cysteine-directed MS ABPP analysis as described above.

#### Data processing

Raw files processed as described above with the following modification: A dynamic modification for IA-DTB labelling (+398.25292 Da) was included with a maximum number of two differential modifications per peptide. We included a keratin filter, accepted non-unique peptides and required at least one tryptic cleavage site. A variability metric was calculated across replicate channels as the median absolute deviation divided by the mean and expressed as a percentage.

#### Criteria for liganding in cysteine-directed ABPP

A cysteine site was considered enantioselectively liganded if the variability did not exceed 20%, and at least one of the following additional criteria were met: (1) the average IA-DTB blockade by the probe was >66.7% and >2.5-fold that of its enantiomer, and either (a) more than one other cysteine is showing <25% IA-DTB blockade or (b) this site was liganded in another cysteine-directed ABPP experiment in this study; (2) the average IA-DTB blockade by the probe was >50% and >1.5-fold that of its enantiomer, and the protein was enantioselectively liganded by protein-directed ABPP in this study. All proteins passing the initial filters for liganding were manually reviewed to remove proteins showing additional evidence of high variability. Liganded peptides were required to be unique.

### HPLC fractionation

The protein- and cysteine-directed ABPP samples were resuspended in 500 µL of Velos buffer (95% H_2_O, 5% acetonitrile, 0.1% formic acid) and fractionated with an Agilent HPLC system into a 96-deep-well plate containing 20 μL of 20% formic acid to acidify the eluting peptides, as previously reported^32^. The peptides were eluted onto a capillary column (ZORBAX 300Extend-C18, 3.5 μm) and separated at a flow rate of 0.5 mL min^−1^ using the following gradient: 100% buffer A (10 mM NH_4_HCO_3_) from 0 min to 2 min, 0–13% buffer B (HPLC-grade acetonitrile; Fisher Scientific) from 2 min to 3 min, 13–42% buffer B from 3 min to 60 min, 42–100% buffer B from 60 min to 61 min, 100% buffer B from 61 min to 65 min, 100–0% buffer B from 65 min to 66 min, 100% buffer A from 66 min to 75 min, 0–13% buffer B from 75 min to 78 min, 13–80% buffer B from 78 min to 80 min, 80% buffer B from 80 min to 85 min, 100% buffer A from 86 min to 91 min, 0–13% buffer B from 91 min to 94 min, 13–80% buffer B from 94 min to 96 min, 80% buffer B from 96 min to 101 min, and 80–0% buffer B from 101 min to 102 min. Peptides were collected in 300 μL fractions from minute 5 to 75. The plates were evaporated to dryness using a SpeedVac and peptides resuspended in 80% acetonitrile, 20% H_2_O, 0.1% formic acid, and combined to a total of 12 fractions (for example, fraction1 = well 1A + 1B … 1H, fraction 2 = well 2A + 2B …. 2H) (3 × 300 μL column^−1^). Samples were dried with a SpeedVac, and the resulting 12 fractions were resuspended in Velos buffer and analyzed by LC-MS/MS.

### TMT LC-MS/MS analysis

Fractions were resuspended in Velos buffer (95% H_2_O, 5% acetonitrile, 0.1% formic acid) and analyzed by liquid chromatography tandem mass-spectrometry using an Orbitrap Fusion Tribrid Mass Spectrometer (Thermo Scientific) coupled to an UltiMate 3000 Series Rapid Separation LC system and autosampler (Thermo Scientific Dionex). The peptides were eluted onto a capillary column (75-μm-inner-diameter fused silica, packed with C18 (Waters, Acquity BEH C18, 1.7 μm, 25 cm) or an EASY-Spray HPLC column (Thermo, #ES902, #ES903) using an Acclaim PepMap 100 (Thermo, #164535) loading column, and separated at a flow rate of 0.25 μL min^-1^. Peptides were separated across a 10 min gradient of 5%, 150 min gradient of 5-20%, 20 min 20-45%, and then 5 min 45-95% acetonitrile (0.1% formic acid) in H_2_O (0.1% formic acid) followed by column equilibration. Data were acquired using an MS3-based TMT method on Orbitrap Fusion or Eclipse Tribrid mass spectrometers.

Fusion instruments: The scan sequence began with an MS1 master scan (Orbitrap analysis, resolution 120,000, 400-1,700 m/z, RF lens 60%, maximum injection time 50 ms) with dynamic exclusion enabled (repeat count 1, duration 15 s). The top precursors were then selected for MS2/MS3 analysis. MS2 analysis consisted of quadrupole isolation (isolation window 0.7) of precursor ion followed by collision-induced dissociation in the ion trap (collision energy 35%, maximum injection time 120 ms). Following the acquisition of each MS2 spectrum, synchronous precursor selection enabled the selection of up to 10 MS2 fragment ions for MS3 analysis. MS3 precursors were fragmented by higher-energy collisional dissociation (HCD) and analyzed using the Orbitrap (collision energy 55%, maximum injection time 120 ms, resolution 50,000). For MS3 analysis, we used charge state-dependent isolation windows. For charge state z = 2, the MS isolation window was set at 1.2; for z = 3–6 the MS isolation window was set at 0.7.

Eclipse Tribrid instrument: The scan sequence began with an MS1 master scan (Orbitrap analysis, resolution 120,000, 400-1,700 m/z, RF lens 30%, maximum injection time 50 ms) with dynamic exclusion enabled (repeat count 1, duration 30 s). The top precursors were then selected for MS2/MS3 analysis. MS2 analysis consisted of quadrupole isolation (isolation window 0.7) of precursor ion followed by higher-energy collisional dissociation (HCD) in the ion trap (collision energy 35%, maximum injection time 120 ms). Following the acquisition of each MS2 spectrum, synchronous precursor selection enabled the selection of up to 10 MS2 fragment ions for MS3 analysis. MS3 precursors were fragmented by HCD and analyzed using the Orbitrap (collision energy 55%, maximum injection time 120 ms, resolution 30,000). For MS3 analysis, we used charge state-dependent isolation windows. For charge state z = 2, the MS isolation window was set at 1.2; for z = 3 the MS isolation window was set at 0.7; for z = 4–6, the MS isolation window was set at 0.4.

### Generation of HEK293T cells stably expressing WT-*Zt*GSPT1-FLAG, C441A-*Zt*GSPT1-FLAG, WT-*Hs*GSPT1-FLAG, WT-*Zt*GSPT1-NLuc, and C441A-*Zt*SAR1-NLuc

#### Cloning and mutagenesis of FLAG-modified plasmids

The full-length, attb-flanked and C-term-FLAG epitope-containing *Z. tritici* and human GSPT1 gene constructs (i.e., WT-*Zt*GSPT1-FLAG and *Hs*GSPT1-FLAG) were synthesized from Integrated DNA Technologies and cloned into pDONR221 entry vector (Thermo, #12536017), then into pLEX307 (Addgene, #41392) or pRK5-derived expression vector using Gateway technology (Thermo, #11789013 and #11791019; for ORFs see Supplementary Table 1). The plasmid pDONR221_C441A-*Zt*GSPT1-FLAG was cloned by mutagenesis using a Q5 site-directed mutagenesis kit (NEB, E0554S) with the primers described below.

C441A-*Zt*GSPT1-FLAG_FWD: GCCCTAGCAAAACTGACTC C441A-*Zt*GSPT1-FLAG_REV: TTCTTTATACCTCTCCTC

#### Cloning and mutagenesis of NLuc-modified plasmids

For generating WT-*Zt*GSPT1-NLuc and C441A-ZtGSPT1-NLuc construct, XhoI was first inserted into CDK2-NLuc vector (gift from Dr. Yuanjin Zhang) using Q5 site-directed mutagenesis kit with the primers described below.

Forward primer: CTCGTTTCTCGAGGTGGTTCAGGTG

Reverse primer: TCGAAGATGGGGTACTGG

The WT-*Zt*GSPT1 and C441A-ZtGSPT1 plasmids were then cloned into AsiSI-digested and XhoI-digested CDK2-NLuc plasmid.

Forward primer: TTTGCGATCGC ATGGCCAACGGACAG

Reverse primer: AAATAACTAAGCTTATCACGGATGCTACTCTCGAAGAAA

#### Virus production

HEK293T cells (2.0 x 10^6^ in 5 mL DMEM) were seeded into 6 cm dishes and allowed to grow for one day. psPAX2 (2.25 μg), pMD2.G (0.75 μg), and pLEX307 backbone-containing plasmids (3 μg) were mixed in 600 μL serum-free DMEM along with 20 µL polyethylenimine (PEI, 1 mg mL^-1^) and incubated for 15 min. The mixture was gently added to the cells. The virus was collected after two days and filtered with 0.45 μm syringe filters (Millipore). Virus was either used immediately for transduction or was stored at −80 °C until a later date.

#### Transduction

HEK293T cells (0.5 x 10^6^ in 2 mL DMEM/well) were seeded in a 6-well dish and allowed to grow for one day. On the next day, 1 mL of media was removed and replaced with 1 mL virus supernatant and 10 μg mL^-1^ polybrene (Millipore). On day 1 post-infection, the virus containing media was removed, and 5 mL of fresh DMEM was added. On day 2, 2 μg mL^-1^ puromycin was added to start the selection (3-5 days).

#### Transient transfection

HEK293T cells (2 x 10^6^ in 10 mL DMEM lacking penicillin/streptomycin) were seeded in 10 cm plate and allowed to grow for one day. The next day a transfection mix containing WT or C441A GSPT1-NLuc plasmids (10 µg), PEI (30 µL of 1 mg mL^-1^ stock), and 1 mL serum-free DMEM was incubated at r.t. for 15 min. After incubation, the mix was added to the cells dropwise and allowed to sit for 48 h.

### Gel-ABPP analysis of recombinant GSPT1

#### In situ treatment

HEK293T (parental control) or stably expressing GSPT1-FLAG HEK293T cells (3.0 x 10^6^ in 6 mL DMEM) were seeded into 6 cm plates the day before. Cells were treated the next day with DMSO (1% v/v) or parent stereoprobe (2–40 µM, 2 h), followed by treatment with alkyne stereoprobe (5 µM, 1 h) at 37 °C, 5% CO_2_. Cells were scraped in cold DPBS, collected by centrifugation at 500 x *g* for 5 min at 4 °C and washed twice with cold DPBS. The cell pellets were resuspended in 500 μL cold lysis buffer (DPBS) with EDTA-free complete protease inhibitor cocktail and lysed by probe pulse sonication (15 pulses, 10% power output, 1s on/1s off). 50 µL of 1 mg mL^-1^ of normalized whole cells lysates (DC protein assay) were treated with 5.5 µL of click master mix (r.t., 1 h). Afterwards, 17 µL of 4x Laemmli loading dye was added and samples were resolved by SDS-PAGE (160 V, 1 h, 4–20% tris-glycine; Invitrogen, XP04205BOX).

#### *In vitro* treatment

Cell lysates (50 µL, 1 mg mL^-1^) were treated with 1 µL of 50x alkyne probe (5 µM, 1 h, final DMSO 2% v/v) and analyzed as described above for click gel-ABPP analysis and western blot analysis (see Western Blotting below).

#### Quantification

Rhodamine band intensities were quantified using the Image Lab (v6.1.0) software (Bio-Rad).

### Western Blotting

Cells were lysed in cold DPBS by probe pulse sonication (15 pulses, 10% power output, 1s on/1s off). Cell lysates (50 µL) were normalized to 1 mg mL^-1^ (DC protein assay), followed by the addition of 4x Laemmli gel loading buffer (36 μL). Proteins were resolved using SDS-PAGE gel analysis (160 V, 1 h, 4–20% tris-glycine). Proteins were then transferred onto nitrocellulose membrane (semi-dry, 25 V, 10 min), blocked with 3% bovine serum albumin (BSA) in tris-buffered saline with 0.1% Tween-20 (TBST) for 60 min at r.t., incubated with primary antibodies (see Supplementary Table 2 for dilution) either at r.t. for 2 h or at 4 °C overnight. Membranes were washed with TBST (5 times) before treatment with secondary antibodies at r.t. for 2 h or directly imaged if a primary HRP-conjugate was used. Membranes were developed with SuperSignal™ West Femto PLUS Chemiluminescent Substrate and visualized by chemiluminescence scan on a ChemiDoc MP Imaging System (Bio-Rad).

#### Quantification

Band intensities were quantified using the Image Lab software (v6.1.0). Quantified immunoblot signals were calculated as a ratio of the band intensity of the recombinant tag epitope (e.g., FLAG or V5) over the relevant housekeeping protein (e.g., Na^+^/K^+^-ATPase or β-Actin). IC_50_ curves were generated using GraphPad Prism (v10.4.1), applying a four-parameter variable slope nonlinear regression.

### GSPT1 NanoBRET assay

NanoLuc-tagged protein of interest (WT-*Zt*GSPT1-NLuc or C441A-*Zt*GSPT1-NLuc) was recombinantly expressed in HEK293T cells by transient transfection following the above mentioned transfection protocol for epitope-tagged proteins. Cells were harvested by scraping 48 h after transfection and lysed in BRET assay buffer (50 mM HEPES pH 7.5 and 150 mM NaCl) by probe sonication (2 x 15 pulses), and the protein concentration was adjusted to 2 mg mL^-1^. The lysate was mixed with vivazine (Promega, 1% v/v), CJR-88A or B (final DMSO 1% v/v), and competitors for the competition assay, then dispensed to 384-well plates (Greiner, 784075) and allowed to incubate at room temperature. The BRET signal was read out on CLARIOstar microplate reader (BMG LABTECH) equipped with a 450 nm BP filter (donor) and a 610 nm LP filter (acceptor). The BRET ratio of each sample was calculated as follows: BRET ratio = (Acceptor_sample_/Donor_sample_) – (Acceptor_control_/Donor_control_), where sample = signal from lysates from the HEK293T cells transfected with WT-*Zt*GSPT1-NLuc, control = signal from lysates from the HEK293T cells transfected with C441A-*Zt*GSPT1-NLuc.

### Generation of HEK293T cells stably expressing WT-*Zt*SAR1-FLAG, C64A-*Zt*SAR1-FLAG, WT-*Bc*SAR1-FLAG, V64C-*Bc*SAR1-FLAG, WT-*Pg*SAR1-FLAG, V64C-*Pg*SAR1-FLAG, WT-*Hs*SAR1A-V5 or M69C-*Hs*SAR1A-V5

#### Cloning and mutagenesis

The full-length, attb-flanked and cterm-FLAG-STOP epitope-containing *Z. tritici*, *B. cinerea*, and *P. graminis* SAR1 gene constructs (i.e., WT-*Zt*SAR1-FLAG, WT-*Bc*SAR1-FLAG, V64C-*Bc*SAR1-FLAG, WT-*Pg*SAR1-FLAG, V64C-*Pg*SAR1-FLAG) were synthesized from Twist Biosciences and cloned first into a pDONR221 (Thermo, #12536017) entry vector. The full-length, attl-flanked human SAR1A (Q5SQT9) gene construct (WT-*Hs*SAR1A) inserted into a pDONR223 entry vector were synthesized from Dharmacon Inc (hORFeome v8.1 library). Both pDONR221 and pDONR223 entry vectors were cloned into the pLEX307 destination (Addgene, #136403) expression vector using Gateway technology (for ORFs see Supplementary Table 1).

The plasmid pLEX307_C64A-*Zt*SAR1-FLAG was cloned by mutagenesis using a Q5 site-directed mutagenesis kit (NEB, E0554S) with the primers described below.

C64A-*Zt*SAR1-FLAG_FWD: GCTAGGTTCACTACCTTCGAC C64A-*Zt*SAR1-FLAG_REV: GCTTCCTATGGATAGTTC

The plasmid pLEX307_M69C-*Hs*SAR1A-V5 was cloned by mutagenesis using a Q5 site-directed mutagenesis kit with the primers described below.

M69C-*Hs*SAR1A-V5_FWD: ATTGCTGGATGCACCTTTACAACT M69C-*Hs*SAR1A-V5_REV: TGTTAGCTCTTCTGA

#### Virus production

Virus was prepared as described above.

#### Transduction

HEK293T cells were transduced as described above.

### FLAG/V5-IP Gel-ABPP analysis of recombinant SAR1

#### *In situ* treatment

HEK293T (parental control) or stably expressing SAR1-FLAG/V5 HEK293T cells (5.0 x 10^6^ in 6 mL DMEM) were seeded into 6 cm plates the day before. Cells were treated the next day with DMSO (1% v/v) or parent stereoprobe (0.01–40 µM, 0.5–4 h), followed by treatment with alkyne stereoprobe (5 µM, 1 h) at 37 °C, 5% CO_2_. Cells were scraped in cold DPBS, collected by centrifugation at 500 x *g* for 5 min at 4 °C and washed twice with cold DPBS. The cell pellets were resuspended in 250 μL cold lysis buffer (DPBS, 150 mM NaCl, and 0.1% Triton X-100) with EDTA-free complete protease inhibitor cocktail and lysed by probe pulse sonication (15 pulses, 10% power output, 1s on/1s off). The lysate was separated with 150 µL being centrifugated at 100,000 x *g* for 45 min at 4 °C to separate soluble and particulate fractions; the remaining 100 µL that was not centrifugated at this step was the ‘whole cells’. After centrifugation, the supernatant was separated (‘soluble’ fraction) and the pellet was resuspended in 100 µL of lysis buffer via probe pulse sonication (15 pulses, 10% power output, 1s on/1s off); ‘particulate fraction’.

#### For the input

30 µL of 1 mg mL^-1^ of normalized lysate fractions (DC protein assay) were treated with 3 μL of click master-mix (r.t., 1 h). Afterwards, 10 µL of 4x Laemmli loading dye was added and samples were resolved by SDS-PAGE (160 V, 1 h, 4–20% tris-glycine). Samples were processed for Western blot analysis as described above.

#### For the IP

300 μL of 1 mg mL^-1^ of normalized lysate fractions (DC protein assay) were incubated with washed Pierce™ Anti-DYKDDDDK Magnetic Agarose (Thermo, #A36797; 15 μL of 25% slurry/sample) or washed Sigma-Aldrich Anti-V5 Magnetic Beads (Sigma-Aldrich, SAE0203; 15 μL/sample) for 2 h at r.t. with rotation. After incubation, the beads were isolated with a magnetic stand and washed with 0.5 % Nonidet P-40 in DPBS (3 x 0.5 mL). The beads were resuspended in 30 μL DPBS and treated with 3 μL of click Master-mix (r.t., 1 h). Afterwards, 10 µL of 4x Laemmli loading dye was added and samples were resolved by SDS-PAGE (160 V, 1 h, 4–20% tris-glycine). Samples were processed for Western blot analysis as described above.

#### *In vitro* treatment

Cell lysates (50 µL, 1 mg mL^-1^) were treated with 1 µL of 50x alkyne stereoprobe (5 µM final, 1 h) at r.t., followed by 5.5 µL of click master mix (r.t., 1 h). Afterwards, 17 µL of 4x Laemmli loading dye was added and samples were resolved by SDS-PAGE (160 V, 1 h, 4–20% tris-glycine). Samples were processed for Western blot analysis as described above.

#### Quantification

Rhodamine and immunoblot band intensities were calculated as described above.

### FLAG-IP Gel-ABPP analysis of recombinant SAR1 with guanosine nucleotide treatment

#### In vitro treatment

HEK293T (parental control) or stably expressing SAR1-FLAG HEK293T cells (5.0 x 10^6^ in 6 mL DMEM) were seeded into 6 cm plates the day before. Cells were scraped in cold DPBS, collected by centrifugation at 500 x *g* for 5 min at 4 °C and washed twice with cold DPBS. The cell pellets were resuspended in 300 μL cold lysis buffer (40 mM HEPES pH 7.5, 200 mM NaCl, and 5 mM MgCl_2_) with EDTA-free complete protease inhibitor cocktail and lysed by probe pulse sonication (15 pulses, 10% power output, 1s on/1s off). The lysate was centrifugated at 100,000 x *g* for 45 min at 4 °C and the particulate fraction was prepared as described above. 50 µL of 1 mg mL^-1^ of normalized (DC protein assay) particulate lysates were incubated with either GDP (Sigma-Aldrich, G7127) or GppCp (Abcam, ab146660) dissolved in lysis buffer for 1 h on ice. The samples were then treated with 1 µL of 50x alkyne stereoprobe (5 µM, 1 h, final DMSO 2% v/v) at r.t. Afterwards, 17 µL of 4x Laemmli loading dye was added and samples were resolved by SDS-PAGE (160 V, 1 h, 4–20% tris-glycine). Samples were processed for western blot analysis as described above.

#### Quantification

Rhodamine and immunoblot band intensities were calculated as described above.

### Generation of HEK293T cells stably expressing WT-*Zt*SAR1-eGFP, C64A-*Zt*SAR1-eGFP, WT-*Bc*SAR1-eGFP, V64C-*Bc*SAR1-eGFP, WT-*Pg*SAR1-eGFP, V64C-*Pg*SAR1-eGFP, WT-*Hs*SAR1A-eGFP or M69C-*Hs*SAR1A-eGFP

#### Cloning and mutagenesis

The full-length, attb-flanked and cterm-eGFP-STOP epitope-containing *Z. tritici*, *B. cinerea*, *P. graminis* SAR1, and *H. sapiens* SAR1A gene constructs (i.e., WT-*Zt*SAR1-eGFP, C64A-*Zt*SAR1-eGFP, WT-*Bc*SAR1-eGFP, V64C-*Bc*SAR1-eGFP, WT-*Pg*SAR1-eGFP, V64C-*Pg*SAR1-eGFP, WT-*Hs*SAR1A-eGFP, and M69C-*Hs*SAR1A-eGFP) were synthesized from Twist Biosciences and cloned first into a pDONR221 entry vector then into a pLEX307 expression vector using Gateway technology (for ORFs see Supplementary Table 1). The stable cell lines were generated as described above.

#### Virus production

Virus was prepared as described above.

#### Transduction

HEK293T cells were transduced as described above.

### Generation of HEK293T cells co-expressing WT-*Zt*SAR1-eGFP and Halo-*Hs*COPII-related proteins

#### Cloning and mutagenesis

The full-length, attl-flanked human SAR1A (Q5SQT9), SEC23B (B4DJW8), SEC31A (A0A024RDD3), STX5 (B4DKR0), TFG (Q05BK6), TMED2 (Q15363), and TRAPPC1 (Q9Y5R8) gene construct inserted into pDONR223 entry vectors were synthesized from Dharmacon Inc (hORFeome v8.1 library) and cloned into a pHTNW destination (Addgene, #136403) expression vector using Gateway technology (for ORFs see Supplementary Table 1).

#### Transient transfection

WT-*Zt*SAR1-eGFP HEK293T cells (0.5 x 10^6^ in 2 mL DMEM lacking penicillin/streptomycin) were seeded in 27 mm glass bottom dishes pretreated with poly-D-lysine (1 mL of 100 µg mL^-1^ at r.t. for 1 h; Thermo, A3890401) overnight. The next day a transfection mix containing Halo-*Hs*COPII plasmids (2 µg), PEI (10 µL of 1 mg mL^-1^ stock), and 200 µL serum-free DMEM was incubated at r.t. for 15 min. After incubation, the mix was added to the cells dropwise and allowed to sit for 48 h.

### Live-cell immunofluorescence using confocal microscopy

#### Cell culture

27 mm glass bottom dishes were coated with poly-D-lysine (1 mL of 100 µg mL^-1^; Gibco, A3890401) at r.t. for 1 h and then washed twice with DPBS. SAR1-eGFP expressing HEK293T cell lines were seeded (0.5 x 10^6^ in 2 mL DMEM lacking phenol red; Thermo, 21063029) in poly-D-lysine-treated glass bottom dishes. Cells were stained for the ER membrane using ER-Tracker^TM^ Blue-White DPX (1 µM for 30 min; Invitrogen, E12353) and had the media replaced with fresh DMEM lacking phenol red in preparation for live-cell confocal microscopy imaging. Conversely, SAR1-eGFP expressing *Z. tritici* cells were seeded (1 x 10^6^ cells mL^-1^ in 2 mL PDB media) into untreated 27 mm glass bottom dishes prior to immunofluorescence imaging.

#### Imaging

Live cell imaging was performed using an Abberior Facility Line 3D STED microscope equipped with 5 lasers for fluor excitation (405, 440, 485, 561, & 640 nm), an Olympus IX83 microscope body, and spectral detection (3 APD detectors and 1 Matrix detector) using the 60x oil objective lens (NA 1.42). For monitoring SAR1-eGFP fluorescence aggregation, DMSO (1% v/v) or parent stereoprobe (MY1A, MY1B; 5–40 µM, 2 h) was added to HEK293T or *Z. tritici* cells expressing GFP-conjugated SAR1. For monitoring Halo-COPII and SAR1-eGFP fluorescence colocalization, JFX650 HaloTag^®^ ligand resuspended in DMEM (200 nM final; Promega, HT1070) was added to cells for 30 min, then replaced with phenol red-free media prior to imaging. In HEK293T cells, SAR1-eGFP fluorescence (excitation λ = 488 nm), Halo-COPII fluorescence (excitation λ = 640 nm), and ER membrane fluorescence (excitation λ = 405 nm) were detected under confocal acquisition. In *Z. tritici* cells, SAR1-eGFP fluorescence (excitation λ = 488 nm) and ER membrane fluorescence (excitation λ = 561 nm) were detected under confocal acquisition. The cell stage in the Abberior Facility Line 3D STED microscope was prepared beforehand (chamber held at 37 °C, 5% CO_2_) to accommodate live cell imaging for mammalian cell samples. Imaging of *Z. tritici* cell samples was performed at ambient conditions.

#### Data analysis

SAR1-eGFP fluorescence aggregation was measured using the CellProfiler software (v4.2.6) to both identify and quantify cells and puncta. In HEK293T cells, cell objects were predicted based on an expected diameter of 30–200 pixel units using an Otsu threshold correction method to remove background fluorescence with declumping of objects based on shape. Puncta objects were predicted based on an expected diameter of 6–30 pixel units using an Otsu threshold correction method to remove background fluorescence. In *Z. tritici* cells, cell objects were predicted based on an expected diameter of 10–500 pixel units using an Otsu threshold correction method to remove background fluorescence with declumping of objects based on shape. Puncta objects were predicted based on an expected diameter of 3–20 pixel units using an Otsu threshold correction method to remove background fluorescence with declumping of objects based on intensity. Whole number cell and puncta counts were reported for each provided microscopy image and used to calculate the ratio of puncta-to-cells for each treatment condition.

SAR1-eGFP and Halo-COPII fluorescence colocalization was performed using the ImageJ software (Fiji) Coloc 2 plugin to measure the Pearson’s R value under a Costes threshold regression of both the eGFP (488 nm) and HaloTag (640 nm) channels for each microscopy image. The average of these values was used to calculate the colocalization rates of DMSO- or MY-1A-treated cells.

### Generation of *Z. tritici* cells stably expressing epitope-tagged proteins of interest (ATMT)

#### Cloning and mutagenesis

Full gene replacement constructs for *in locus* gene swap by homologous recombination were designed following the dual markers flanking strategy as previously described^11^. The full insert including 1kb of the genomic upstream and downstream flanking regions of *Zt*SAR1, selection cassettes (Hygromycin B and Isofetamid), TetOff promoter^75^, *Zt*SAR1-WT-GFP or its *Zt*SAR1-C64A-GFP mutant were synthesized by GENEWIZ Inc. (Germany) and assembled into the binary plasmid pNOV2114^108^.

#### *Agrobacterium tumefaciens* mediated transformations - ATMT

*A. tumefaciens-*mediated transformations of the *Z. tritici* IPO323-ΔKU70::TtA* strain^11^ were performed following already described procedures^108^ using the *A. tumefaciens* AGL1 strain. ATMT induction media (IM)-agar plates were prepared containing 50 µg mL^-1^ spectinomycin and 40 µg mL^-1^ acetosyringone. They were incubated at 19 °C for 2 days before filters were transferred onto selective AE media plates (*Aspergillus* Essential medium) containing 250 µg mL^-1^ cefotaxime (Sigma), 100 µg mL^-1^ ampicillin (Sigma), 100 µg mL^-1^ streptomycin (Sigma), 200 µg mL^-1^ hygromycin B (Invitrogen), and 10 µg mL^-1^ isofetamid (Angene). Plates were incubated at 19 °C for 7–10 days. Positive colonies were confirmed by PCR and Western blot using anti-GFP antibodies (Abcam, ab6556)

### Sensitivity assay of *Z. tritici* SAR1 cell lines

*Z. tritici* SAR1-WT and SAR1-C64A cells were collected from 6 days-old YPD plates and resuspended in liquid minimal media for *Z. tritici* (MM-Zt + NO_3_)^109^. Cell concentration was adjusted to 5×10^5^ cells mL^-1^ and distributed into 96-well plates (Corning Costar 3370), 198 µL/well (10^5^ cells/well). A 1:3 dilution series for the stereoprobes MY1A and MY1B was prepared starting from 10 mM in DMSO and 2 µL were transferred to the testing plate (final highest concentration 100 µM). Plates were incubated at 19 °C for 7 days. OD_600_ nm was measured using an EnVision plate reader (PerkinElmer Life Sciences). Nonlinear regression was performed using log(inhibitor) vs. response to calculate IC_50_ values using GraphPad Prism.

### Structural Modeling

Non-conserved residues near *Zt*GSPT1_C441 between *Zt*GSPT1 (AF-F9X8V1-F1) and *Hs*GSPT1 (PDB ID: 3J5Y) were highlighted in ChimeraX (v1.10). Structural alignment between *Hs*NMT1 bound to IMP-1088 (PDB: 5MU6) and *Zt*NMT1 (AF-F9XBB7-F1) was performed using the ‘Match’ command in ChimeraX (v1.10). Model of the *Zt*NMT1 structure in complex with myristoyl-CoA cofactor were generated using the ‘Boltz’ structure prediction tool available in ChimeraX. The amino acid sequence for *Zt*NMT1 was retrieved from Uniprot (ID: F9XBB7). Models of *Zt*SAR1 structures (AF-F9X159-F1) in complex with GDP and GTP were generated using Boltz-2^58,110^ (v2.1.1, GPU: RTXA6000). The amino acid sequence for *Zt*SAR1 was retrieved from Uniprot (ID: F9X159) and the co-factors were specified using the Cambridge Chemical Dictionary (CCD 3-letter codes: GDP and GTP).

### Statistics and reproducibility

Statistical analyses and data visualization in this paper were performed using GraphPad Prism 10 (v10.4.2). To compare group means, we performed two-tailed unpaired *t*-tests between group pairs. For experiments comparing multiple treatments to a single treatment group, we used one-way ANOVA followed by Tukey’s post hoc test; reported *p* values are adjusted p values. For experiments comparing a single treatment across multiple treatment groups, we used two-way ANOVA followed by Šídák post hoc test; reported *p* values are adjusted *p* values

## Data availability

The mass spectrometry proteomics data have been deposited to the ProteomeXchange Consortium via the PRIDE81 partner repository with the dataset identifier PXD069812. Processed proteomic data are provided in Supplementary Dataset 1. Data associated with the paper can be found in Supplementary Information. Source data are provided with this paper. All human cell models and parental (i.e., non-modified) fungal cell samples (IPO323 strain) generated and analyzed during the current study are available from the corresponding authors upon reasonable request. Only genetically-modified fungal cell samples are unavailable for distribution, however, the description of their generation, propagation, and analysis are described in the Methods section.

