## Supplementary Information for "Mapping ortholog-restricted ligandable cysteines in the wheat pathogen *Zymoseptoria tritici*"

**Supplementary Table 1.** ORF sequences of gene constructs used in this study.

| Gene Name | ORF sequence |
|---|---|
| **WT-*Zt*GSPT1-FLAG** | ATGGCCAACGGACAGCAACCTGAGAGCTGGGAAGACGAAATGAGGGAT<br>GAGGACTTGGCTAATCAAACCCAACAGCAGATGAATATGCAGGGCGGGG<br>CGCCTAGGGGAGGGACATTCGTTCCTGGAGCCCAGAGCTTCACTCCAG<br>GAGCGCAATCCTTCAATCCTGGTCAGCAGTACCAGCAATATGGATACGG<br>GCAGCAGTATGGGGGAGGATATCAGCAGTACGGTCAGCAGCAAGGCTA<br>CAACCAGTACCAGCAATATGGCCAATATCAACAACAGGGGTACGGTCAG<br>CAGCAATATGCACAACAAGGTTACAACCAGTACCAGCAACCCCAACAGC<br>AAGCATATGTACCCCCAGCCGCCCGTCAGGCCCCGATGATAGCCAAGAG<br>GGGAGAGGCCCTTCCTACCGCCGCCGCTGCGAAACCAGCTGCTGCCA<br>ATGGCGCAGCCGCCCCCGTTAAGACACTGTCATTGAGTGCTGCAGATAC<br>TGGGGCAGCCGCGCCTAAGGTGGAGAAAGCGACCGGTGGAGCAAAGG<br>TGTTGTCCCTAGGTGGTGCACCGGCTGCTGCAACGCCGAAGGCTGCAC<br>CCGCCCCTAAAGAGGCAGTGAGCAAGGACGCGCCAGAGGCCGGCACT<br>AAGGTCGCTGCCGCCAAAGCTATCGAGAAAACCGGGGAACCGACGGCT<br>GCAGTATCTGGAGCTAGCTCTCCCACACCCAGTGGCAGATCTTCTCCCA<br>CAAGAGCAGACATAAAAGCGGAGGAGAAACGTAAAGCCGACCAGGTGC<br>TAGCCGACCAAGCCGCAGAAGTAGATGAAGAAGCGCTAGCCGAGATGTA<br>TGGTAAGGAGCACGTCAACGTTATCTTTCTTGGCCATGTGGATGCAGGC<br>AAGTCTACCTTAGGTGGGAGCATTCTTTATGCGACTGGGATGGTTGACG<br>AACGTACCATGGATAAATATAAGCGTGATGCTAAGGATATGGGAAGGGAA<br>TCATGGTACTTGTCATGGGCTCTAGACCAGACTAAGGAGGAGAGAGCCC<br>AGGGTAAGACCGTGGAGGTCGGGCGTGGCTTCTTCGAAACGGAAAAGA<br>GGCGTTACTCCATACTAGACGCCCCTGGGCATAAGACTTTCGTACCTAAC<br>ATGCTATCAGGCGCGTCACAGGCAGACGTCGGGGTACTAGTCATCTCCG<br>CTAGGAAAGGAGAGTATGAAACAGGTTTTGAGAAAGGCGGCCAGACGA<br>GGGAGCATGCAGTTTTGGCCAAGACGCAGGGGATTAACAAACTGATAGT<br>AGTTGTGAATAAGATGGACGACATTACGGTCGAGTGGTCCGAGGAGAG<br>GTATAAAGAATGCCTAGCAAAACTGACTCAGTTCTTAAAAGGATTGGGAT<br>ATAACCCTAAGACAGATCTAACTTTCATGCCAGTAGCGGCCCAGCAAACG<br>ATGGGGATCAAAGACAGAGTGCCGAAGGACTTATGCCCCTGGTATGATG<br>GACCGAGCCTATTGGAATTCCTTGATTCCATGCAAGCATTGGAGCGTAAG<br>TTATCCGCCCCATTCATGATGCCCATCAGTGCCAAATATAGAGATATGGGA<br>ACAATGATCGAGGGGAAAATCGAAGCTGGTTTTATAAAGAAAGACCAAAA<br>ATATCTAATGATGCCCAATAAAGCGGAAATACAAATATCTGCACTTTATGGT<br>GAGTCTGAAGAAGAGATCGCCCACGCCACCTCAGGGGAACAAATCCGT<br>TTGCGTATTAAGGGGGCCGAGGAAGAGGACATTTACCCGGGGTTCGTCT<br>TATGTTCCCCCAAACGTCCCGTCCATTGCGTGACAACTTTCGAGGCGCA<br>GATCAGACTACTTGAATTGAAATCAATTTTAAGCGCGGGTTTCAACTGCG<br>TTATACACGTGCATTCCGCTACTGAGGAGGTCACGTTTACAGAGCTGTTG<br>CACAAACTGGAACCGAAGACAAATAGGAAATCCAAGAAGCCACCGGGTT<br>TCGCGAAACAGGGTATGAATATCATTGCGAGGCTGGAAGTTACGGACA |

| | |
|---|---|
| | AGCGGGTAGCCTGTGTGTCGAGAAGTTCGAGGATTACCCTCAACTTGGA<br>AGGTTCACACTAAGAGATCAAGGGCAGACTATCGCAATAGGTAAAATAAC<br>TAAGCTTATCACGGATGCTACTGGCGGCTCTTCTGACTACAAGGATGAC<br>GATGACAAGTGA |
| **C441A-*Zt*GSPT1-FLAG** | ATGGCCAACGGACAGCAACCTGAGAGCTGGGAAGACGAAATGAGGGAT<br>GAGGACTTGGCTAATCAAACCCAACAGCAGATGAATATGCAGGGCGGGG<br>CGCCTAGGGGAGGGACATTCGTTCCTGGAGCCCAGAGCTTCACTCCAG<br>GAGCGCAATCCTTCAATCCTGGTCAGCAGTACCAGCAATATGGATACGG<br>GCAGCAGTATGGGGGAGGATATCAGCAGTACGGTCAGCAGCAAGGCTA<br>CAACCAGTACCAGCAATATGGCCAATATCAACAACAGGGGTACGGTCAG<br>CAGCAATATGCACAACAAGGTTACAACCAGTACCAGCAACCCCAACAGC<br>AAGCATATGTACCCCCAGCCGCCCGTCAGGCCCCGATGATAGCCAAGAG<br>GGGAGAGGCCCTTCCTACCGCCGCCGCTGCGAAACCAGCTGCTGCCA<br>ATGGCGCAGCCGCCCCCGTTAAGACACTGTCATTGAGTGCTGCAGATAC<br>TGGGGCAGCCGCGCCTAAGGTGGAGAAAGCGACCGGTGGAGCAAAGG<br>TGTTGTCCCTAGGTGGTGCACCGGCTGCTGCAACGCCGAAGGCTGCAC<br>CCGCCCCTAAAGAGGCAGTGAGCAAGGACGCGCCAGAGGCCGGCACT<br>AAGGTCGCTGCCGCCAAAGCTATCGAGAAAACCGGGGAACCGACGGCT<br>GCAGTATCTGGAGCTAGCTCTCCCACACCCAGTGGCAGATCTTCTCCCA<br>CAAGAGCAGACATAAAAGCGGAGGAGAAACGTAAAGCCGACCAGGTGC<br>TAGCCGACCAAGCCGCAGAAGTAGATGAAGAAGCGCTAGCCGAGATGTA<br>TGGTAAGGAGCACGTCAACGTTATCTTTCTTGGCCATGTGGATGCAGGC<br>AAGTCTACCTTAGGTGGGAGCATTCTTTATGCGACTGGGATGGTTGACG<br>AACGTACCATGGATAAATATAAGCGTGATGCTAAGGATATGGGAAGGGAA<br>TCATGGTACTTGTCATGGGCTCTAGACCAGACTAAGGAGGAGAGAGCCC<br>AGGGTAAGACCGTGGAGGTCGGGCGTGGCTTCTTCGAAACGGAAAAGA<br>GGCGTTACTCCATACTAGACGCCCCTGGGCATAAGACTTTCGTACCTAAC<br>ATGCTATCAGGCGCGTCACAGGCAGACGTCGGGGTACTAGTCATCTCCG<br>CTAGGAAAGGAGAGTATGAAACAGGTTTTGAGAAAGGCGGCCAGACGA<br>GGGAGCATGCAGTTTTGGCCAAGACGCAGGGGATTAACAAACTGATAGT<br>AGTTGTGAATAAGATGGACGACATTACGGTCGAGTGGTCCGAGGAGAG<br>GTATAAAGAAGCCCTAGCAAAACTGACTCAGTTCTTAAAAGGATTGGGAT<br>ATAACCCTAAGACAGATCTAACTTTCATGCCAGTAGCGGCCCAGCAAACG<br>ATGGGGATCAAAGACAGAGTGCCGAAGGACTTATGCCCCTGGTATGATG<br>GACCGAGCCTATTGGAATTCCTTGATTCCATGCAAGCATTGGAGCGTAAG<br>TTATCCGCCCCATTCATGATGCCCATCAGTGCCAAATATAGAGATATGGGA<br>ACAATGATCGAGGGGAAAATCGAAGCTGGTTTTATAAAGAAAGACCAAAA<br>ATATCTAATGATGCCCAATAAAGCGGAAATACAAATATCTGCACTTTATGGT<br>GAGTCTGAAGAAGAGATCGCCCACGCCACCTCAGGGGAACAAATCCGT<br>TTGCGTATTAAGGGGGCCGAGGAAGAGGACATTTACCCGGGGTTCGTCT<br>TATGTTCCCCCAAACGTCCCGTCCATTGCGTGACAACTTTCGAGGCGCA<br>GATCAGACTACTTGAATTGAAATCAATTTTAAGCGCGGGTTTCAACTGCG<br>TTATACACGTGCATTCCGCTACTGAGGAGGTCACGTTTACAGAGCTGTTG |

| | |
|---|---|
| | CACAAACTGGAACCGAAGACAAATAGGAAATCCAAGAAGCCACCGGGTT<br>TCGCGAAACAGGGTATGAATATCATTGCGAGGCTGGAAGTTACGGACA<br>AGCGGGTAGCCTGTGTGTCGAGAAGTTCGAGGATTACCCTCAACTTGGA<br>AGGTTCACACTAAGAGATCAAGGGCAGACTATCGCAATAGGTAAAATAAC<br>TAAGCTTATCACGGATGCTACTGGCGGCTCTTCTGACTACAAGGATGAC<br>GATGACAAGTGA |
| **WT-*Hs*GSPT1-FLAG** | ATGGATCCTGGGTCTGGAGGTGGTGGCGGCGGTGGTGGTGGTGGAGG<br>TTCTAGTAGTGGCTCCAGTAGCTCCGACTCCGCTCCGGACTGTTGGGA<br>TCAGGCTGACATGGAGGCTCCTGGCCCTGGTCCCTGCGGTGGTGGAG<br>GATCACTCGCGGCGGCCGCAGAGGCGCAAAGAGAAAACCTGAGCGCA<br>GCATTTAGTCGGCAGTTGAATGTTAACGCAAAACCATTTGTTCCGAATGT<br>GCATGCGGCCGAGTTTGTCCCGAGTTTCCTCAGAGGACCGGCCGCGC<br>CTCCTCCACCAGTAGGCGGCGCGGCGAATAATCACGGCGCCGGGAGC<br>GGGGCTGGAGGCAGGGCGGCTCCCGTCGAATCAAGTCAGGAAGAGCA<br>GTCACTTTGCGAGGGCTCCAATAGTGCGGTATCCATGGAACTGTCAGAA<br>CCGATCGTCGAAAATGGTGAAACTGAAATGTCCCCGGAAGAGTCATGG<br>GAGCATAAAGAGGAAATTAGCGAAGCAGAGCCTGGAGGGGGCTCTCTG<br>GGAGATGGCCGACCCCCTGAAGAATCTGCACACGAAATGATGGAGGAA<br>GAGGAAGAAATCCCAAAGCCGAAAAGCGTAGTGGCCCCACCCGGTGCA<br>CCGAAGAAAGAACATGTAAACGTAGTATTTATAGGACACGTCGATGCGG<br>GTAAGAGCACGATCGGAGGGCAAATCATGTATCTCACAGGAATGGTCG<br>ATAAACGAACGCTGGAAAAGTACGAAAGAGAGGCAAAAGAAAAGAATAG<br>GGAGACATGGTATCTGTCTTGGGCTTTGGATACAAACCAGGAAGAACG<br>CGATAAAGGAAAGACGGTAGAGGTCGGGAGGGCCTACTTCGAGACAGA<br>GAAAAAACACTTCACGATTTTGGACGCTCCTGGACACAAATCCTTTGTC<br>CCTAATATGATAGGCGGAGCATCTCAGGCGGACCTTGCTGTGCTCGTC<br>ATCTCCGCCAGAAAAGGGGAATTCGAGACCGGCTTCGAGAAAGGGGGA<br>CAAACTCGAGAGCACGCTATGTTGGCTAAAACTGCGGGCGTTAAACAC<br>CTTATCGTGTTGATCAACAAAATGGACGATCCGACGGTCAACTGGTCTA<br>ACGAAAGATATGAGGAGTGTAAAGAGAAATTGGTACCGTTTCTGAAGAA<br>GGTTGGATTCAATCCCAAGAAGGATATCCATTTTATGCCCTGCTCCGGG<br>CTCACTGGAGCAAATCTCAAAGAGCAGTCCGACTTTTGCCCCTGGTATA<br>TTGGATTGCCGTTTATACCTTACTTGGACAACCTCCCTAACTTCAACCGG<br>TCCGTTGACGGTCCGATTAGGCTTCCCATTGTAGACAAATATAAAGACA<br>TGGGCACTGTTGTCTTGGGGAAGCTTGAAAGTGGAAGCATATGCAAGG<br>GTCAGCAATTGGTGATGATGCCGAACAAACATAACGTCGAGGTGCTCG<br>GCATCTTGAGTGATGATGTAGAAACGGATACGGTAGCACCCGGGGAGA<br>ACCTTAAGATCCGACTGAAGGGTATTGAAGAGGAGGAAATCCTTCCAGG<br>TTTCATCCTGTGCGACCCTAACAACCTGTGTCACAGCGGTAGAACCTTC<br>GATGCCCAAATCGTAATTATAGAACATAAGAGCATCATTTGTCCGGGCT<br>ACAATGCTGTATTGCACATACACACCTGTATAGAAGAAGTTGAAATTACA<br>GCCCTGATCTGCTTGGTGGATAAGAAGTCCGGCGAGAAAAGTAAGACG<br>AGACCGCGATTTGTCAAACAGGACCAGGTATGTATAGCTAGACTGCGG<br>ACTGCCGGAACTATATGCTTGGAAACATTTAAGGATTTTCCCCAGATGG<br>GGCGCTTTACGCTTCGCGATGAGGGTAAAACGATTGCCATCGGTAAGG<br>TCTTGAAACTTGTTCCAGAGAAGGACGGCGGCTCTTCTGACTACAAGGA<br>TGACGATGACAAGTGA |

| | |
|---|---|
| **WT-*Zt*GSPT1-NLuc** | ATGGCCAACGGACAGCAACCTGAGAGCTGGGAAGACGAAATGAGGGATGA<br>GGACTTGGCTAATCAAACCCAACAGCAGATGAATATGCAGGGCGGGGCGC<br>CTAGGGGAGGGACATTCGTTCCTGGAGCCCAGAGCTTCACTCCAGGAGCG<br>CAATCCTTCAATCCTGGTCAGCAGTACCAGCAATATGGATACGGGCAGCAG<br>TATGGGGGAGGATATCAGCAGTACGGTCAGCAGCAAGGCTACAACCAGTA<br>CCAGCAATATGGCCAATATCAACAACAGGGGTACGGTCAGCAGCAATATGC<br>ACAACAAGGTTACAACCAGTACCAGCAACCCCAACAGCAAGCATATGTACC<br>CCCAGCCGCCCGTCAGGCCCCGATGATAGCCAAGAGGGGAGAGGCCCT<br>TCCTACCGCCGCCGCTGCGAAACCAGCTGCTGCCAATGGCGCAGCCGC<br>CCCCGTTAAGACACTGTCATTGAGTGCTGCAGATACTGGGGCAGCCGCGC<br>CTAAGGTGGAGAAAGCGACCGGTGGAGCAAAGGTGTTGTCCCTAGGTGGT<br>GCACCGGCTGCTGCAACGCCGAAGGCTGCACCCGCCCCTAAAGAGGCA<br>GTGAGCAAGGACGCGCCAGAGGCCGGCACTAAGGTCGCTGCCGCCAAA<br>GCTATCGAGAAAACCGGGGAACCGACGGCTGCAGTATCTGGAGCTAGCTC<br>TCCCACACCCAGTGGCAGATCTTCTCCCACAAGAGCAGACATAAAAGCGG<br>AGGAGAAACGTAAAGCCGACCAGGTGCTAGCCGACCAAGCCGCAGAAGT<br>AGATGAAGAAGCGCTAGCCGAGATGTATGGTAAGGAGCACGTCAACGTTAT<br>CTTTCTTGGCCATGTGGATGCAGGCAAGTCTACCTTAGGTGGGAGCATTCTT<br>TATGCGACTGGGATGGTTGACGAACGTACCATGGATAAATATAAGCGTGATG<br>CTAAGGATATGGGAAGGGAATCATGGTACTTGTCATGGGCTCTAGACCAGAC<br>TAAGGAGGAGAGAGCCCAGGGTAAGACCGTGGAGGTCGGGCGTGGCTTC<br>TTCGAAACGGAAAAGAGGCGTTACTCCATACTAGACGCCCCTGGGCATAAG<br>ACTTTCGTACCTAACATGCTATCAGGCGCGTCACAGGCAGACGTCGGGGTA<br>CTAGTCATCTCCGCTAGGAAAGGAGAGTATGAAACAGGTTTTGAGAAAGGC<br>GGCCAGACGAGGGAGCATGCAGTTTTGGCCAAGACGCAGGGGATTAACA<br>AACTGATAGTAGTTGTGAATAAGATGGACGACATTACGGTCGAGTGGTCCGA<br>GGAGAGGTATAAAGAATGCCTAGCAAAACTGACTCAGTTCTTAAAAGGATTG<br>GGATATAACCCTAAGACAGATCTAACTTTCATGCCAGTAGCGGCCCAGCAAA<br>CGATGGGGATCAAAGACAGAGTGCCGAAGGACTTATGCCCCTGGTATGATG<br>GACCGAGCCTATTGGAATTCCTTGATTCCATGCAAGCATTGGAGCGTAAGTT<br>ATCCGCCCCATTCATGATGCCCATCAGTGCCAAATATAGAGATATGGGAACA<br>ATGATCGAGGGGAAAATCGAAGCTGGTTTTATAAAGAAAGACCAAAAATATCT<br>AATGATGCCCAATAAAGCGGAAATACAAATATCTGCACTTTATGGTGAGTCTG<br>AAGAAGAGATCGCCCACGCCACCTCAGGGGAACAAATCCGTTTGCGTATTA<br>AGGGGGCCGAGGAAGAGGACATTTACCCGGGGTTCGTCTTATGTTCCCCC<br>AAACGTCCCGTCCATTGCGTGACAACTTTCGAGGCGCAGATCAGACTACTT<br>GAATTGAAATCAATTTTAAGCGCGGGTTTCAACTGCGTTATACACGTGCATTC<br>CGCTACTGAGGAGGTCACGTTTACAGAGCTGTTGCACAAACTGGAACCGAA |

| | |
|---|---|
| | GACAAATAGGAAATCCAAGAAGCCACCGGGTTTCGCGAAACAGGGTATGA<br>ATATCATTGCGAGGCTGGAAGTTACGGGACAAGCGGGTAGCCTGTGTGTCG<br>AGAAGTTCGAGGATTACCCTCAACTTGGAAGGTTCACACTAAGAGATCAAGG<br>GCAGACTATCGCAATAGGTAAAATAACTAAGCTTATCACGGATGCTACTACTC<br>GAGGTGGTTCAGGTGGTGGCGGGAGCGGTGGAGGGAGCAGCGGTGGAG<br>TCTTCACACTCGAAGATTTCGTTGGGGACTGGCGACAGACAGCCGGCTACA<br>ACCTGGACCAAGTCCTTGAACAGGGAGGTGTGTCCAGTTTGTTTCAGAATCT<br>CGGGGTGTCCGTAACTCCGATCCAAAGGATTGTCCTGAGCGGTGAAAATGG<br>GCTGAAGATCGACATCCATGTCATCATCCCGTATGAAGGTCTGAGCGGCGA<br>CCAAATGGGCCAGATCGAAAAAATTTTTAAGGTGGTGTACCCTGTGGATGAT<br>CATCACTTTAAGGTGATCCTGCACTATGGCACACTGGTAATCGACGGGGTTA<br>CGCCGAACATGATCGACTATTTCGGACGGCCGTATGAAGGCATCGCCGTGT<br>TCGACGGCAAAAAGATCACTGTAACAGGGACCCTGTGGAACGGCAACAAA<br>ATTATCGACGAGCGCCTGATCAACCCCGACGGCTCCCTGCTGTTCCGAGTA<br>ACCATCAACGGAGTGACCGGCTGGCGGCTGTGCGAACGCATTCTGGCGTA<br>A |
| **C441A-*Zt*GSPT1-NLuc** | ATGGCCAACGGACAGCAACCTGAGAGCTGGGAAGACGAAATGAGGGAT<br>GAGGACTTGGCTAATCAAACCCAACAGCAGATGAATATGCAGGGCGGGG<br>CGCCTAGGGGAGGGACATTCGTTCCTGGAGCCCAGAGCTTCACTCCAG<br>GAGCGCAATCCTTCAATCCTGGTCAGCAGTACCAGCAATATGGATACGG<br>GCAGCAGTATGGGGGAGGATATCAGCAGTACGGTCAGCAGCAAGGCTA<br>CAACCAGTACCAGCAATATGGCCAATATCAACAACAGGGGTACGGTCAG<br>CAGCAATATGCACAACAAGGTTACAACCAGTACCAGCAACCCCAACAGC<br>AAGCATATGTACCCCCAGCCGCCCGTCAGGCCCCGATGATAGCCAAGAG<br>GGGAGAGGCCCTTCCTACCGCCGCCGCTGCGAAACCAGCTGCTGCCA<br>ATGGCGCAGCCGCCCCCGTTAAGACACTGTCATTGAGTGCTGCAGATAC<br>TGGGGCAGCCGCGCCTAAGGTGGAGAAAGCGACCGGTGGAGCAAAGG<br>TGTTGTCCCTAGGTGGTGCACCGGCTGCTGCAACGCCGAAGGCTGCAC<br>CCGCCCCTAAAGAGGCAGTGAGCAAGGACGCGCCAGAGGCCGGCACT<br>AAGGTCGCTGCCGCCAAAGCTATCGAGAAAACCGGGGAACCGACGGCT<br>GCAGTATCTGGAGCTAGCTCTCCCACACCCAGTGGCAGATCTTCTCCCA<br>CAAGAGCAGACATAAAAGCGGAGGAGAAACGTAAAGCCGACCAGGTGC<br>TAGCCGACCAAGCCGCAGAAGTAGATGAAGAAGCGCTAGCCGAGATGTA<br>TGGTAAGGAGCACGTCAACGTTATCTTTCTTGGCCATGTGGATGCAGGC<br>AAGTCTACCTTAGGTGGGAGCATTCTTTATGCGACTGGGATGGTTGACG<br>AACGTACCATGGATAAATATAAGCGTGATGCTAAGGATATGGGAAGGGAA<br>TCATGGTACTTGTCATGGGCTCTAGACCAGACTAAGGAGGAGAGAGCCC<br>AGGGTAAGACCGTGGAGGTCGGGCGTGGCTTCTTCGAAACGGAAAAGA<br>GGCGTTACTCCATACTAGACGCCCCTGGGCATAAGACTTTCGTACCTAAC<br>ATGCTATCAGGCGCGTCACAGGCAGACGTCGGGGTACTAGTCATCTCCG<br>CTAGGAAAGGAGAGTATGAAACAGGTTTTGAGAAAGGCGGCCAGACGA<br>GGGAGCATGCAGTTTTGGCCAAGACGCAGGGGATTAACAAACTGATAGT<br>AGTTGTGAATAAGATGGACGACATTACGGTCGAGTGGTCCGAGGAGAG<br>GTATAAAGAAGCCCTAGCAAAACTGACTCAGTTCTTAAAAGGATTGGGAT<br>ATAACCCTAAGACAGATCTAACTTTCATGCCAGTAGCGGCCCAGCAAACG |

| | |
|---|---|
| | ATGGGGATCAAAGACAGAGTGCCGAAGGACTTATGCCCCTGGTATGATG<br>GACCGAGCCTATTGGAATTCCTTGATTCCATGCAAGCATTGGAGCGTAAG<br>TTATCCGCCCCATTCATGATGCCCATCAGTGCCAAATATAGAGATATGGGA<br>ACAATGATCGAGGGGAAAATCGAAGCTGGTTTTATAAAGAAAGACCAAAA<br>ATATCTAATGATGCCCAATAAAGCGGAAATACAAATATCTGCACTTTATGGT<br>GAGTCTGAAGAAGAGATCGCCCACGCCACCTCAGGGGAACAAATCCGT<br>TTGCGTATTAAGGGGGCCGAGGAAGAGGACATTTACCCGGGGTTCGTCT<br>TATGTTCCCCCAAACGTCCCGTCCATTGCGTGACAACTTTCGAGGCGCA<br>GATCAGACTACTTGAATTGAAATCAATTTTAAGCGCGGGTTTCAACTGCG<br>TTATACACGTGCATTCCGCTACTGAGGAGGTCACGTTTACAGAGCTGTTG<br>CACAAACTGGAACCGAAGACAAATAGGAAATCCAAGAAGCCACCGGGTT<br>TCGCGAAACAGGGTATGAATATCATTGCGAGGCTGGAAGTTACGGGACA<br>AGCGGGTAGCCTGTGTGTCGAGAAGTTCGAGGATTACCCTCAACTTGGA<br>AGGTTCACACTAAGAGATCAAGGGCAGACTATCGCAATAGGTAAAATAAC<br>TAAGCTTATCACGGATGCTACTACTCGAGGTGGTTCAGGTGGTGGCGGG<br>AGCGGTGGAGGGAGCAGCGGTGGAGTCTTCACACTCGAAGATTTCGTT<br>GGGGACTGGCGACAGACAGCCGGCTACAACCTGGACCAAGTCCTTGAA<br>CAGGGAGGTGTGTCCAGTTTGTTTCAGAATCTCGGGGTGTCCGTAACTC<br>CGATCCAAAGGATTGTCCTGAGCGGTGAAAATGGGCTGAAGATCGACAT<br>CCATGTCATCATCCCGTATGAAGGTCTGAGCGGCGACCAAATGGGCCAG<br>ATCGAAAAAATTTTTAAGGTGGTGTACCCTGTGGATGATCATCACTTTAAG<br>GTGATCCTGCACTATGGCACACTGGTAATCGACGGGGTTACGCCGAACA<br>TGATCGACTATTTCGGACGGCCGTATGAAGGCATCGCCGTGTTCGACGG<br>CAAAAAGATCACTGTAACAGGGACCCTGTGGAACGGCAACAAAATTATC<br>GACGAGCGCCTGATCAACCCCGACGGCTCCCTGCTGTTCCGAGTAACC<br>ATCAACGGAGTGACCGGCTGGCGGCTGTGCGAACGCATTCTGGCGTAA |
| **WT-*Zt*SAR1-FLAG** | ATGTGGATCGTCGATTGGTTCTGGGACCTACTGGCCAACCTTGGACTTG<br>CAAACAAACACGCTAAGTTGCTATTCCTAGGTTTAGATAATGCGGGGAA<br>AACGACTCTTCTGCACATGCTGAAAAATGACAGGGTAGCAATCCTTCAG<br>CCAACACTACATCCAACTAGCGAAGAACTATCCATAGGAAGCTGTAGGT<br>TCACTACCTTCGACCTTGGAGGCCACCAACAAGCGAGGCGTTTATGGA<br>AAGATTATTTTCCTGAAGTCAGCGGAATTGTCTTTCTAGTTGACGCAAAA<br>GACCCCGAAAGATTTTCTGAATCTAAGGCGGAGCTGGACGCGCTATTAA<br>GTATGGAAGATTTAGCGAAGACCCCCTTTCTTATACTGGGTAATAAGAT<br>CGACCACCCAAACGCCGTTTCAGAAGATCAGCTACGTCAGCATCTAGG<br>GTTGTACCAAACCACGGGGAAGGGTAAGGTCCCACTAGACGGTATAAG<br>GCCGATTGAAATTTTTATGTGTAGCGTGGTCATGAGACAGGGGTACGG<br>GGATGGAATCAGGTGGTTGAGCCAGTACGTTGGCGGCTCTTCTGACTA<br>CAAGGATGACGATGACAAGTGA |
| **C64A-*Zt*SAR1-FLAG** | ATGTGGATCGTCGATTGGTTCTGGGACCTACTGGCCAACCTTGGACTTG<br>CAAACAAACACGCTAAGTTGCTATTCCTAGGTTTAGATAATGCGGGGAA<br>AACGACTCTTCTGCACATGCTGAAAAATGACAGGGTAGCAATCCTTCAG<br>CCAACACTACATCCAACTAGCGAAGAACTATCCATAGGAAGCGCTAGGT<br>TCACTACCTTCGACCTTGGAGGCCACCAACAAGCGAGGCGTTTATGGA<br>AAGATTATTTTCCTGAAGTCAGCGGAATTGTCTTTCTAGTTGACGCAAAA<br>GACCCCGAAAGATTTTCTGAATCTAAGGCGGAGCTGGACGCGCTATTAA<br>GTATGGAAGATTTAGCGAAGACCCCCTTTCTTATACTGGGTAATAAGAT<br>CGACCACCCAAACGCCGTTTCAGAAGATCAGCTACGTCAGCATCTAGG<br>GTTGTACCAAACCACGGGGAAGGGTAAGGTCCCACTAGACGGTATAAG<br>GCCGATTGAAATTTTTATGTGTAGCGTGGTCATGAGACAGGGGTACGG |

| | |
|---|---|
| | GGATGGAATCAGGTGGTTGAGCCAGTACGTTGGCGGCTCTTCTGACTA<br>CAAGGATGACGATGACAAGTGA |
| **WT-*Bc*SAR1-FLAG** | ATGTGGCTGCTGGACTGGTTCTGGGACACCCTGGCCAGCCTGGGCCTG<br>CTGAACAAGCACGCCAAGCTGCTGTTCCTGGGCCTGGACAACGCCGGC<br>AAGACCACCCTGCTGCACATGCTGAAGAACGACAGGGTGGCCATCCTG<br>CAGCCCACCCTGCACCCCACCAGCGAGGAGCTGGCCATAGGCAACGT<br>GAAGTTCACCACCTTCGACCTGGGCGGCCACCAGCAGGCCAGGAGGC<br>TGTGGAAGGACTACTTCCCCGAGGTGAGCGGCATCGTGTTCCTGGTGG<br>ACAGCAAGGACCACGAGAGGTTCATCGAGAGCAAGGCCGAGCTGGAC<br>GCCCTGCTGAGCATGGAGGACCTGAGCAAGGTGCCCTTCCTGATCCTG<br>GGCAACAAGATCGACCACCCCGACGCCATCAGCGAGGACCAGCTGAG<br>GCACGAGCTGGGCCTGTACCAGACCACCGGCAAGGGCAAGGTGCCCC<br>TGGAGGGCATCAGGCCCATCGAGGTGTTCATGTGCAGCGTGGTGATGA<br>GGCAGGGCTACGGCGAGGGCATCAGGTGGCTGAGCCAGTACGTGGGC<br>GGCTCTTCTGACTACAAGGATGACGATGACAAGTGA |
| **V64C-*Bc*SAR1-FLAG** | ATGTGGCTGCTGGACTGGTTCTGGGACACCCTGGCCAGCCTGGGCCTG<br>CTGAACAAGCACGCCAAGCTGCTGTTCCTGGGCCTGGACAACGCCGGC<br>AAGACCACCCTGCTGCACATGCTGAAGAACGACAGGGTGGCCATCCTG<br>CAGCCCACCCTGCACCCCACCAGCGAGGAGCTGGCCATCGGCAACTG<br>CAAGTTCACCACCTTCGACCTGGGCGGCCACCAGCAGGCCAGGAGGC<br>TGTGGAAGGACTACTTCCCCGAGGTGAGCGGCATCGTGTTCCTGGTGG<br>ACAGCAAGGACCACGAGAGGTTCATCGAGAGCAAGGCCGAGCTGGAC<br>GCCCTGCTGAGCATGGAGGACCTGAGCAAGGTGCCCTTCCTGATCCTG<br>GGCAACAAGATCGACCACCCCGACGCCATCAGCGAGGACCAGCTGAG<br>GCACGAGCTGGGCCTGTACCAGACCACCGGCAAGGGCAAGGTGCCCC<br>TGGAGGGCATCAGGCCCATCGAGGTGTTCATGTGCAGCGTGGTGATGA<br>GGCAGGGCTACGGCGAGGGCATCAGGTGGCTGAGCCAGTACGTGGGC<br>GGCTCTTCTGACTACAAGGATGACGATGACAAGTGA |
| **WT-*Pg*SAR1-FLAG** | ATGTTCATCCTGAACTGGTTCTGGGACGTGCTGGCCAACCTGGGCCTG<br>GTGAACAAGAACGCCAAGATCCTGTTCCTGGGCCTGGACAACGCCGGC<br>AAGACCACCCTGCTGCACATGCTGAAGAACGACAGGCTGGCCACCCTG<br>CAGCCCACCCTGCACCCCACCAGCGAGGAGCTGGCCATCGGCAACGT<br>GAAGTTCACCACCTACGACCTGGGCGGCCACCAGCAGGCCAGGAGGC<br>TGTGGAAGGAGTACTTCCCCGAGGTGAACGGCATCGTGTTCCTGGTGG<br>ACGCCCAGGACCCCGAGAGGTTCAGCGAGAGCAAGATCGAGCTGGAC<br>GCCCTGCTGAGCATCGAGGAGCTGAGCAAGGTGCCCTTCCTGATCCTG<br>GGCAACAAGATCGACGCCCCCGGCGCCGTGAGCGAGGAGGACCTGAG<br>GCACTGCCTGGGCCTGTACCAGACCACCGGCAAGGGCAAGGTGCCCC<br>TGATCGACATCAGGCCCATCGAGGTGTTCATGTGCAGCATCGTGATGAG<br>GCAGGGCTACGGCGACGGCTTCAGGTGGCTGGCCCAGTACATCGGCG<br>GCTCTTCTGACTACAAGGATGACGATGACAAGTGA |
| **V64C-*Pg*SAR1-FLAG** | ATGTTCATCCTGAACTGGTTCTGGGACGTGCTGGCCAACCTGGGCCTG<br>GTGAACAAGAACGCCAAGATCCTGTTCCTGGGCCTGGACAACGCCGGC<br>AAGACCACCCTGCTGCACATGCTGAAGAACGACAGGCTGGCCACCCTG<br>CAGCCCACCCTGCACCCCACCAGCGAGGAGCTGGCCATCGGCAACTG<br>CAAGTTCACCACCTACGACCTGGGCGGCCACCAGCAGGCCAGGAGGC<br>TGTGGAAGGAGTACTTCCCCGAGGTGAACGGCATCGTGTTCCTGGTGG<br>ACGCCCAGGACCCCGAGAGGTTCAGCGAGAGCAAGATCGAGCTGGAC<br>GCCCTGCTGAGCATCGAGGAGCTGAGCAAGGTGCCCTTCCTGATCCTG<br>GGCAACAAGATCGACGCCCCCGGCGCCGTGAGCGAGGAGGACCTGAG |

| | |
|---|---|
| | GCACTGCCTGGGCCTGTACCAGACCACCGGCAAGGGCAAGGTGCCCC<br>TGATCGACATCAGGCCCATCGAGGTGTTCATGTGCAGCATCGTGATGAG<br>GCAGGGCTACGGCGACGGCTTCAGGTGGCTGGCCCAGTACATCGGCG<br>GCTCTTCTGACTACAAGGATGACGATGACAAGTGA |
| **WT-*Hs*SAR1A-V5** | ATGTCTTTCATCTTTGAGTGGATCTACAATGGCTTCAGCAGTGTGCTCCA<br>GTTCCTAGGACTGTACAAGAAATCTGGAAAACTTGTATTCTTAGGTTTGG<br>ATAATGCAGGCAAAACCACTCTTCTTCACATGCTCAAAGATGACAGATTG<br>GGCCAACATGTTCCAACACTACATCCGACATCAGAAGAGCTAACAATTGC<br>TGGAATGACCTTTACAACTTTTGATCTTGGTGGGCACGAGCAAGCACGT<br>CGCGTTTGGAAAAATTATCTCCCAGCAATTAATGGGATTGTCTTTCTGGT<br>GGACTGTGCAGATCATTCTCGCCTCGTGGAATCCAAAGTTGAGCTTAAT<br>GCTTTAATGACTGATGAAACAATATCCAATGTGCCAATCCTTATCTTGGGT<br>AACAAAATTGACAGAACAGATGCAATCAGTGAAGAAAAACTCCGTGAGAT<br>ATTTGGGCTTTATGGACAGACCACAGGAAAGGGGAATGTGACCCTGAAG<br>GAGCTGAATGCTCGCCCCATGGAAGTGTTCATGTGCAGTGTGCTCAAGA<br>GGCAAGGTTACGGCGAGGGTTTCCGCTGGCTCTCCCAGTATATTGACG<br>GTAAGCCTATCCCTAACCCTCTCCTCGGTCTCGATTCTACGTAG |
| **M69C-*Hs*SAR1A-V5** | ATGTCTTTCATCTTTGAGTGGATCTACAATGGCTTCAGCAGTGTGCTCCA<br>GTTCCTAGGACTGTACAAGAAATCTGGAAAACTTGTATTCTTAGGTTTGG<br>ATAATGCAGGCAAAACCACTCTTCTTCACATGCTCAAAGATGACAGATTG<br>GGCCAACATGTTCCAACACTACATCCGACATCAGAAGAGCTAACAATTGC<br>TGGATGCACCTTTACAACTTTTGATCTTGGTGGGCACGAGCAAGCACGT<br>CGCGTTTGGAAAAATTATCTCCCAGCAATTAATGGGATTGTCTTTCTGGT<br>GGACTGTGCAGATCATTCTCGCCTCGTGGAATCCAAAGTTGAGCTTAAT<br>GCTTTAATGACTGATGAAACAATATCCAATGTGCCAATCCTTATCTTGGGT<br>AACAAAATTGACAGAACAGATGCAATCAGTGAAGAAAAACTCCGTGAGAT<br>ATTTGGGCTTTATGGACAGACCACAGGAAAGGGGAATGTGACCCTGAAG<br>GAGCTGAATGCTCGCCCCATGGAAGTGTTCATGTGCAGTGTGCTCAAGA<br>GGCAAGGTTACGGCGAGGGTTTCCGCTGGCTCTCCCAGTATATTGACG<br>GTAAGCCTATCCCTAACCCTCTCCTCGGTCTCGATTCTACGTAG |
| **WT-*Zt*SAR1-eGFP** | ATGTGGATCGTCGATTGGTTCTGGGACCTACTGGCCAACCTTGGACTTG<br>CAAACAAACACGCTAAGTTGCTATTCCTAGGTTTAGATAATGCGGGGAAA<br>ACGACTCTTCTGCACATGCTGAAAAATGACAGGGTAGCAATCCTTCAGC<br>CAACACTACATCCAACTAGCGAAGAACTATCCATAGGAAGCTGTAGGTTC<br>ACTACCTTCGACCTTGGAGGCCACCAACAAGCGAGGCGTTTATGGAAAG<br>ATTATTTTCCTGAAGTCAGCGGAATTGTCTTTCTAGTTGACGCAAAAGAC<br>CCCGAAAGATTTTCTGAATCTAAGGCGGAGCTGGACGCGCTATTAAGTAT<br>GGAAGATTTAGCGAAGACCCCCTTTCTTATACTGGGTAATAAGATCGACC<br>ACCCAAACGCCGTTTCAGAAGATCAGCTACGTCAGCATCTAGGGTTGTA<br>CCAAACCACGGGGAAGGGTAAGGTCCCACTAGACGGTATAAGGCCGATT<br>GAAATTTTTATGTGTAGCGTGGTCATGAGACAGGGGTACGGGGATGGAA<br>TCAGGTGGTTGAGCCAGTACGTGGGTTCAGGAATGGTGAGCAAGGGCG<br>AGGAGCTGTTCACCGGGGTGGTGCCCATCCTGGTCGAGCTGGACGGC<br>GACGTAAACGGCCACAAGTTCAGCGTGTCCGGCGAGGGCGAGGGCGA<br>TGCCACCTACGGCAAGCTGACCCTGAAGTTCATCTGCACCACCGGCAA<br>GCTGCCCGTGCCCTGGCCCACCCTCGTGACCACCCTGACCTACGGCGT<br>GCAGTGCTTCAGCCGCTACCCCGACCACATGAAGCAGCACGACTTCTT<br>CAAGTCCGCCATGCCCGAAGGCTACGTCCAGGAGCGCACCATCTTCTT<br>CAAGGACGACGGCAACTACAAGACCCGCGCCGAGGTGAAGTTCGAGG<br>GCGACACCCTGGTGAACCGCATCGAGCTGAAGGGCATCGACTTCAAGG<br>AGGACGGCAACATCCTGGGGCACAAGCTGGAGTACAACTACAACAGCC |

| | |
|---|---|
| | ACAACGTCTATATCATGGCCGACAAGCAGAAGAACGGCATCAAGGTGAA CTTCAAGATCCGCCACAACATCGAGGACGGCAGCGTGCAGCTCGCCGA CCACTACCAGCAGAACACCCCCATCGGCGACGGCCCCGTGCTGCTGCC CGACAACCACTACCTGAGCACCCAGTCCGCCCTGAGCAAAGACCCCAA CGAGAAGCGCGATCACATGGTCCTGCTGGAGTTCGTGACCGCCGCCGG GATCACTCTCGGCATGGACGAGCTGTACAAGTACGTTGGCGGCTCTTCT GACTACAAGGATGACGATGACAAGTGA |
| **C64A-*Zt*SAR1-eGFP** | ATGTGGATCGTCGATTGGTTCTGGGACCTACTGGCCAACCTTGGACTTG CAAACAAACACGCTAAGTTGCTATTCCTAGGTTTAGATAATGCGGGGAAA ACGACTCTTCTGCACATGCTGAAAAATGACAGGGTAGCAATCCTTCAGC CAACACTACATCCAACTAGCGAAGAACTATCCATAGGAAGCGCTAGGTTC ACTACCTTCGACCTTGGAGGCCACCAACAAGCGAGGCGTTTATGGAAAG ATTATTTTCCTGAAGTCAGCGGAATTGTCTTTCTAGTTGACGCAAAAGAC CCCGAAAGATTTTCTGAATCTAAGGCGGAGCTGGACGCGCTATTAAGTAT GGAAGATTTAGCGAAGACCCCCTTTCTTATACTGGGTAATAAGATCGACC ACCCAAACGCCGTTTCAGAAGATCAGCTACGTCAGCATCTAGGGTTGTA CCAAACCACGGGGAAGGGTAAGGTCCCACTAGACGGTATAAGGCCGATT GAAATTTTTATGTGTAGCGTGGTCATGAGACAGGGGTACGGGGATGGAA TCAGGTGGTTGAGCCAGTACGTGGGTTCAGGAATGGTGAGCAAGGGCG AGGAGCTGTTCACCGGGGTGGTGCCCATCCTGGTCGAGCTGGACGGC GACGTAAACGGCCACAAGTTCAGCGTGTCCGGCGAGGGCGAGGGCGA TGCCACCTACGGCAAGCTGACCCTGAAGTTCATCTGCACCACCGGCAA GCTGCCCGTGCCCTGGCCCACCCTCGTGACCACCCTGACCTACGGCGT GCAGTGCTTCAGCCGCTACCCCGACCACATGAAGCAGCACGACTTCTT CAAGTCCGCCATGCCCGAAGGCTACGTCCAGGAGCGCACCATCTTCTT CAAGGACGACGGCAACTACAAGACCCGCGCCGAGGTGAAGTTCGAGG GCGACACCCTGGTGAACCGCATCGAGCTGAAGGGCATCGACTTCAAGG AGGACGGCAACATCCTGGGGCACAAGCTGGAGTACAACTACAACAGCC ACAACGTCTATATCATGGCCGACAAGCAGAAGAACGGCATCAAGGTGAA CTTCAAGATCCGCCACAACATCGAGGACGGCAGCGTGCAGCTCGCCGA CCACTACCAGCAGAACACCCCCATCGGCGACGGCCCCGTGCTGCTGCC CGACAACCACTACCTGAGCACCCAGTCCGCCCTGAGCAAAGACCCCAA CGAGAAGCGCGATCACATGGTCCTGCTGGAGTTCGTGACCGCCGCCGG GATCACTCTCGGCATGGACGAGCTGTACAAGTACGTTGGCGGCTCTTCT GACTACAAGGATGACGATGACAAGTGA |
| **Halo-*Hs*SAR1A** | ATGGCAGAAATCGGTACTGGCTTTCCATTCGACCCCCATTATGTGGAAGT CCTGGGCGAGCGCATGCACTACGTCGATGTTGGTCCGCGCGATGGCAC CCCTGTGCTGTTCCTGCACGGTAACCCGACCTCCTCCTACGTGTGGCG CAACATCATCCCGCATGTTGCACCGACCCATCGCTGCATTGCTCCAGAC CTGATCGGTATGGGCAAATCCGACAAACCAGACCTGGGTTATTTCTTCGA CGACCACGTCCGCTTCATGGATGCCTTCATCGAAGCCCTGGGTCTGGAA GAGGTCGTCCTGGTCATTCACGACTGGGGCTCCGCTCTGGGTTTCCAC TGGGCCAAGCGCAATCCAGAGCGCGTCAAAGGTATTGCATTTATGGAGT TCATCCGCCCTATCCCGACCTGGGACGAATGGCCAGAATTTGCCCGCGA GACCTTCCAGGCCTTCCGCACCACCGACGTCGGCCGCAAGCTGATCAT CGATCAGAACGTTTTTATCGAGGGTACGCTGCCGATGGGTGTCGTCCGC CCGCTGACTGAAGTCGAGATGGACCATTACCGCGAGCCGTTCCTGAATC CTGTTGACCGCGAGCCACTGTGGCGCTTCCCAAACGAGCTGCCAATCG CCGGTGAGCCAGCGAACATCGTCGCGCTGGTCGAAGAATACATGGACT GGCTGCACCAGTCCCCTGTCCCGAAGCTGCTGTTCTGGGGCACCCCAG GCGTTCTGATCCCACCGGCCGAAGCCGCTCGCCTGGCCAAAAGCCTGC |

| | |
|---|---|
| | CTAACTGCAAGGCTGTGGACATCGGCCCGGGTCTGAATCTGCTGCAAG<br>AAGACAACCCGGACCTGATCGGCAGCGAGATCGCGCGCTGGCTGTCGA<br>CGCTCGAGATTTCCGGCGAGCCAACCACTGAGGATCTGTACTTTCAGAG<br>CGATAACGCGATCGCTTCCGAATTCAGAGCTCAACCGCGGATATCTAGAT<br>TTGGGCCCAATTCCTGCAGGATTTTGCGGCCGCGGATCTCCACAAGTTT<br>GTACAAAAAAGTTGGCATGTCTTTCATCTTTGAGTGGATCTACAATGGCT<br>TCAGCAGTGTGCTCCAGTTCCTAGGACTGTACAAGAAATCTGGAAAACT<br>TGTATTCTTAGGTTTGGATAATGCAGGCAAAACCACTCTTCTTCACATGCT<br>CAAAGATGACAGATTGGGCCAACATGTTCCAACACTACATCCGACATCAG<br>AAGAGCTAACAATTGCTGGAATGACCTTTACAACTTTTGATCTTGGTGGG<br>CACGAGCAAGCACGTCGCGTTTGGAAAAATTATCTCCCAGCAATTAATGG<br>GATTGTCTTTCTGGTGGACTGTGCAGATCATTCTCGCCTCGTGGAATCC<br>AAAGTTGAGCTTAATGCTTTAATGACTGATGAAACAATATCCAATGTGCCA<br>ATCCTTATCTTGGGTAACAAAATTGACAGAACAGATGCAATCAGTGAAGA<br>AAAACTCCGTGAGATATTTGGGCTTTATGGACAGACCACAGGAAAGGGG<br>AATGTGACCCTGAAGGAGCTGAATGCTCGCCCCATGGAAGTGTTCATGT<br>GCAGTGTGCTCAAGAGGCAAGGTTACGGCGAGGGTTTCCGCTGGCTCT<br>CCCAGTATATTGACTGCCCAACTTTCTTGTACAAAGTGGTAAGTAACTAA |
| **Halo-*Hs*SEC23B** | ATGGCAGAAATCGGTACTGGCTTTCCATTCGACCCCCATTATGTGGAAGT<br>CCTGGGCGAGCGCATGCACTACGTCGATGTTGGTCCGCGCGATGGCAC<br>CCCTGTGCTGTTCCTGCACGGTAACCCGACCTCCTCCTACGTGTGGCG<br>CAACATCATCCCGCATGTTGCACCGACCCATCGCTGCATTGCTCCAGAC<br>CTGATCGGTATGGGCAAATCCGACAAACCAGACCTGGGTTATTTCTTCGA<br>CGACCACGTCCGCTTCATGGATGCCTTCATCGAAGCCCTGGGTCTGGAA<br>GAGGTCGTCCTGGTCATTCACGACTGGGGCTCCGCTCTGGGTTTCCAC<br>TGGGCCAAGCGCAATCCAGAGCGCGTCAAAGGTATTGCATTTATGGAGT<br>TCATCCGCCCTATCCCGACCTGGGACGAATGGCCAGAATTTGCCCGCGA<br>GACCTTCCAGGCCTTCCGCACCACCGACGTCGGCCGCAAGCTGATCAT<br>CGATCAGAACGTTTTTATCGAGGGTACGCTGCCGATGGGTGTCGTCCGC<br>CCGCTGACTGAAGTCGAGATGGACCATTACCGCGAGCCGTTCCTGAATC<br>CTGTTGACCGCGAGCCACTGTGGCGCTTCCCAAACGAGCTGCCAATCG<br>CCGGTGAGCCAGCGAACATCGTCGCGCTGGTCGAAGAATACATGGACT<br>GGCTGCACCAGTCCCCTGTCCCGAAGCTGCTGTTCTGGGGCACCCCAG<br>GCGTTCTGATCCCACCGGCCGAAGCCGCTCGCCTGGCCAAAAGCCTGC<br>CTAACTGCAAGGCTGTGGACATCGGCCCGGGTCTGAATCTGCTGCAAG<br>AAGACAACCCGGACCTGATCGGCAGCGAGATCGCGCGCTGGCTGTCGA<br>CGCTCGAGATTTCCGGCGAGCCAACCACTGAGGATCTGTACTTTCAGAG<br>CGATAACGCGATCGCTTCCGAATTCAGAGCTCAACCGCGGATATCTAGAT<br>TTGGGCCCAATTCCTGCAGGATTTTGCGGCCGCGGATCTCCACAAGTTT<br>GTACAAAAAAGTTGGCATGGCGACATACCTGGAGTTCATCCAGCAGAAT<br>GAAGAACGGGATGGTGTGCGTTTTAGTTGGAACGTGTGGCCTTCCAGC<br>CGGCTGGAGGCTACAAGAATGGTTGTACCCCTGGCTTGTCTCCTTACTC<br>CTTTGAAAGAACGTCCAGACCTACCTCCTGTACAATATGAACCTGTGCTT<br>TGCAGCAGGCCAACTTGTAAAGCTGTTCTCAACCCACTTTGTCAGGTTG<br>ATTATCGAGCAAAACTTTGGGCCTGTAATTTCTGTTTTCAAAGAAATCAGT<br>TTCCTCCAGCTTATGGAGGCATATCTGAGGTGAATCAACCTGCCGAATTG<br>ATGCCCCAGTTTTCTACAATTGAGTACGTGATACAGCGAGGTGCTCAGTC<br>CCCTCTGATCTTTCTCTATGTGGTTGACACATGCCTGGAGGAAGATGAC<br>CTTCAAGCACTCAAAGAGTCCCTGCAGATGTCCCTGAGTCTTCTTCCTC<br>CAGATGCTCTGGTGGGTCTGATCACATTTGGAAGGATGGTGCAGGTTCA<br>TGAGCTAAGCTGTGAAGGAATCTCCAAAAGTTATGTCTTCCGAGGGACC |

| | |
|---|---|
| | AAGGATTTAACTGCAAAGCAAATACAGGATATGTTGGGCCTGACCAAGCC<br>AGCCATGCCCATGCAGCAAGCACGACCTGCACAACCACAGGAGCACCC<br>TTTTGCTTCAAGCAGATTTCTGCAGCCTGTTCACAAGATTGATATGAACC<br>TCACTGATCTTCTTGGGGAGCTACAGAGGGACCCATGGCCAGTAACTCA<br>GGGGAAGAGACCTTTGCGATCCACTGGTGTGGCTTTGTCCATTGCTGTT<br>GGCTTGCTGGAGGGCACTTTTCCAAACACAGGAGCCAGGATCATGCTG<br>TTTACTGGAGGTCCCCCTACCCAAGGGCCTGGCATGGTGGTTGGAGAT<br>GAATTAAAGATTCCTATTCGTTCTTGGCATGATATTGAGAAAGATAATGCA<br>CGATTCATGAAAAAGGCAACCAAGCACTATGAGATGCTTGCTAATCGAAC<br>AGCTGCAAATGGTCACTGCATTGATATTTATGCTTGTGCCCTTGATCAAA<br>CTGGACTTTTGGAGATGAAGTGTTGTGCAAATCTTACTGGAGGCTACGT<br>GGTAATGGGAGATTCTTTCAACACTTCTCTCTTCAAGCAGACATTCCAAA<br>GAATCTTTACTAAAGATTTTAATGGAGATTTCCGAATGGCATTTGGTGCTA<br>CTTTGGACGTAAAGACCTCTCGGGAACTGAAGATTGCAGGAGCCATTGG<br>TCCATGCGTATCTCTGAATGTGAAAGGACTGTGTGTGTCAGAAAATGAG<br>CTTGGTGTTGGTGGCACGAGTCAGTGGAAAATCTGTGGCCTAGATCCTA<br>CATCTACACTTGGCATCTATTTTGAAGTTGTCAATCAGCACAACACCCCG<br>ATCCCCCAAGGAGGCAGAGGAGCCATCCAGTTTGTCACGCATTATCAGC<br>ACTCCAGCACCCAGAGACGCATCCGCGTGACCACCATCGCCCGAAATT<br>GGGCAGATGTACAGAGTCAGCTCAGGCACATAGAAGCAGCATTTGACCA<br>GGAGGCTGCGGCAGTGTTGATGGCACGGCTTGGGGTGTTCCGAGCGG<br>AGTCAGAGGAGGGGCCCGATGTGCTCCGGTGGCTGGACCGACAACTC<br>ATCCGACTGTGTCAAAAGTTTGGACAGTATAACAAAGAAGACCCCACTTC<br>TTTTAGGTTATCAGATTCCTTTTCTCTATATCCTCAGTTTATGTTCCATCTG<br>AGAAGATCTCCATTTCTTCAAGTGTTTAACAACAGTCCTGATGAGTCGTC<br>ATATTACAGACATCATTTTGCCCGGCAGGACCTGACCCAGTCCCTCATCA<br>TGATCCAGCCCATTCTCTACTCTTACTCCTTTCATGGGCCACCAGAGCCA<br>GTACTCTTGGATAGCAGCAGCATTCTAGCTGACAGAATTTTGCTGATGGA<br>TACTTTCTTTCAAATTGTCATTTATCTTGGTGAGACCATAGCCCAGTGGCG<br>TAAAGCTGGCTACCAGGACATGCCCGAGTATGAAAACTTCAAGCACCTT<br>CTGCAGGCACCACTGGATGATGCTCAAGAAATTCTGCAAGCACGCTTCC<br>CGATGCCACGTTACATCAACACGGAGCATGGAGGCAGTCAGGCTCGATT<br>CCTTTTGTCCAAAGTGAACCCATCTCAGACACACAATAACCTGTATGCTT<br>GGGGACAGGAAACTGGAGCACCCATCCTAACTGATGATGTTAGCCTGCA<br>GGTGTTCATGGACCATTTGAAGAAGCTGGCTGTCTCCAGTGCCTGTTAC<br>CCAACTTTCTTGTACAAAGTGGTAAGTAACTAA |
| **Halo-*Hs*SEC31A** | ATGGCAGAAATCGGTACTGGCTTTCCATTCGACCCCCATTATGTGGAAGT<br>CCTGGGCGAGCGCATGCACTACGTCGATGTTGGTCCGCGCGATGGCAC<br>CCCTGTGCTGTTCCTGCACGGTAACCCGACCTCCTCCTACGTGTGGCG<br>CAACATCATCCCGCATGTTGCACCGACCCATCGCTGCATTGCTCCAGAC<br>CTGATCGGTATGGGCAAATCCGACAAACCAGACCTGGGTTATTTCTTCGA<br>CGACCACGTCCGCTTCATGGATGCCTTCATCGAAGCCCTGGGTCTGGAA<br>GAGGTCGTCCTGGTCATTCACGACTGGGGCTCCGCTCTGGGTTTCCAC<br>TGGGCCAAGCGCAATCCAGAGCGCGTCAAAGGTATTGCATTTATGGAGT<br>TCATCCGCCCTATCCCGACCTGGGACGAATGGCCAGAATTTGCCCGCGA<br>GACCTTCCAGGCCTTCCGCACCACCGACGTCGGCCGCAAGCTGATCAT<br>CGATCAGAACGTTTTTATCGAGGGTACGCTGCCGATGGGTGTCGTCCGC<br>CCGCTGACTGAAGTCGAGATGGACCATTACCGCGAGCCGTTCCTGAATC<br>CTGTTGACCGCGAGCCACTGTGGCGCTTCCCAAACGAGCTGCCAATCG<br>CCGGTGAGCCAGCGAACATCGTCGCGCTGGTCGAAGAATACATGGACT<br>GGCTGCACCAGTCCCCTGTCCCGAAGCTGCTGTTCTGGGGCACCCCAG |

| | |
|---|---|
| | GCGTTCTGATCCCACCGGCCGAAGCCGCTCGCCTGGCCAAAAGCCTGC<br>CTAACTGCAAGGCTGTGGACATCGGCCCGGGTCTGAATCTGCTGCAAG<br>AAGACAACCCGGACCTGATCGGCAGCGAGATCGCGCGCTGGCTGTCGA<br>CGCTCGAGATTTCCGGCGAGCCAACCACTGAGGATCTGTACTTTCAGAG<br>CGATAACGCGATCGCTTCCGAATTCAGAGCTCAACCGCGGATATCTAGAT<br>TTGGGCCCAATTCCTGCAGGATTTTGCGGCCGCGGATCTCCACAAGTTT<br>GTACAAAAAAGTTGGCATGAAGTTAAAGGAAGTAGATCGTACAGCCATGC<br>AGGCATGGAGCCCTGCCCAGAATCACCCCATTTACCTAGCAACAGGAAC<br>ATCTGCTCAGCAATTGGATGCAACATTTAGTACGAATGCTTCCCTTGAGA<br>TATTTGAATTAGACCTCTCTGATCCATCCTTGGATATGAAATCTTGTGCCA<br>CATTCTCCTCTTCTCACAGGTACCACAAGTTGATTTGGGGGCCTTATAAA<br>ATGGATTCCAAAGGAGATGTCTCTGGAGTTCTGATTGCAGGTGGTGAAA<br>ATGGAAATATTATTCTCTATGATCCTTCTAAAATTATAGCTGGAGACAAGGA<br>AGTTGTGATTGCCCAGAATGACAAGCATACTGGCCCAGTGAGAGCCTTG<br>GATGTGAACATTTTCCAGACTAATCTGGTAGCTTCTGGTGCTAATGAATCT<br>GAAATCTACATATGGGATCTAAATAATTTTGCAACCCCAATGACACCAGGA<br>GCCAAAACACAGCCGCCAGAAGATATCAGCTGCATTGCATGGAACAGAC<br>AAGTTCAGCATATTTTAGCATCAGCCAGTCCCAGTGGCCGGGCCACTGT<br>ATGGGATCTTAGAAAAAATGAGCCAATCATCAAAGTCAGTGACCATAGTA<br>ACAGAATGCATTGTTCTGGGTTGGCATGGCATCCTGATGTTGCTACTCAG<br>ATGGTCCTTGCCTCCGAGGATGACCGGTTACCAGTGATCCAGATGTGGG<br>ATCTTCGATTTGCTTCCTCTCCACTTCGTGTCCTGGAAAACCATGCCAGG<br>GGGATTTTGGCAATTGCTTGGAGCATGGCAGATCCTGAATTGTTACTGAG<br>CTGTGGAAAAGATGCTAAGATTCTCTGCTCCAATCCAAACACAGGAGAG<br>GTGTTATATGAACTTCCCACCAACACACAGTGGTGCTTCGATATTCAGTG<br>GTGTCCCCGAAATCCTGCTGTCTTATCAGCTGCTTCGTTTGATGGGCGT<br>ATCAGTGTTTATTCTATCATGGGAGGAAGCACAGATGGTTTAAGACAGAA<br>ACAAGTTGACAAGCTTTCATCATCTTTTGGGAATCTTGATCCCTTTGGCA<br>CAGGACAGCCCCTTCCTCCGTTACAAATTCCACAGCAGACTGCTCAGCA<br>TAGTATAGTGCTGCCTCTGAAGAAGCCGCCCAAGTGGATTCGAAGGCCT<br>GTTGGTGCTTCTTTTTCATTTGGAGGCAAACTGGTTACGTTTGAGAATGT<br>CAGAATGCCTTCTCATCAGGGAGCTGAGCAGCAGCAGCAGCAGCACCA<br>TGTGTTCATTAGTCAGGTTGTAACAGAAAAGGAGTTCCTCAGCCGATCA<br>GACCAACTTCAGCAGGCTGTGCAGTCACAAGGATTTATCAATTATTGCCA<br>AAAAAAAATTGATGCTTCTCAGACTGAATTTGAGAAAAATGTGTGGTCCT<br>TTTTGAAGGTAAACTTTGAGGATGATTCTCGTGGAAAATACCTTGAACTT<br>CTAGGATACAGAAAAGAAGATCTAGGAAAGAAGGTAAACTTTTGGGAAAG<br>TTACCCAACTTTCTTGTACAAAGTGGTAAGTAACTAA |
| **Halo-*Hs*TMED2** | TTAGTTACTTACCACTTTGTACAAGAAAGTTGGGTAAACAACTCTCCGGA<br>CTTCAAAAAATCTCTTCAGGTAGTAGATCTGTCCCAATGTCATGGCAACT<br>AGAACAAGAGCTTCAAAGAAGGACCAAAGGACCACTCTGCTGTTTGTGT<br>TGTCGTTGATGGCTCTGTGTATTCTCTCCCGGACTTCCATGTATTCCTGT<br>TCGTGCTTTACAGCTGTCATCGCCACTGCTAGCTCATTGATCATTTCTTCT<br>AGCTTGTTCTGGTGAGCTTCTGTTTCCATATCTTGTCCTTTTGGAGCCTC<br>CCCAATATCAATGGTGAACATCACTATTTTTGGAGTCATGGTGGACATCC<br>GGTTACTAAAACAAAATTTGTATGTTCCATCCATGTGAGCAGCAAATGTGT<br>ATTTCCCACTGGATTCTCTGTCTCCTTTGTAAATTCCTTTGTTATCTGGTC<br>CTGTAATCTCCACGTCGATGTCCAGGAAGCCGCCCTCCGCCACCTCGAA<br>GATGAGGCCCATCTTGGTGCCCGAGGTGACCCGCTCAAAGAAGCACTC<br>TTCAGCATGGGCGTCGATGCTAACGAAATAGCCCGAGACCGTGGCCAG<br>GAGAGCGGCCAGGAGCACCAGCAGTTCAGCAAGCGTCACCATGCCAAT |

| | |
|---|---|
| | TTTTTTGTACAAACTTGTGGAGATCCGCGGCCGCAAAATCCTGCAGGAA<br>TTGGGCCCAAATCTAGATATCCGCGGTTGAGCTCTGAATTCGGAAGCGA<br>TCGCGTTATCGCTCTGAAAGTACAGATCCTCAGTGGTTGGCTCGCCGGA<br>AATCTCGAGCGTCGACAGCCAGCGCGATCTCGCTGCCGATCAGGTC<br>CGGGTTGTCTTCTTGCAGCAGATTCAGACCCGGGCCGATGTCCACAGC<br>CTTGCAGTTAGGCAGGCTTTTGGCCAGGCGAGCGGCTTCGGCCGGTG<br>GGATCAGAACGCCTGGGGTGCCCCAGAACAGCAGCTTCGGGACAGGG<br>GACTGGTGCAGCCAGTCCATGTATTCTTCGACCAGCGCGACGATGTTCG<br>CTGGCTCACCGGCGATTGGCAGCTCGTTTGGGAAGCGCCACAGTGGCT<br>CGCGGTCAACAGGATTCAGGAACGGCTCGCGGTAATGGTCCATCTCGA<br>CTTCAGTCAGCGGGCGGACGACACCCATCGGCAGCGTACCCTCGATAA<br>AAACGTTCTGATCGATGATCAGCTTGCGGCCGACGTCGGTGGTGCGGA<br>AGGCCTGGAAGGTCTCGCGGGCAAATTCTGGCCATTCGTCCCAGGTCG<br>GGATAGGGCGGATGAACTCCATAAATGCAATACCTTTGACGCGCTCTGG<br>ATTGCGCTTGGCCCAGTGGAAACCCAGAGCGGAGCCCCAGTCGTGAAT<br>GACCAGGACGACCTCTTCCAGACCCAGGGCTTCGATGAAGGCATCCAT<br>GAAGCGGACGTGGTCGTCGAAGAAATAACCCAGGTCTGGTTTGTCGGA<br>TTTGCCCATACCGATCAGGTCTGGAGCAATGCAGCGATGGGTCGGTGCA<br>ACATGCGGGATGATGTTGCGCCACACGTAGGAGGAGGTCGGGTTACCG<br>TGCAGGAACAGCACAGGGGTGCCATCGCGCGGACCAACATCGACGTAG<br>TGCATGCGCTCGCCCAGGACTTCCACATAATGGGGGTCGAATGGAAAG<br>CCAGTACCGATTTCTGCCAT |
| **Halo-*Hs*TRAPPC1** | ATGGCAGAAATCGGTACTGGCTTTCCATTCGACCCCCATTATGTGGAAGT<br>CCTGGGCGAGCGCATGCACTACGTCGATGTTGGTCCGCGCGATGGCAC<br>CCCTGTGCTGTTCCTGCACGGTAACCCGACCTCCTCCTACGTGTGGCG<br>CAACATCATCCCGCATGTTGCACCGACCCATCGCTGCATTGCTCCAGAC<br>CTGATCGGTATGGGCAAATCCGACAAACCAGACCTGGGTTATTTCTTCGA<br>CGACCACGTCCGCTTCATGGATGCCTTCATCGAAGCCCTGGGTCTGGAA<br>GAGGTCGTCCTGGTCATTCACGACTGGGGCTCCGCTCTGGGTTTCCAC<br>TGGGCCAAGCGCAATCCAGAGCGCGTCAAAGGTATTGCATTTATGGAGT<br>TCATCCGCCCTATCCCGACCTGGGACGAATGGCCAGAATTTGCCCGCGA<br>GACCTTCCAGGCCTTCCGCACCACCGACGTCGGCCGCAAGCTGATCAT<br>CGATCAGAACGTTTTTATCGAGGGTACGCTGCCGATGGGTGTCGTCCGC<br>CCGCTGACTGAAGTCGAGATGGACCATTACCGCGAGCCGTTCCTGAATC<br>CTGTTGACCGCGAGCCACTGTGGCGCTTCCCAAACGAGCTGCCAATCG<br>CCGGTGAGCCAGCGAACATCGTCGCGCTGGTCGAAGAATACATGGACT<br>GGCTGCACCAGTCCCCTGTCCCGAAGCTGCTGTTCTGGGGCACCCCAG<br>GCGTTCTGATCCCACCGGCCGAAGCCGCTCGCCTGGCCAAAAGCCTGC<br>CTAACTGCAAGGCTGTGGACATCGGCCCGGGTCTGAATCTGCTGCAAG<br>AAGACAACCCGGACCTGATCGGCAGCGAGATCGCGCGCTGGCTGTCGA<br>CGCTCGAGATTTCCGGCGAGCCAACCACTGAGGATCTGTACTTTCAGAG<br>CGATAACGCGATCGCTTCCGAATTCAGAGCTCAACCGCGGATATCTAGAT<br>TTGGGCCCAATTCCTGCAGGATTTTGCGGCCGCGGATCTCCACAAGTTT<br>GTACAAAAAAGTTGGCATGACTGTCCACAACCTGTACCTGTTTGACCGG<br>AATGGAGTGTGTCTGCACTACAGCGAATGGCACCGCAAGAAGCAAGCA<br>GGGATTCCCAAGGAGGAGGAGTATAAGCTGATGTACGGGATGCTCTTCT<br>CTATCCGCTCGTTTGTCAGCAAGATGTCCCCGCTAGACATGAAGGATGG<br>CTTCCTGGCCTTCCAAACTAGCCGTTACAAACTCCATTACTACGAGACGC<br>CCACTGGGATCAAAGTTGTCATGAATACTGACTTGGGCGTGGACCCAT<br>CCGAGATGTGCTGCACCACATCTACAGTGCGCTGTATGTGGAGCTGGTG<br>GTGAAGAATCCCCTGTGCCCGCTGGGCCAAACTGTGCAAAGTGAGCTC |

| | |
|---|---|
| | TTTCGCTCCCGACTGGACTCCTATGTTCGCTCTCTGCCCTTCTTCTCCG<br>CCCGGGCTGGCTGCCCAACTTTCTTGTACAAAGTGGTAAGTAACTAA |
| **Halo-*Hs*STX5** | ATGGCAGAAATCGGTACTGGCTTTCCATTCGACCCCCATTATGTGGAAGT<br>CCTGGGCGAGCGCATGCACTACGTCGATGTTGGTCCGCGCGATGGCAC<br>CCCTGTGCTGTTCCTGCACGGTAACCCGACCTCCTCCTACGTGTGGCG<br>CAACATCATCCCGCATGTTGCACCGACCCATCGCTGCATTGCTCCAGAC<br>CTGATCGGTATGGGCAAATCCGACAAACCAGACCTGGGTTATTTCTTCGA<br>CGACCACGTCCGCTTCATGGATGCCTTCATCGAAGCCCTGGGTCTGGAA<br>GAGGTCGTCCTGGTCATTCACGACTGGGGCTCCGCTCTGGGTTTCCAC<br>TGGGCCAAGCGCAATCCAGAGCGCGTCAAAGGTATTGCATTTATGGAGT<br>TCATCCGCCCTATCCCGACCTGGGACGAATGGCCAGAATTTGCCCGCGA<br>GACCTTCCAGGCCTTCCGCACCACCGACGTCGGCCGCAAGCTGATCAT<br>CGATCAGAACGTTTTTATCGAGGGTACGCTGCCGATGGGTGTCGTCCGC<br>CCGCTGACTGAAGTCGAGATGGACCATTACCGCGAGCCGTTCCTGAATC<br>CTGTTGACCGCGAGCCACTGTGGCGCTTCCCAAACGAGCTGCCAATCG<br>CCGGTGAGCCAGCGAACATCGTCGCGCTGGTCGAAGAATACATGGACT<br>GGCTGCACCAGTCCCCTGTCCCGAAGCTGCTGTTCTGGGGCACCCCAG<br>GCGTTCTGATCCCACCGGCCGAAGCCGCTCGCCTGGCCAAAAGCCTGC<br>CTAACTGCAAGGCTGTGGACATCGGCCCGGGTCTGAATCTGCTGCAAG<br>AAGACAACCCGGACCTGATCGGCAGCGAGATCGCGCGCTGGCTGTCGA<br>CGCTCGAGATTTCCGGCGAGCCAACCACTGAGGATCTGTACTTTCAGAG<br>CGATAACGCGATCGCTTCCGAATTCAGAGCTCAACCGCGGATATCTAGAT<br>TTGGGCCCAATTCCTGCAGGATTTTGCGGCCGCGGATCTCCACAAGTTT<br>GTACAAAAAAGTTGGCATGTCCTGCCGGGATCGGACCCAGGAGTTTCTG<br>TCTGCCTGCAAGTCGCTGCAGACCCGTCAGAATGGAATCCAGACAAATA<br>AGCCAGCTTTGCGTGCTGTCCGACAACGCAGTGAATTCACCCTCATGGC<br>CAAGCGCATTGGGAAAGACCTTAGCAACACATTTGCCAAGCTGGAGAAG<br>CTGACAATCTTGGCAAAGCGCAAGTCCCTCTTTGATGATAAAGCAGTGG<br>AAATTGAAGAGCTAACATATATCATCAAACAGGACATCAATAGCCTCAACA<br>AACAAATTGCTCAGCTCCAGGATTTCGTGAGAGCCAAGGGCAGCCAGA<br>GTGGCCGGCACCTGCAGACCCACTCCAACACCATTGTGGTCTCCTTGC<br>AGTCGAAACTGGCTTCTATGTCCAATGACTTCAAATCGGTTTTAGAAGTG<br>AGGACAGAGAACCTGAAGCAGCAGAGGAGCCGGAGAGAGCAGTTCTC<br>CCGGGCACCTGTGTCAGCCCTGCCCCTTGCCCCTAACCACCTGGGCGG<br>TGGTGCTGTGGTTCTGGGGGCAGAGTCCCATGCCTCCAAGGATGTCGC<br>CATCGACATGATGGACTCTCGGACCAGCCAGCAGCTGCAGCTCATTGAC<br>GAGCAGGATTCCTACATCCAGAGTCGGGCAGACACCATGCAGAACATTG<br>AGTCGACAATTGTTGAGTTGGGCTCCATCTTTCAGCAGTTGGCACACAT<br>GGTTAAGGAACAGGAGGAAACCATTCAGAGTGTCCTTTTGTTTCCTCTT<br>CTCCCGGCCTTGTCCCCAGGATCGACGAGAACGTGCTACCCAACTTTCT<br>TGTACAAAGTGGTAAGTAACTAA |
| **Halo-*Hs*TFG** | ATGGCAGAAATCGGTACTGGCTTTCCATTCGACCCCCATTATGTGGAAGT<br>CCTGGGCGAGCGCATGCACTACGTCGATGTTGGTCCGCGCGATGGCAC<br>CCCTGTGCTGTTCCTGCACGGTAACCCGACCTCCTCCTACGTGTGGCG<br>CAACATCATCCCGCATGTTGCACCGACCCATCGCTGCATTGCTCCAGAC<br>CTGATCGGTATGGGCAAATCCGACAAACCAGACCTGGGTTATTTCTTCGA<br>CGACCACGTCCGCTTCATGGATGCCTTCATCGAAGCCCTGGGTCTGGAA<br>GAGGTCGTCCTGGTCATTCACGACTGGGGCTCCGCTCTGGGTTTCCAC<br>TGGGCCAAGCGCAATCCAGAGCGCGTCAAAGGTATTGCATTTATGGAGT<br>TCATCCGCCCTATCCCGACCTGGGACGAATGGCCAGAATTTGCCCGCGA<br>GACCTTCCAGGCCTTCCGCACCACCGACGTCGGCCGCAAGCTGATCAT |

| | |
|---|---|
| | CGATCAGAACGTTTTTATCGAGGGTACGCTGCCGATGGGTGTCGTCCGC<br>CCGCTGACTGAAGTCGAGATGGACCATTACCGCGAGCCGTTCCTGAATC<br>CTGTTGACCGCGAGCCACTGTGGCGCTTCCCAAACGAGCTGCCAATCG<br>CCGGTGAGCCAGCGAACATCGTCGCGCTGGTCGAAGAATACATGGACT<br>GGCTGCACCAGTCCCCTGTCCCGAAGCTGCTGTTCTGGGGCACCCCAG<br>GCGTTCTGATCCCACCGGCCGAAGCCGCTCGCCTGGCCAAAAGCCTGC<br>CTAACTGCAAGGCTGTGGACATCGGCCCGGGTCTGAATCTGCTGCAAG<br>AAGACAACCCGGACCTGATCGGCAGCGAGATCGCGCGCTGGCTGTCGA<br>CGCTCGAGATTTCCGGCGAGCCAACCACTGAGGATCTGTACTTTCAGAG<br>CGATAACGCGATCGCTTCCGAATTCAGAGCTCAACCGCGGATATCTAGAT<br>TTGGGCCCAATTCCTGCAGGATTTTGCGGCCGCGGATCTCCACAAGTTT<br>GTACAAAAAAGTTGGCATGAACGGACAGTTGGATCTAAGTGGGAAGCTA<br>ATCATCAAAGCTCAACTTGGGGAGGATATTCGGCGAATTCCTATTCATAAT<br>GAAGATATTACTTATGATGAATTAGTGCTAATGATGCAACGAGTTTTCAGA<br>GGAAAACTTCTGAGTAATGATGAAGTAACAATAAAGTATAAAGATGAAGAT<br>GGAGATCTTATAACAATTTTTGATAGTTCTGACCTTTCCTTTGCAATTCAG<br>TGCAGTAGGATACTGAAACTGACATTATTTGTTAATGGCCAGCCAAGACC<br>CCTTGAATCAAGTCAGGTGAAATATCTCCGTCGAGAACTGATAGAACTTC<br>GAAATAAAGTGAATCGTTTATTGGATAGCTTGGAACCACCTGGAGAACCA<br>GGACCTTCCACCAATATTCCTGAAAATGATACTGTGGATGGTAGGGAAGA<br>AAAGTCTGCTTCTGATTCTTCTGGAAAACAGTCTACTCAGGTTATGGCAG<br>CAAGTATGTCTGCTTTTGATCCTTTAAAAAACCAAGATGAAATCAATAAAA<br>ATGTTATGTCAGCGTTTGGCTTAACAGATGATCAGGTTTCAGGGCCACCC<br>AGTGCTCCTGCAGAAGATCGTTCAGGAACACCCGACAGCATTGCTTCCT<br>CCTCCTCAGCAGCTCACCCACCAGGCGTTCAGCCACAGCAGCCACCAT<br>ATACAGGAGCTCAGACTCAAGCAGGTCAGATTGAAGGTCAGATGTACCA<br>ACAGTACCAGCAACAGGCCGGCTATGGTGCACAGCAGCCGCAGGCTCC<br>ACCTCAGCAGCCTCAACAGTATGGTATTCAGTATTCAGCAAGCTATAGTC<br>AGCAGACTGGACCTCAACAACCTCAGCAGTTCCAGGGATATGGCCAGC<br>AACCAACTTCCCAGGCACCAGCTCCTGCCTTTTCTGGTCAGCCTCAACA<br>ACTGCCTGCTCAGCCGCCACAGCAGTACCAGGCGAGCAATTATCCTGCA<br>CAAACTTACACTGCCCAAACTTCTCAGCCTACTAATTATACTGTGGCTCCT<br>GCCTCTCAACCTGGAATGGCTCCAAGCCAACCTGGGGCCTATCAACCAA<br>GACCAGGTTTTACTTCACTTCCTGGAAGTACCATGACCCCTCCTCCAAGT<br>GGGCCTAATCCTTATGCGCGTAACCGTCCTCCCTTTGGTCAGGGCTATA<br>CCCAACCTGGACGTGGTTATCGATACCCAACTTTCTTGTACAAAGTGGTA<br>AGTAACTAA |
| **WT-*Bc*SAR1-eGFP** | ATGTGGCTGCTGGACTGGTTCTGGGACACCCTGGCCAGCCTGGGCCTG<br>CTGAACAAGCACGCCAAGCTGCTGTTCCTGGGCCTGGACAACGCCGGC<br>AAGACCACCCTGCTGCACATGCTGAAGAACGACAGGGTGGCCATCCTG<br>CAGCCCACCCTGCACCCCACCAGCGAGGAGCTGGCCATCGGCAACGT<br>GAAGTTCACCACCTTCGACCTGGGCGGCCACCAGCAGGCCAGGAGGC<br>TGTGGAAGGACTACTTCCCCGAGGTGAGCGGCATCGTGTTCCTGGTGG<br>ACAGCAAGGACCACGAGAGGTTCATCGAGAGCAAGGCCGAGCTGGAC<br>GCCCTGCTGAGCATGGAGGACCTGAGCAAGGTGCCCTTCCTGATCCTG<br>GGCAACAAGATCGACCACCCCGACGCCATCAGCGAGGACCAGCTGAG<br>GCACGAGCTGGGCCTGTACCAGACCACCGGCAAGGGCAAGGTGCCCC<br>TGGAGGGCATCAGGCCCATCGAGGTGTTCATGTGCAGCGTGGTGATGA<br>GGCAGGGCTACGGCGAGGGCATCAGGTGGCTGAGCCAGTACGTGGGT<br>TCAGGAATGGTGAGCAAGGGCGAGGAGCTGTTCACCGGGGTGGTGCC<br>CATCCTGGTCGAGCTGGACGGCGACGTAAACGGCCACAAGTTCAGCGT |

| | |
|---|---|
| | GTCCGGCGAGGGCGAGGGCGATGCCACCTACGGCAAGCTGACCCTGA<br>AGTTCATCTGCACCACCGGCAAGCTGCCCGTGCCCTGGCCCACCCTCG<br>TGACCACCCTGACCTACGGCGTGCAGTGCTTCAGCCGCTACCCCGACC<br>ACATGAAGCAGCACGACTTCTTCAAGTCCGCCATGCCCGAAGGCTACGT<br>CCAGGAGCGCACCATCTTCTTCAAGGACGACGGCAACTACAAGACCCG<br>CGCCGAGGTGAAGTTCGAGGGCGACACCCTGGTGAACCGCATCGAGC<br>TGAAGGGCATCGACTTCAAGGAGGACGGCAACATCCTGGGGCACAAGC<br>TGGAGTACAACTACAACAGCCACAACGTCTATATCATGGCCGACAAGCA<br>GAAGAACGGCATCAAGGTGAACTTCAAGATCCGCCACAACATCGAGGAC<br>GGCAGCGTGCAGCTCGCCGACCACTACCAGCAGAACACCCCCATCGGC<br>GACGGCCCCGTGCTGCTGCCCGACAACCACTACCTGAGCACCCAGTCC<br>GCCCTGAGCAAAGACCCCAACGAGAAGCGCGATCACATGGTCCTGCTG<br>GAGTTCGTGACCGCCGCCGGGATCACTCTCGGCATGGACGAGCTGTAC<br>AAGTACGTTGGCGGCTCTTCTGACTACAAGGATGACGATGACAAGTGA |
| **V64C-*Bc*SAR1-eGFP** | ATGTGGTTACTTGATTGGTTTTGGGATACGTTAGCATCCTTGGGTCTCTT<br>GAATAAACATGCAAAATTACTCTTCTTGGGATTGGATAATGCTGGGAAAAC<br>TACATTGCTCCATATGCTCAAGAATGATAGAGTAGCTATTCTTCAACCAAC<br>TCTCCATCCAACGAGTGAAGAACTTGCTATTGGGAATTGTAAATTTACGA<br>CATTTGATCTTGGTGGGCATCAACAAGCGAGACGTCTCTGGAAAGATTAT<br>TTTCCGGAAGTCTCAGGGATTGTCTTTCTCGTTGATTCCAAAGATCATGA<br>ACGTTTTATTGAATCCAAAGCAGAACTCGATGCGCTTTTATCCATGGAAG<br>ATTTATCAAAAGTCCCATTCCTCATTTTGGGGAATAAAATTGATCATCCAG<br>ATGCTATTAGTGAAGATCAATTACGACATGAACTTGGGCTTTATCAAACTA<br>CGGGAAAAGGTAAAGTCCCATTGGAAGGAATACGGCCAATTGAAGTATTT<br>ATGTGTTCTGTCGTCATGCGCCAAGGATATGGAGAAGGTATACGCTGGTT<br>GTCCCAATATGTCGGCTCTGGCATGGTCTCAAAAGGTGAAGAACTGTTT<br>ACTGGTGTCGTCCCAATTCTTGTGGAATTAGATGGGGATGTGAATGGAC<br>ATAAATTTTCTGTATCTGGTGAAGGGGAAGGAGACGCTACGTATGGGAAA<br>TTGACTCTCAAATTTATTTGTACGACTGGTAAATTACCTGTCCCTTGGCCT<br>ACACTTGTAACGACATTGACATATGGGGTCCAATGTTTTAGCAGGTATCC<br>GGATCATATGAAACAACATGATTTCTTCAAATCAGCAATGCCAGAGGGAT<br>ATGTTCAAGAAAGAACAATTTTCTTTAAAGACGACGGTAACTATAAAACGA<br>GGGCCGAAGTCAAATTTGAGGGAGATACTTTAGTTAATAGAATAGAATTAA<br>AAGGGATAGATTTTAAAGAAGATGGGAATATTCTGGGACATAAGTTAGAAT<br>ATAATTATAACTCCCACAATGTTTACATTATGGCAGATAAACAGAAGAATG<br>GAATTAAAGTTAATTTCAAAATTCGTCATAATATTGAAGATGGTTCCGTCCA<br>GTTGGCAGATCATTATCAACAAAATACACCAATTGGGGATGGACCTGTTC<br>TCCTTCCTGATAATCATTATCTCTCCACTCAAAGTGCACTCAGTAAGGATC<br>CAAATGAAAAGCGGGACCATATGGTTCTTTTAGAATTTGTTACGGCAGCA<br>GGTATTACCCTGGGTATGGATGAACTTTATAAATATGTCGGCGGGAGTAG<br>CGATTATAAAGACGATGACGATAAATAA |
| **WT-*Pg*SAR1-eGFP** | ATGTTTATTCTCAATTGGTTTTGGGATGTTCTTGCAAATCTTGGATTAGTTA<br>ATAAGAATGCGAAAATTCTCTTTCTCGGTCTTGATAATGCTGGCAAAACAA<br>CACTTCTCCATATGCTCAAGAATGATCGGCTCGCTACTCTCCAACCAACT<br>CTTCATCCTACTTCCGAAGAATTGGCAATAGGTAATGTTAAATTTACGACT<br>TATGATTTAGGTGGACATCAACAAGCGCGCAGACTTTGGAAAGAATATTT<br>TCCAGAAGTAAATGGTATCGTATTTCTCGTTGATGCTCAAGATCCTGAAC<br>GGTTTTCAGAAAGCAAAATTGAGTTAGATGCATTATTGTCAATTGAAGAAC<br>TTTCCAAAGTCCCATTTCTCATATTGGGTAATAAAATCGATGCTCCAGGGG<br>CTGTTAGTGAAGAAGATCTCAGACATTGTCTCGGACTCTATCAAACTACA<br>GGAAAAGGAAAAGTCCCGCTTATTGATATTCGGCCAATTGAAGTATTTATG |

| | |
|---|---|
| | TGTTCCATTGTTATGCGTCAAGGTTATGGTGATGGGTTTCGATGGCTCGC<br>TCAATATATAGGCTCTGGCATGGTCTCAAAAGGAGAAGAACTGTTTACAG<br>GAGTTGTACCTATTCTCGTGGAACTCGATGGGGATGTGAATGGACATAAA<br>TTTAGTGTTAGCGGAGAAGGAGAAGGTGACGCTACATATGGGAAACTTA<br>CACTTAAATTTATTTGTACAACAGGGAAACTCCCAGTCCCGTGGCCAACT<br>CTGGTAACTACACTTACATATGGGGTTCAATGCTTTTCCCGGTATCCTGAT<br>CATATGAAACAACATGATTTCTTTAAATCTGCGATGCCTGAGGGATATGTG<br>CAAGAACGTACTATTTTCTTCAAGGATGACGGGAATTATAAAACTCGAGC<br>CGAAGTAAAATTTGAAGGAGATACTCTTGTCAATCGTATTGAACTTAAAGG<br>GATTGATTTTAAAGAGGACGGGAACATTTTGGGCCACAAACTTGAATACA<br>ATTACAACTCACATAATGTTTACATTATGGCTGATAAACAAAAGAACGGTAT<br>TAAGGTTAATTTTAAAATTAGGCATAATATTGAAGATGGGAGTGTACAATTG<br>GCCGACCATTATCAACAAAATACACCTATTGGGGATGGACCTGTACTCTT<br>ACCTGATAATCATTATCTCTCTACACAAAGCGCTCTTTCCAAGGATCCAAA<br>TGAAAAGCGAGACCATATGGTGCTTTTGGAATTTGTCACTGCAGCAGGC<br>ATTACCCTGGGAATGGATGAATTGTATAAATATGTGGGTGGTAGCAGCGA<br>TTATAAAGACGATGACGATAAATAG |
| **V64C-*Pg*SAR1-eGFP** | ATGTTTATACTCAATTGGTTTTGGGATGTTCTTGCTAATCTCGGTCTTGTC<br>AATAAGAATGCTAAAATATTATTTCTTGGATTGGATAATGCAGGAAAGACA<br>ACTCTCCTTCATATGTTGAAGAATGATCGACTTGCCACATTACAACCAAC<br>GTTACATCCTACTTCAGAAGAATTAGCAATAGGAAATTGTAAATTTACAAC<br>GTATGATTTGGGCGGTCATCAACAAGCACGAAGATTGTGGAAAGAATATT<br>TTCCTGAAGTTAATGGAATTGTATTTCTTGTAGATGCACAAGATCCTGAGA<br>GATTTTCCGAATCAAAGATAGAACTCGATGCACTCCTCAGTATTGAAGAA<br>CTTTCCAAAGTCCCTTTTCTGATTCTGGGAAATAAAATTGATGCACCAGG<br>AGCAGTTTCAGAAGAAGATCTTCGACATTGTTTGGGGTTGTATCAAACAA<br>CTGGTAAAGGAAAAGTACCTCTCATTGATATTCGACCTATTGAAGTTTTCA<br>TGTGTTCAATTGTTATGCGCCAAGGGTATGGTGATGGGTTTCGCTGGTTG<br>GCGCAATATATTGGCAGCGGCATGGTATCCAAAGGAGAAGAACTCTTTAC<br>AGGCGTTGTCCCAATTCTTGTGGAATTAGATGGGGATGTCAATGGTCATA<br>AATTTTCCGTCAGCGGGGAAGGTGAAGGTGACGCGACGTATGGGAAAC<br>TCACTCTCAAATTTATTTGTACGACGGGAAAATTGCCAGTCCCTTGGCCG<br>ACACTGGTCACTACATTGACATATGGAGTCCAATGTTTTAGTAGATATCCA<br>GATCATATGAAACAACATGATTTCTTTAAAAGCGCAATGCCAGAGGGATAT<br>GTGCAAGAACGGACTATTTTCTTTAAAGATGACGGAAATTATAAAACGAG<br>GGCTGAAGTAAAGTTTGAAGGGGACACATTAGTAAATAGAATTGAGCTCA<br>AAGGGATTGATTTTAAGGAAGATGGGAATATTTTGGGACATAAACTCGAAT<br>ATAATTATAATTCCCATAATGTATACATTATGGCGGATAAACAAAAGAATGG<br>GATTAAAGTCAATTTTAAAATTAGGCATAATATCGAAGATGGTTCCGTCCA<br>ACTGGCTGATCATTATCAACAAAATACACCTATTGGAGATGGTCCAGTACT<br>CCTGCCTGATAATCATTATCTCTCAACACAAAGCGCGCTTAGTAAGGATC<br>CGAATGAGAAACGTGACCATATGGTACTCCTCGAATTTGTCACAGCTGCT<br>GGAATTACCCTGGGAATGGATGAACTCTATAAATATGTCGGCGGTAGCTC<br>AGATTATAAAGACGATGACGATAAATAA |
| **WT-*Hs*SAR1A-eGFP** | ATGTCATTTATCTTTGAATGGATTTATAATGGTTTCAGTTCCGTCTTGCAAT<br>TTCTTGGTCTTTATAAGAAAAGCGGAAAATTAGTTTTCCTTGGACTCGATA<br>ACGCTGGCAAAACAACATTGCTCCATATGCTCAAAGATGATCGGTTAGGG<br>CAACATGTCCCTACTCTCCATCCTACGTCAGAAGAATTAACTATAGCAGG<br>AATGACTTTTACTACGTTTGATTTGGGTGGGCATGAACAAGCTAGACGAG<br>TATGGAAGAATTATCTTCCTGCTATTAATGGGATAGTATTTCTCGTCGATTG<br>TGCAGATCATTCCCGCCTTGTTGAATCCAAAGTCGAATTGAATGCACTCA |

| | |
|---|---|
| | TGACAGATGAAACGATTAGTAATGTACCGATACTCATACTTGGGAATAAAA<br>TTGATCGCACAGATGCAATTTCTGAAGAAAAGTTACGAGAAATCTTTGGC<br>CTTTATGGACAAACAACTGGTAAAGGGAATGTCACGTTAAAAGAACTTAA<br>TGCACGCCCTATGGAAGTCTTTATGTGTTCCGTTTTGAAACGACAAGGAT<br>ATGGAGAAGGTTTTAGATGGCTTTCCCAATATATTGATGGAAGCGGTATG<br>GTTTCAAAAGGAGAAGAACTCTTTACGGGCGTCGTTCCTATACTTGTGGA<br>ACTCGATGGGGATGTGAATGGTCATAAATTTTCTGTCAGCGGAGAAGGA<br>GAAGGGGACGCTACATATGGTAAATTGACTCTTAAATTTATTTGTACAACA<br>GGGAAATTACCTGTCCCATGGCCTACACTGGTCACAACTTTAACTTATGG<br>TGTCCAATGTTTCTCCAGATATCCAGATCATATGAAACAACATGATTTCTTT<br>AAATCTGCGATGCCTGAGGGTTACGTACAAGAACGGACAATATTCTTTAA<br>AGATGATGGTAATTATAAAACACGGGCGGAAGTCAAATTTGAGGGAGATA<br>CCCTTGTCAACAGAATCGAATTGAAAGGGATAGATTTTAAAGAGGATGGT<br>AATATATTGGGTCATAAACTCGAGTATAATTATAATTCCCATAATGTGTACAT<br>TATGGCTGATAAACAGAAGAATGGAATTAAAGTTAATTTTAAAATTAGGCA<br>CAATATTGAAGATGGGTCAGTTCAGCTTGCAGATCATTATCAACAAAATAC<br>ACCAATTGGGGATGGGCCTGTTCTGCTCCCGGATAATCATTATCTCTCCA<br>CACAAAGCGCTCTTTCCAAGGATCCTAATGAGAAAAGGGACCATATGGTA<br>CTTTTGGAATTTGTCACAGCTGCTGGCATAACCCTTGGGATGGATGAACT<br>TTATAAATATGTGGGCGGAAGCTCCGATTATAAAGACGATGACGATAAATA<br>G |
| **M69C-*Hs*SAR1A-eGFP** | ATGTCATTTATTTTCGAATGGATATATAATGGATTTAGTTCTGTCCTTCAATT<br>TCTCGGGCTCTATAAGAAATCAGGGAAATTAGTCTTCTTGGGGTTGGATA<br>ATGCAGGTAAAACAACACTCTTGCATATGCTTAAAGATGATAGATTGGGG<br>CAACATGTTCCTACACTCCATCCAACTTCTGAAGAACTCACTATTGCTGG<br>TTGCACTTTTACAACGTTTGATCTTGGCGGACATGAACAAGCACGTCGA<br>GTATGGAAGAATTATTTGCCGGCGATTAATGGAATTGTCTTTCTTGTCGAT<br>TGTGCTGATCATTCTAGATTGGTCGAAAGTAAAGTCGAACTCAATGCTTT<br>GATGACTGATGAAACGATATCTAATGTTCCAATTCTCATTCTCGGAAATAA<br>AATTGATCGCACTGATGCGATTTCCGAAGAGAAACTCAGAGAAATTTTCG<br>GGCTCTATGGGCAAACGACTGGGAAAGGGAATGTTACACTTAAAGAATTA<br>AATGCTCGGCCAATGGAAGTATTTATGTGTTCAGTCCTCAAACGGCAAGG<br>ATATGGAGAAGGGTTTCGGTGGCTTAGTCAATATATTGATGGGAGTGGGA<br>TGGTATCCAAAGGAGAAGAACTTTTCACGGGTGTTGTCCCAATATTGGTT<br>GAACTCGATGGTGATGTTAATGGACATAAATTTTCCGTATCTGGTGAAGG<br>AGAAGGAGACGCAACATATGGAAAATTGACTCTTAAATTTATTTGTACTAC<br>TGGGAAATTGCCTGTACCGTGGCCTACATTGGTTACAACATTAACTTATG<br>GAGTCCAATGTTTTAGTAGGTATCCGGATCACATGAAACAACATGATTTCT<br>TTAAAAGTGCGATGCCAGAGGGATACGTGCAAGAACGAACAATATTCTTT<br>AAAGATGATGGAAATTACAAAACTCGGGCCGAGGTTAAATTTGAAGGAGA<br>TACTCTTGTTAATAGAATTGAGTTGAAAGGGATAGATTTCAAAGAAGATGG<br>GAATATACTCGGCCATAAACTCGAATATAATTATAATTCTCATAATGTGTACA<br>TAATGGCTGATAAACAAAAGAATGGAATTAAAGTTAATTTTAAAATCCGAC<br>ATAATATTGAAGATGGAAGTGTACAACTGGCTGATCATTACCAACAAAATA<br>CTCCGATAGGGGATGGTCCTGTCTTGTTACCAGATAATCATTATTTGTCCA<br>CTCAAAGCGCTTTGAGTAAGGATCCGAATGAAAAGCGGGACCATATGGT<br>ATTGCTCGAATTTGTTACAGCAGCTGGCATTACGCTGGGAATGGATGAAC<br>TCTATAAATATGTGGGCGGGTCCAGCGATTATAAAGACGATGACGATAAAT<br>AA |

**Supplementary Table 2.** Antibody product identifiers and dilutions used for this study.

| Antibody | Dilution factor | Catalog number | RRID |
|---|---|---|---|
| **α-FLAG-HRP** | 1:2000 | Cell Signaling Technology; #86861 | AB_2800094 |
| **α-GAPDH-HRP** | 1:2000 | Cell Signaling Technology; #3683 | AB_1642205 |
| **α-GSPT1 (α-eRF3)** | 1:2000 | Cell Signaling Technology; #14980 | AB_2798677 |
| **α-Luciferase-HPR** | 1:2000 | Rockland; #200-103-150-0100 | AB_2610914 |
| **α-$Na^+$/$K^+$-ATPase-HRP** | 1:2000 | Cell Signaling Technology; #96124 | AB_2800256 |
| **α-β-actin-HRP** | 1:2000 | Cell Signaling Technology; #5125 | AB_1903890 |
| **α-V5** | 1:2000 | Cell Signaling Technology; #13202 | AB_2687461 |
| **Goat anti-rabbit-HRP** | 1:5000 | Thermo Fisher Scientific; #31460 | AB_228341 |

**Supplementary Synthetic Chemistry Information**

**Synthetic Chemistry Procedures and Analytical Data**

All chemical reagents were purchased from commercial suppliers and used without further purification unless otherwise noted. WX-01-06 and WX-01-08 were prepared as previously reported (>95% ee)[1].

Routine liquid chromatography-mass spectrometry (LCMS) analysis was performed on a Waters I-Class LC and a Waters Acquity QDA MS with a Waters Cortecs C18 column (1.6 um, 2.1 x 55 mm) using a 5–99% B gradient lasting 2.5 minutes (A: 0.1% aqueous formic acid, B: 0.06% formic acid in acetonitrile, 0.8 mL/min flow rate, 35 °C column temperature). The product purity was quantified on a combined 220 nm + 260 nm UV channel and the product identity was verified by mass spectrometry. Final compounds were confirmed to be greater than 95% pure by LCMS (UV) prior to use.

Preparative high-pressure liquid chromatography (prep-HPLC) was performed on a Waters Autopurification LC with a Waters BEH C18 column (5 um, 19 x 160 mm) and/or Teledyne ACCQPrep HP150 LC with a RediSep® Prep C18 Column (20 mm x 150 mm).

NMR spectra were recorded at room temperature on a Bruker AVIII HD 600 MHz NMR equipped with 5 mm CPDCH CryoProbe. Spectra are reported as follows: chemical shift (δ, ppm relative to residual solvent of 3.50 ppm), apparent multiplicity (s = singlet, br. s = broad singlet, d = doublet, t = triplet, q = quartet, p = hextet, m = multiplet, or a combination thereof), and coupling constant (*J*, Hz).

Mass measurements for high-resolution mass spectrometry (HRMS) were performed on a Waters Xevo G2-XS TOF calibrated against sodium formate clusters and using a LeuEnk lockmass. Expected monoisotopic masses were calculated using MassLynx 4.1 and the m/z values for calibrant and lockmass were MassLynx-default values.

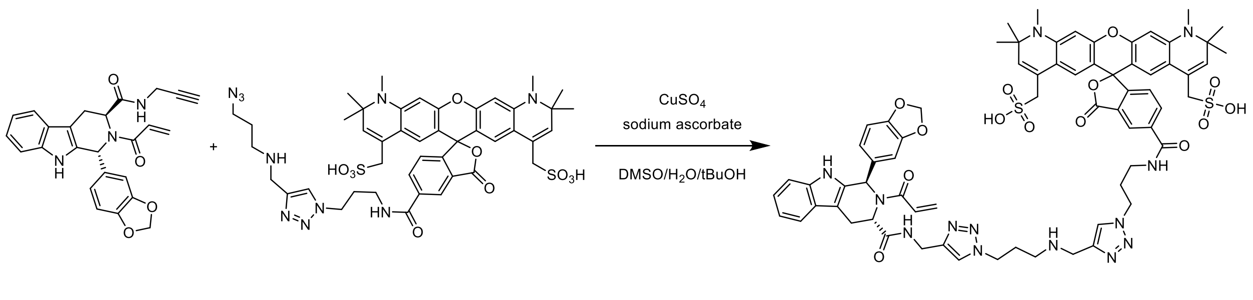


WX-01-06 AZDye 594 plus CJR-88A

**Scheme 1:** Synthesis of CJR-88A from WX-01-06 and AZDye 594 plus.

(5-((3-(4-(((3-(4-(((1*R*,3*S*)-2-acryloyl-1-(benzo[d][1,3]dioxol-5-yl)-2,3,4,9-tetrahydro-1H-pyrido[3,4-b]indole-3-carboxamido)methyl)-1H-1,2,3-triazol-1-yl)propyl)amino)methyl)-1H-1,2,3-triazol-1-yl)propyl)carbamoyl)-1',2',2',10',10',11'-hexamethyl-3-oxo-1',2',10',11'-tetrahydro-3H-spiro[isobenzofuran-1,6'-pyrano[3,2-g:5,6-g']diquinoline]-4',8'-diyl)dimethanesulfonic acid **(CJR-88A).**

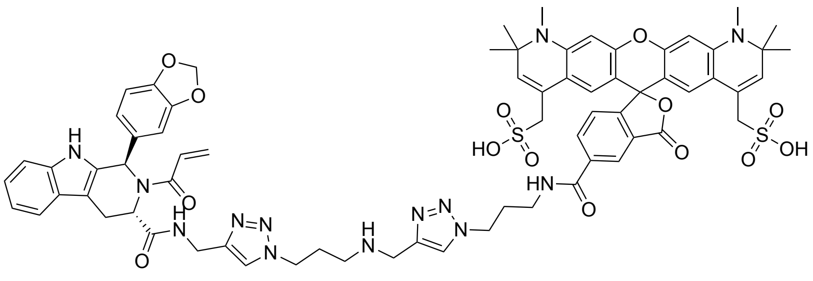

A 2-dram vial was charged with WX-01-06[1] (0.243 µL of a 50 mM DMSO solution, 5.19 mg, 12.1 µmol, 1 equiv.), copper sulfate (24.3 µL of a 50 mM aq. solution, 0.194 mg, 1.21 µmol, 0.1 equiv.), AZDye 594 Azide Plus (4.08 mL of a 3.125 mM solution in 50% DMSO/water v/v, 12.0 mg, 12.7 µmol, 1.05 equiv., Vector Laboratories CCT-1481) and degassed *tert*-butanol (1.21 mL). The reaction was placed under a nitrogen atmosphere and then treated with a degassed, aq. solution of sodium ascorbate (0.36 mL of a 50 mM solution, 3.61 mg, 18.2 µmol, 1.5 equiv.). The solution was stirred at room temperature for 2 h and monitored by LCMS. The crude reaction was stored at -80°C prior to purification. The product was purified by reverse-phase preparatory HPLC in water/MeCN containing 0.1% formic acid (v/v). Pure fractions were pooled and concentrated by lyophilization to afford a blue-purple solid (9.39 mg, 6.85 µmol, 56% yield).

**$^1$H NMR** (600 MHz, DMSO-$d_6$ with minimal $D_2O$, mixture of rotamers, open/closed forms) δ 11.28 – 10.79 (m, 1H), 9.13 – 8.84 (m, 1H), 8.60 – 8.38 (m, 2H), 8.23 (s, 1H), 8.19 – 8.11 (m, 1H), 7.44 – 7.23 (m, 3H), 7.13 – 6.5 (m, 10H), 6.36 – 5.97 (m, 1H), 5.96 – 5.85 (m, 2H), 5.83 – 5.24 (m, 3H), 4.52 (t, *J* = 6.8 Hz, 2H), 4.27 – 4.03 (m, 4H), 3.98 – 3.86 (m, 2H), 3.50 (s, 6H, overlaps with water), 3.37 (q, *J* = 6.3 Hz, 2H), 3.31 – 3.14 (m, 5H), 3.03 (br. s, 6H), 2.86 (t, *J* = 7.8 Hz, 2H), 2.19 (p, *J* = 6.7 Hz, 2H), 1.90 (p, *J* = 7.3 Hz, 2H), 1.38 (s, 12H). 1 exchangeable proton not observed.

**HRMS** ESI-TOF m/z calculated for $C_{69}H_{72}N_{13}O_{14}S_2$ $[M+H]^+$ 1370.4763. Found 1370.4738.

**LCMS** m/z calculated for $C_{69}H_{73}N_{13}O_{14}S_2$ $[M+2H]^{2+}$ 685.7. Found. 686.4 Retention time: 1.15 min.

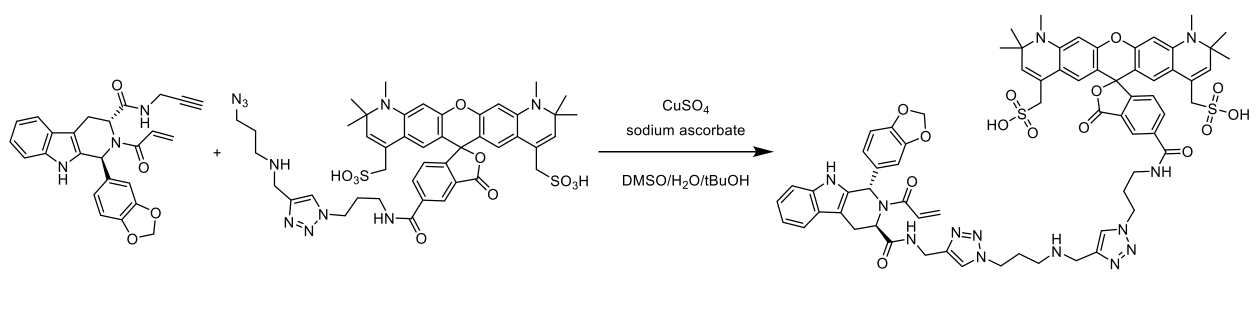


WX-01-08 AZDye 594 plus CJR-88B

**Scheme 2:** Synthesis of CJR-88B from WX-01-08 and AZDye 594 plus.

(5-((3-(4-(((3-(4-(((1*S*,3*R*)-2-acryloyl-1-(benzo[d][1,3]dioxol-5-yl)-2,3,4,9-tetrahydro-1H-pyrido[3,4-b]indole-3-carboxamido)methyl)-1H-1,2,3-triazol-1-yl)propyl)amino)methyl)-1H-1,2,3-triazol-1-yl)propyl)carbamoyl)-1',2',2',10',10',11'-hexamethyl-3-oxo-1',2',10',11'-tetrahydro-3H-spiro[isobenzofuran-1,6'-pyrano[3,2-g:5,6-g']diquinoline]-4',8'-diyl)dimethanesulfonic acid **(CJR-88B).**

Prepared from WX-01-08[1] in an analogous fashion to CJR-88A to afford a blue-purple solid (8.35 mg, 6.09 µmol, 51% yield).

**$^1$H NMR** (600 MHz, DMSO with minimal $D_2O$, mixture of rotamers, open/closed forms) δ 11.28 – 10.90 (m, 1H), 9.13 – 8.84 (m, 1H), 8.65 – 8.37 (m, 2H), 8.24 (s, 1H), 8.20 – 8.10 (m, 1H), 7.47 – 7.24 (m, 3H), 7.13 – 6.48 (m, 10H), 6.35 – 5.97 (m, 2H), 5.96 – 5.86 (m, 2H), 5.80 – 5.21 (m, 3H), 4.52 (t, *J* = 6.8 Hz, 2H), 4.28 – 4.03 (m, 4H), 3.99 – 3.86 (m, 2H), 3.45 (s, 6H, overlaps with water), 3.37 (q, *J* = 6.3 Hz, 2H), 3.31 – 3.13 (m, 5H), 3.01 (br. s, 6H), 2.87 (t, *J* = 7.8 Hz, 2H), 2.18 (p, *J* = 6.7 Hz, 2H), 1.90 (p, *J* = 7.1 Hz, 2H), 1.37 (s, 12H).

**HRMS** ESI-TOF m/z calculated for $C_{69}H_{72}N_{13}O_{14}S_2$ $[M+H]^+$ 1370.4763. Found 1370.4717.

**LCMS** m/z calculated for $C_{69}H_{73}N_{13}O_{14}S_2$ $[M+2H]^{2+}$ 685.7. Found. 686.3 Retention time: 1.15 min.

$^{1}H$ NMR spectrum of CJR-88A (600 MHz, DMSO-$d_6$ with minimal $D_2O$)

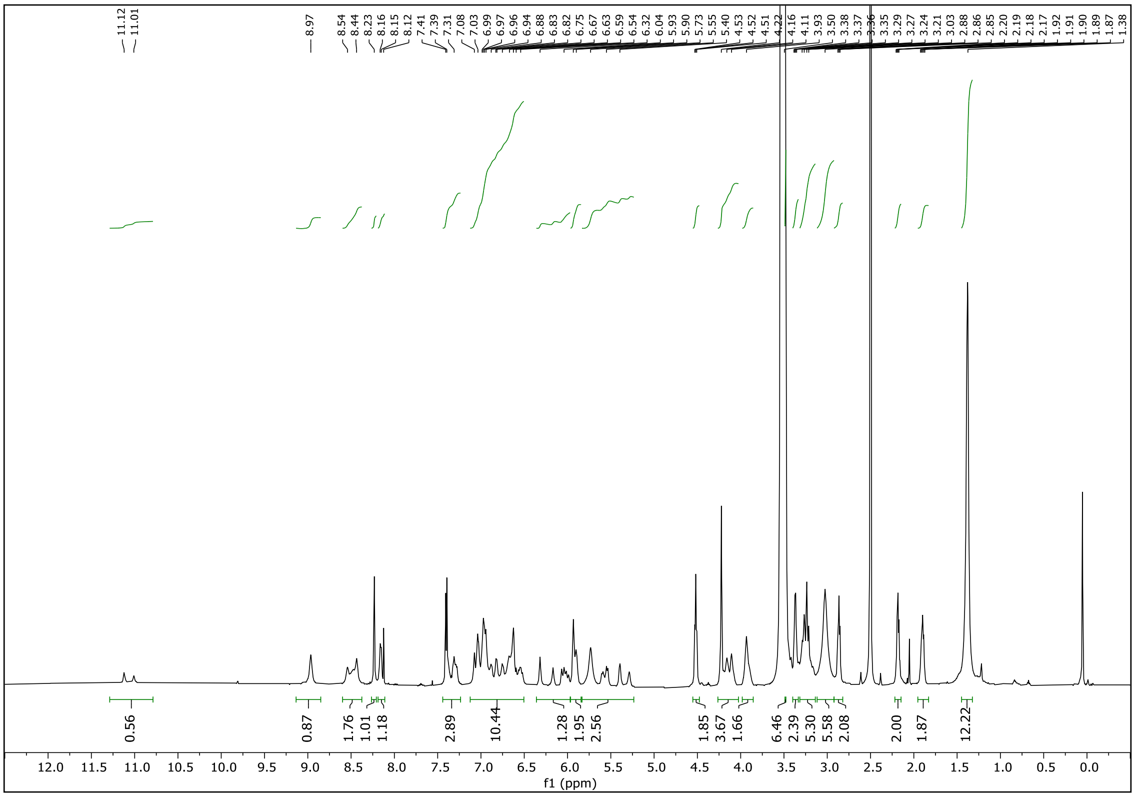

$^{1}H$ NMR spectrum of CJR-88B (600 MHz, DMSO-$d_6$ with minimal $D_2O$)

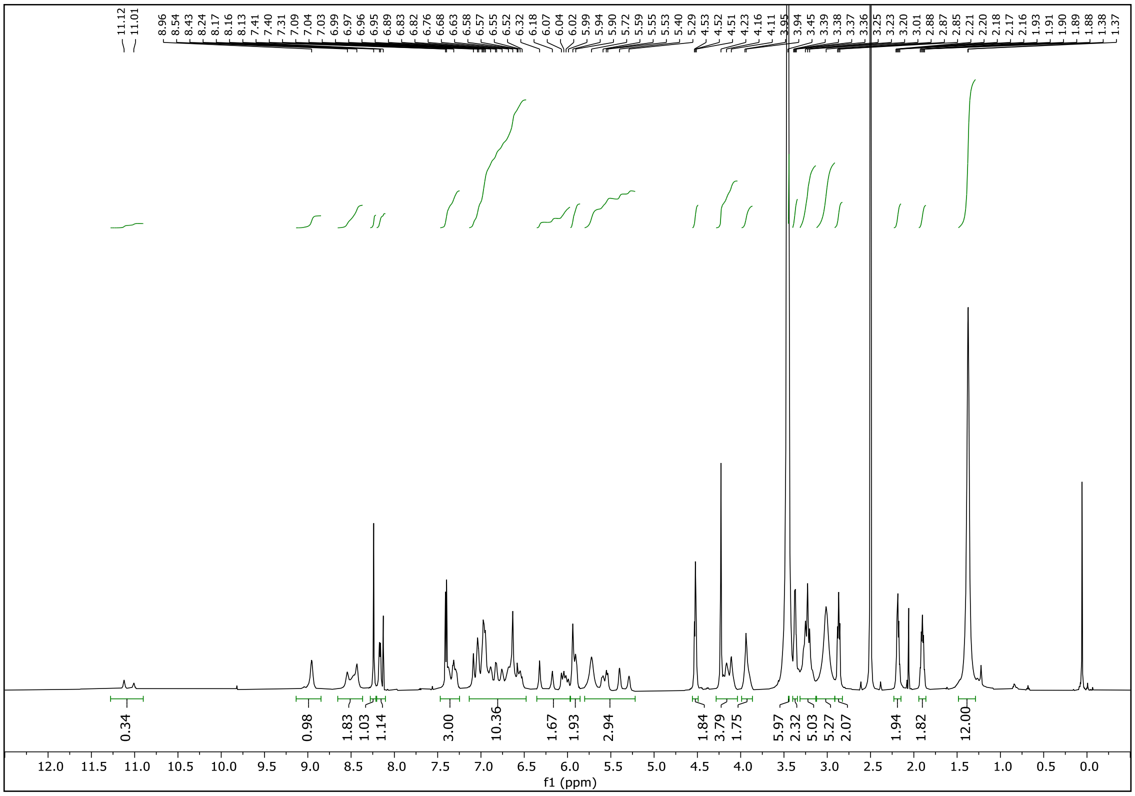
